# Inferring the direction of introgression using genomic sequence data

**DOI:** 10.1101/2023.06.16.545313

**Authors:** Yuttapong Thawornwattana, Jun Huang, Tomáš Flouri, James Mallet, Ziheng Yang

## Abstract

Genomic data are informative about the history of species divergence and interspecific gene flow, including the direction, timing, and strength of gene flow. However, gene flow in opposite directions generates similar patterns in multilocus sequence data, such as reduced sequence divergence between the hybridizing species. As a result, inference of the direction of gene flow is challenging. Here we investigate the information about the direction of gene flow present in genomic sequence data using likelihood-based methods under the multispecies-coalescent-with-introgression (MSci) model. We analyze the case of two species, and use simulation to examine cases with three or four species. We find that it is easier to infer gene flow from a small population to a large one than in the opposite direction, and easier to infer inflow (gene flow from outgroup species to an ingroup species) than outflow (gene flow from an ingroup species to an outgroup species). It is also easier to infer gene flow if there is a longer time of separate evolution between the initial divergence and subsequent introgression. When introgression is assumed to occur in the wrong direction, the time of introgression tends to be correctly estimated and the Bayesian test of gene flow is often significant, while estimates of introgression probability can be even greater than the true probability. We analyze genomic sequences from *Heliconius* butterflies to demonstrate that typical genomic datasets are informative about the direction of interspecific gene flow, as well as its timing and strength.

## Introduction

Gene flow between species occurs as a result of hybridization followed by backcrossing in one of the hybridizing species. While it has a predominantly homogenizing effect, interspecific gene flow may create new beneficial combinations of alleles at multiple loci, facilitating species diversification and adaptation (Arnold and Kunte, 2017; Campbell *et al*., 2018; Feurtey and Stukenbrock, 2018; Marques *et al*., 2019; Edelman and Mallet, 2021). The outcome of introgression in each direction is influenced by multiple factors including mate choice (Peters *et al*., 2017), ecological selection, and hybrid incompatibility (for reviews, see Coyne and Orr, 2004; Martin and Jiggins, 2017; Moran *et al*., 2021). Given that these factors typically differ between species and that selection on introgressed material acts independently in different recipient species, it is likely that gene flow is often asymmetrical, being more prevalent in one direction than in the other. Reliable inference of the direction of introgression, as well as its timing and rate, will advance our understanding of this important evolutionary process and its consequences, including the role of gene flow during speciation and the adaptive nature of introgressed alleles.

Two models of interspecific gene flow have been developed in the multispecies coalescent (MSC) framework, representing different modes of gene flow (Jiao *et al*., 2021; Hibbins and Hahn, 2022). The MSC-with-introgression (MSci; Flouri *et al*. 2020) model, also known as multispecies network coalescent (MSNC, Yu *et al*., 2012; Wen and Nakhleh, 2018; Zhang *et al*., 2018), assumes that gene flow occurs at a particular time point in the past. The magnitude of gene flow is measured by the introgression probability (*φ*), the proportion of immigrants in the recipient population at the time of introgression. The MSC-with-migration (MSC-M) model, also known as the isolation-with-migration (IM) model, assumes that gene flow occurs continuously at a certain rate every generation after species divergence (Nielsen and Wakeley, 2001; Hey *et al*., 2018). The rate of gene flow is measured by the expected number of immigrants from populations *A* to *B* per generation, *M*_*AB*_ = *N*_*B*_*m*_*AB*_, where *N*_*B*_ is the (effective) population size of population *B* and *m*_*AB*_ is the proportion of immigrants in population *B* from *A*. In both models, the rates of gene flow (*φ* or *M*) are ‘effective’ rates, reflecting combined effects of gene flow and negative or positive natural selection on introgressed alleles, influenced by the local recombination rate (Petry, 1983; Barton and Bengtsson, 1986).

Interspecific gene flow alters gene genealogies, causing fluctuations over the genome in the genealogical history of sequences sampled from extant species. Under both the MSC-M and MSci models, gene trees and coalescent times have probabilistic distributions specified by the model and parameters, including species divergence times, population sizes for extant and extinct species, and the rate of gene flow (see Yang, 2014; Jiao *et al*., 2021 for reviews). Multilocus sequence alignments are informative about gene tree topologies and coalescent times, and thus about the direction of gene flow as well as its timing and strength. However, opposite directions of gene flow often create similar features in gene genealogies and in the sequence data. For example, gene flow in either direction reduces the average and minimum divergence between the hybridizing species. In the special case of sampling one sequence per species per locus, the data cannot identify introgression direction between two sister species (say *A* and *B*), because the coalescent time (*t*_*ab*_) between the two sequences at each locus (*a, b*) has the same distribution under the models with *A* → *B* or *B* → *A* introgression (Yang and Flouri, 2022, fig. 10; see also Discussion). If multiple sequences are sampled per species per locus, introgression direction becomes identifiable (Yang and Flouri, 2022). Even so, inference of introgression direction may be expected to be a challenging task. This is particularly so for heuristic methods for inferring gene flow based on summary statistics. For example, the *D* statistic (Green *et al*., 2010; Durand *et al*., 2011) operates on species quartets and cannot identify the direction of gene flow. Although heuristic methods exist for inferring the direction of gene flow, based on estimated local genomic divergences (Green *et al*., 2010, fig. S39) or genome-wide site-pattern counts (*D*_FOIL_, Pease and Hahn, 2015), they do not make an efficient use of information in the data, often require a specific species phylogeny and sampling setup, and cannot infer gene flow between sister lineages. For recent discussions of the strengths and weaknesses of heuristic versus likelihood methods, see Jiao *et al*. (2021), Hibbins and Hahn (2022), Huang *et al*. (2022), and Yang and Flouri (2022).

Here, we study the inference of introgression direction, focusing on the Bayesian method under the MSci model (Flouri *et al*., 2020). Suppose introgression occurs from species *A* → *B* but we analyze genomic data assuming *B* → *A* introgression. We address the following questions. (a) Will we often detect introgression despite the assumed wrong direction? (b) How will the estimated introgression probability 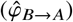 compare with the true introgression probability (*φ*_*A*→*B*_)? (c) How reliable will estimates of the time of introgression be, as well as other parameters such as species divergence times and population sizes? (d) Does the method behavior differ depending on whether gene flow is between sister lineages or between non-sister lineages, and whether gene flow is from a small population to a large one, or in the opposite direction? (e) How can we infer the direction of introgression (*A* → *B* vs. *B* → *A*)? (f) Are typical genomic data informative about the direction of gene flow? We focus on both Bayesian estimation of parameters, in particular, the introgression probability (Flouri *et al*., 2020) and Bayesian testing of introgression (Ji *et al*., 2023).

We use a combination of mathematical analysis and computer simulation to characterize features of sequence data that are informative about the direction of gene flow. We first study the case of two species (*A, B*) by examining the distribution of coalescent times (*t*_*aa*_, *t*_*ab*_, *t*_*bb*_) under the MSci model. The theory allows us to compare and quantify the amount of information in the data under different scenarios. Next, we explore the amount of information gained when a third species is added to branches of the species tree for two species and study the impact of introgression direction when gene flow involves non-sister species. Finally, we test these methods with genomic sequences from three species of *Heliconius* butterflies to verify the applicability of our results derived from the theoretical analysis and computer simulation and to demonstrate how the framework can be applied to infer the direction of gene flow, as well as its timing and strength. Our results provide practical guidelines for inferring introgression and its direction from genomic sequence data.

## Results

### Notation and problem setup

We use the MSci model of figure 1**a** with *A* → *B* introgression to introduce the notation and set up the problem. Species *A* and *B* diverged at time τ_*R*_ and hybridized later at time τ_*X*_. The magnitude of introgression is measured by the introgression probability or admixture proportion *φ*_*Y*_, which is the proportion of immigrants in population *B* from *A* at the time of introgression. There are three types of parameters in the model: species divergence times or introgression times (τ_*R*_, τ_*X*_), population sizes for extant and extinct species (*θ*_*A*_, *θ*_*B*_, *θ*_*X*_, *θ*_*Y*_, *θ*_*R*_), and the introgression probability (*φ*_*Y*_). We measure divergence time (τ) in terms of expected number of mutations per site, with τ = *T µ*, where *T* is the divergence time in generations and *µ* is the mutation rate per site per generation. As time *T* and rate *µ* are confounded in analysis of sequence data, only τ is estimable. Each branch on the species tree represents a species or population and is associated with a population size parameter, *θ* = 4*N*_*e*_ *µ*, where *N*_*e*_ is the (effective) population size of the species. A branch on the species tree is also referred to by its daughter node so that branch *RX* is also branch *X*, with population size *θ*_*X*_. Both τ and *θ* are measured as expected number of mutations per site; i.e., one time unit is the expected time to accumulate one mutation per site. At this time scale, coalescence occurs between any two sequences in a population of size *θ* as a Poisson process with rate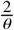

**Figure 1:**
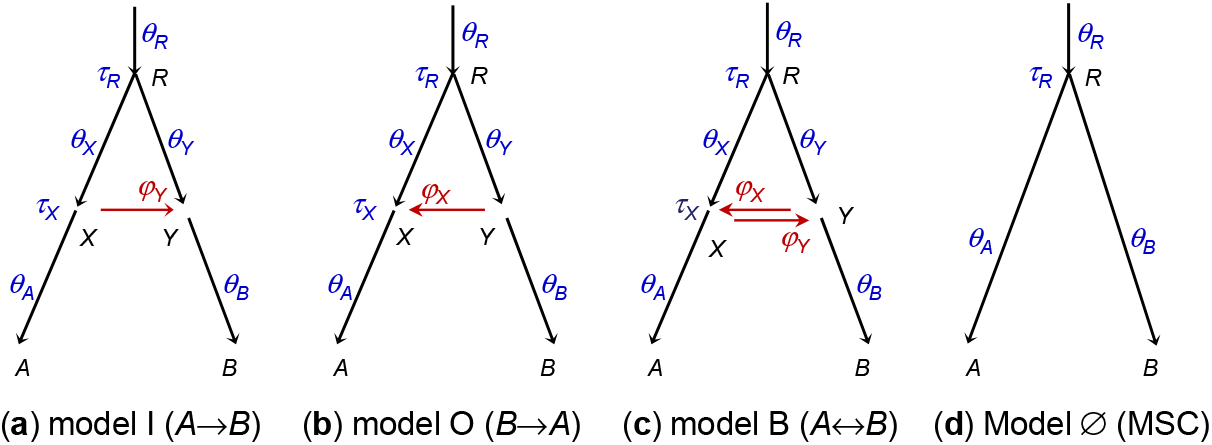
(**a**–**c**) MSci models for two species with different introgression directions showing model parameters: (**a**) *A* → *B* introgression (I for ‘inflow’) with Θ_I_ = (τ_*R*_, τ_*X*_, *θ*_*A*_, *θ*_*B*_, *θ*_*X*_, *θ*_*Y*_, *θ*_*R*_, *φ*_*Y*_), (**b**) *B* → *A* introgression (O for ‘outflow’) with Θ_O_ = (τ_*R*_, τ_*X*_, *θ*_*A*_, *θ*_*B*_, *θ*_*X*_, *θ*_*Y*_, *θ*_*R*_, *φ*_*X*_), or (**c**) bidirectional introgression (B) with Θ_B_ = (τ_*R*_, τ_*X*_, *θ*_*A*_, *θ*_*B*_, *θ*_*X*_, *θ*_*Y*_, *θ*_*R*_, *φ*_*X*_, *φ*_*Y*_). The magnitude of introgression is measured by the introgression probability: *φ*_*Y*_ ≡ *φ*_*A*→*B*_ in **a** and **c** or *φ*_*X*_ ≡ *φ*_*B*→*A*_ in **b** and **c**. Note that in the MSci models studied in this paper, branches *RX* and *X A* represent distinct populations with different population size parameters (*θ*_*X*_, *θ*_*A*_), as are branches *RY* and *Y B*. Horizontal arrows (*XY* and *Y X*) represent introgression events rather than real populations and have no *θ* associated with them. The arrow points to introgression direction in the real world (forward in time). (**d**) MSC model with no gene flow, with Θ_Ø_ = (τ_*R*_, *θ*_*A*_, *θ*_*B*_, *θ*_*R*_).

Each dataset consists of sequence alignments at *L* loci, with *n*_*A*_ sequences from *A* and *n*_*B*_ sequences from *B* at each locus, and with *N* sites in each sequence. Underlying the sequences at each locus is a gene tree with branch lengths (coalescent times), with its probability distribution specified by the MSci model (Yu *et al*., 2014). We assume no recombination among sites in the sequence of the same locus and free recombination between loci; a recent simulation suggests that inference under the MSC is robust to moderate levels of recombination (Zhu *et al*., 2022). Under these assumptions, gene trees and sequence alignments are independent among loci. The data are analyzed under three MSci models that differ in introgression direction: model I with *A* → *B* introgression, model O with *B* → *A* introgression, and model B with bidirectional introgression (*A ⇆ B*) (fig. 1**a**-**c**). The ‘inflow’ (I) and ‘outflow’ (O) labels are used here in anticipation of models involving more than two species to be analyzed later. We use the multilocus sequence data to estimate parameters in the MSci model (Flouri *et al*., 2020). We also use the Bayesian test to detect the presence of gene flow, comparing an MSci model (fig. 1**a**-**c**) with the null model of MSC with no gene flow (fig. 1**d**) (Ji *et al*., 2023).

### The case of two species

#### Distributions of coalescent times and identifiability of introgression direction

We study the distributions of coalescent times between two sequences sampled from the same population (*t*_*aa*_, *t*_*bb*_) or from different populations (*t*_*ab*_). These are analytically tractable and are given in Appendix A. Note that likelihood methods under the MSci model average over the full distribution of the gene tree (*G*) and coalescent times (***t***) for sampled sequences at every locus. However, this distribution depends on the sampling configuration (*n*_*A*_, *n*_*B*_) and is too complex to analyze. Instead we examine the coalescent times (*t*_*aa*_, *t*_*ab*_, *t*_*bb*_) as important summaries of the data, and use their distributions to demonstrate the identifiability of introgression direction, to characterize the information content in estimation of introgression probability, and to predict the behavior of Bayesian parameter estimation (Flouri *et al*., 2020) and the Bayesian test of gene flow (Ji *et al*., 2023).

First we ask whether introgression direction can be inferred using sequence data sampled from extant species. From eq. A2, we have *f*_I_(*t*_*ab*_) = *f*_O_(*t*_*ab*_) for all *t*_*ab*_ > 0, with the parameter mapping 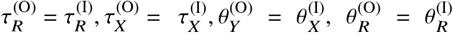 and *φ*_*X*_ = *φ*_*Y*_, where the superscripts indicate the assumed model. Thus *t*_*ab*_ alone cannot distinguish models I and O. In other words, in the case of two species, introgression direction is unidentifiable using data of only one sequence per species per locus (Yang and Flouri 2022, fig. 10; see also Discussion).

However, introgression direction is identifiable if multiple sequences are sampled from *A* and *B*. Information for distinguishing models I and O comes mostly from coalescent times between sequences sampled from the same species (*t*_*aa*_, *t*_*bb*_). If gene flow is *A* → *B*, the coalescent time for sequences from the donor species, *t*_*aa*_, is not affected by the *A* → *B* introgression. If different populations on the species tree have the same size (*θ*_*A*_ = *θ*_*X*_ = *θ*_*R*_), *t*_*aa*_ will have a smooth exponential distribution (e.g., fig. 2**a**, model I). Otherwise the distribution is discontinuous at time points τ_*X*_ and τ_*R*_, because of population size changes. In contrast, *t*_*bb*_ has a mixture distribution, depending on the hybridizing species to which each of the two *B* sequences is traced back on the gene genealogy (i.e., either parental species *RX* or *RY* at node *Y*, fig. 1**a**). Thus the two models make different predictions about coalescent times *t*_*aa*_ and *t*_*bb*_, and the direction of introgression is identifiable when multiple sequences are sampled per species per locus.

**Figure 2:**
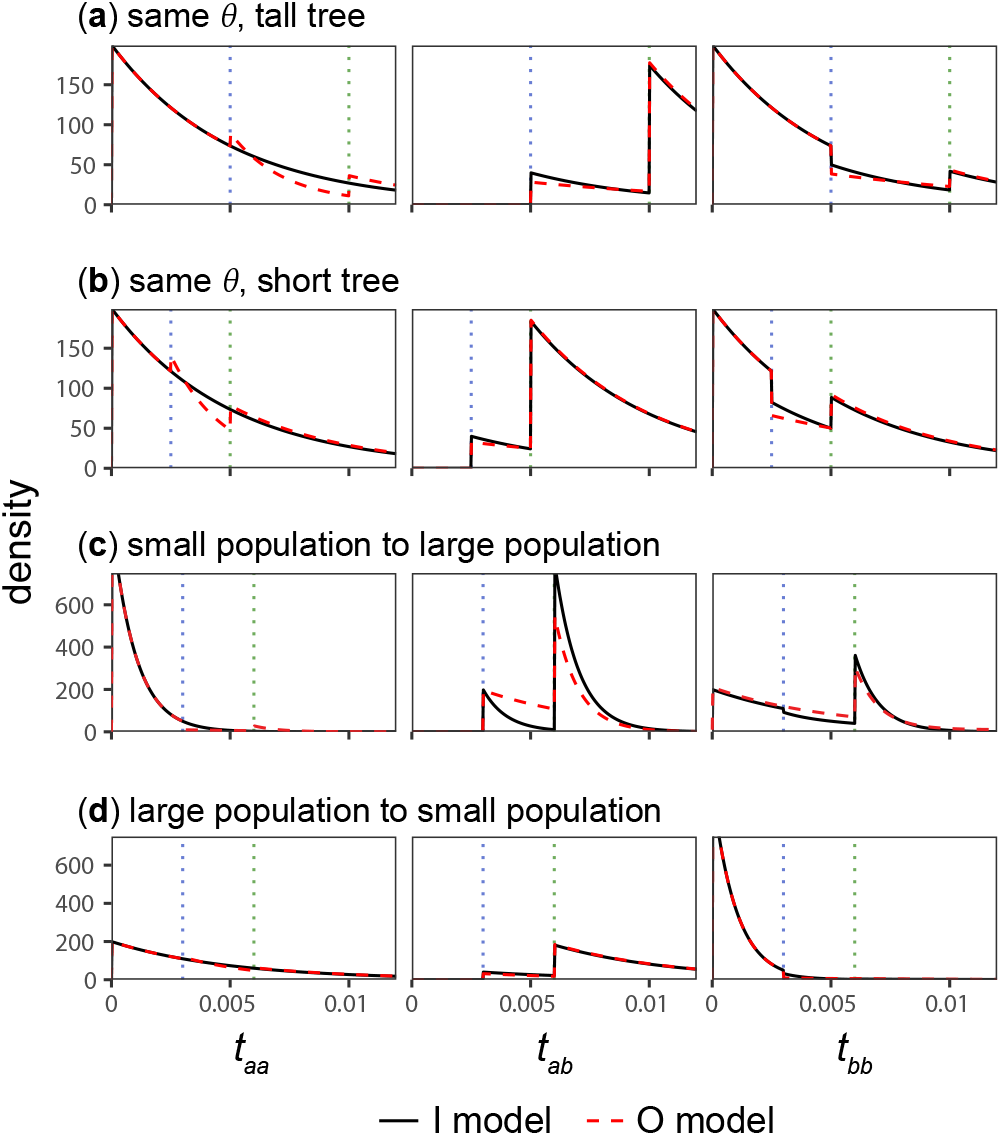
The true (black solid line for model I) and fitted (red dashed line for model O) distributions of coalescent times (*t*_*aa*_, *t*_*ab*_, *t*_*bb*_) for four sets of parameter values (cases **a**–**d**). Data are generated under model I and analyzed under model O of figure 1**a**–**b**. Densities for model I are calculated using the true parameter values (Θ_I_ in table S1); see eqs. A1–A3, while those for model O are calculated using the best-fitting parameter values, approximated by average estimates in bpp analysis of simulated large datasets (with *L* = 4000 loci, *n* = 4 sequences per species per locus and *N* = 500 sites in the sequence) (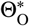 in table S1). Vertical dotted lines indicate discontinuity points at τ_*X*_ and τ_*R*_.

If the introgression direction is specified (that is, under the unidirectional introgression model), introgression probability (e.g., *φ*_*Y*_ given model I) is identifiable using data of one sequence per species per locus. However, the bidirectional introgression (BDI) model (model B) involves an *unidentifiability of the label-switching type*, with two unidentifiable modes or “towers” in the posterior surface if multiple sequences are sampled per species (Yang and Flouri, 2022), or four unidentifiable modes if a single sequence is sampled per species; see Discussion for details.

#### Asymptotic analysis and best-fitting parameter values

We consider multilocus datasets generated under model I with *A* → *B* introgression (fig. 1a) and analyzed under both model I and the misspecified model O with *B* → *A* introgression. We used four sets of parameter values in model I (fig. 1**a**) in the numerical calculation, referred to as cases **a**–**d** (fig. 2, table S1). When the amount of data (the number of loci) *L* → ∞, the maximum likelihood estimates (MLEs) under model I 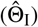 will converge to the true parameter values, i.e. 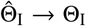. Under model O, the MLEs 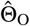 will converge to the *best-fitting* or *pseudo-true* parameter values 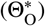, which minimize the Kullback-Leibler (KL) divergence from the true model to the fitting model: 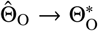 (e.g., Yang and Zhu, 2018). With arbitrary data configurations, it does not seem possible to calculate 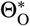 analytically. Instead we use as a substitute the averages of posterior means of parameters in bpp analysis of simulated large datasets (with *L* = 4000 loci, *n*_*A*_ = *n*_*B*_ = 4 sequences per species per locus and *N* = 500 sites per sequence), shown in table S1. At this data size, average estimates under the true model I are extremely close to the true values, i.e. 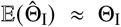 (table S1), suggesting that the average estimate under model O may also be very close to the infinite-data limits, 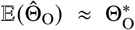. We aim to understand the estimates 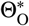 by comparing the true distributions of coalescent times under model I, *f*_I_(*t*_*aa*_), *f*_I_(*t*_*ab*_), and *f*_I_(*t*_*bb*_) (eqs. A1–A3), with fitted distributions *f*_O_(*t*_*aa*_), *f*_O_(*t*_*ab*_), and *f*_O_(*t*_*bb*_), calculated using 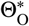. In other words, we treat the true distributions of coalescent times under model I as data, and attempt to derive parameter estimates under the fitting model O to achieve the best fit.

Our theory is summarized in table 1. Note that parameters τ_*R*_, τ_*X*_, *θ*_*A*_, *θ*_*B*_, *θ*_*R*_ in model O are typically well estimated. Introgression time 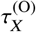 is largely determined by the smallest coalescent time between sequences from the two species (*t*_*ab*_), while the discontinuity in the distributions of *t*_*aa*_, *t*_*ab*_, *t*_*bb*_ should be informative about 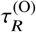. Thus we expect estimates of those parameters to be close to the true values despite the model misspecification: 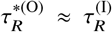 and 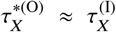 Population sizes 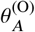 and 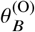 for the extant species should be well-estimated from multiple samples from the same species, while 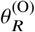 should be well estimated based on coalescent events in the root population. Below we focus on parameters 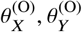, and *φ*_*X*_, which are harder to estimate.

**Table 1.** Features of the data that are informative about parameters in the wrong model O when data are generated under model I with parameter Θ^(I)^ (fig. 1a)

| Parameter estimates in model O | Information in data | Notes |
| --- | --- | --- |
| (a) Introgression time:<br>$\hat{\tau}_X^{(O)} \approx \tau_X^{(I)}$ | $\min\{t_{ab}\}$ | In the fitting model O, $t_{ab} > \tau_X^{(O)}$. Thus introgression time $\tau_X^{(O)}$ is determined by the minimum between-species coalescent time ($t_{ab}$). |
| (b) Species divergence time:<br>$\hat{\tau}_R^{(O)} \approx \tau_R^{(I)}$ | Discontinuities in $f(t_{aa})$, $f(t_{ab})$, and $f(t_{bb})$ | Species divergence time is informed by the discontinuities in the coalescent times ($t_{aa}$, $t_{ab}$, $t_{bb}$). |
| (c) Population sizes for extant species:<br>$\hat{\theta}_A^{(O)} \approx \theta_A^{(I)}$, $\hat{\theta}_B^{(O)} \approx \theta_B^{(I)}$ | $f(t_{aa})$ and $f(t_{bb})$ over $(0, \tau_X)$ | Population size for an extant species is easily estimated by the heterozygosity in the species. |
| (d) Population sizes for ancestral species not involved in introgression:<br>$\hat{\theta}_R^{(O)} \approx \theta_R^{(I)}$ | $f(t_{aa})$, $f(t_{ab})$, and $f(t_{bb})$ over $(\tau_X, \infty)$ | Population sizes for ancestral species not involved in introgression are determined by coalescent times in the ancestral species. |
| (e) Ancestral population size:<br>$\hat{\theta}_X^{(O)} < \theta_X^{(I)}$ | $f(t_{aa})$ over $(\tau_X, \tau_R)$ | The fitting model (O) predicts a deficit of coalescence of *A* sequences ($t_{aa}$) over $(\tau_X, \tau_R)$ due to introgression but there is no such deficit or in the true model I or in the data. Having a larger coalescent rate (or smaller population size $\theta_X^{(O)}$) in model O thus helps to improve the model fit to coalescence of *A* sequences in the data. |
| (f) Ancestral population size:<br>$\hat{\theta}_Y^{(O)} > \theta_Y^{(I)}$ | $f(t_{bb})$ over $(\tau_X, \tau_R)$ | There is a deficit of coalescence of *B* sequences ($t_{bb}$) over $(\tau_X, \tau_R)$ in the true model I or in the data. Having a smaller coalescent rate (or larger population size $\theta_Y^{(O)}$) in model O thus helps with the model fit. |
| (g) Introgression probability:<br>$\hat{\varphi}_X(1 - e^{-2\Delta\tau/\hat{\theta}_Y^{(O)}}) \approx \varphi_Y(1 - e^{-2\Delta\tau/\theta_X^{(I)}})$<br>(eq. 1):<br>$\hat{\varphi}_X > \varphi_Y$ if $\hat{\theta}_Y^{(O)} > \theta_X^{(I)}$,<br>$\hat{\varphi}_X < \varphi_Y$ if $\hat{\theta}_Y^{(O)} < \theta_X^{(I)}$. | $f(t_{ab})$ over $(\tau_X, \tau_R)$ | Introgression probability is informed by the amount of between-species coalescence ($t_{ab}$) over $(\tau_X, \tau_R)$. Eq. 1 means the same amount of coalescence in species *Y* in the fitting model as in species *X* in the true model. |

Note.— The introgression model assumes different population sizes ( $\theta$ s) for species on the tree (fig. 1); the behavior of method may differ if all populations are assumed to have the same size. Also the reasoning here is based on coalescent times and ignores sampling errors in gene trees and estimated coalescent times in analysis of sequence data.

First, by considering the distributions of *t*_*aa*_, we predict 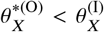 (table 1). In the true model I, both *A* sequences enter *X* and may coalesce during (τ_*X*_, τ_*R*_). In the fitting model O, the two *A* sequences may be separated into different populations due to introgression (one in *X* and the other in *Y*), so they may not coalesce in (τ_*X*_, τ_*R*_) as often. Thus having 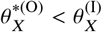 will increase the coalescent rate in *X* and help to fit model O to *f* (*t*_*aa*_) over (τ_*X*_, τ_*R*_).

Next from *t*_*bb*_, we predict 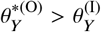 (table 1). In the true model, *A* → *B* introgression reduces the chance of coalescence between sequences from *B* during (τ_*X*_, τ_*R*_). In the fitting model, both *B* sequences enter *Y*, leading to a higher chance of coalescence during (τ_*X*_, τ_*R*_). Thus having 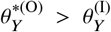 helps to reduce the chance of coalescence in (τ_*X*_, τ*R*)

Finally, by matching the amount of coalescence between sequences *a* and *b* over the time interval (τ_*X*_, τ_*R*_), or by matching the probability densities *f*_I_(*t*_*ab*_) and *f*_O_(*t*_*ab*_) over τ_*X*_ < *t*_*ab*_ < τ_*R*_, we have approximately

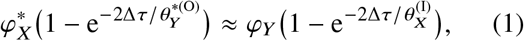

where Δτ = τ_*R*_ − τ_*X*_ is assumed to be the same under models I and O based on the arguments above. Eq. 1 predicts that more gene flow will be inferred under model 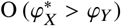 when 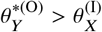; if the coalescent rate between sequences *a* and *b* during (τ, τ) is lower in the fitting model than in the true model, a higher 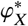 than the true *φ*_*Y*_ will increase the chance of such coalescence and achieve a better fit to *f*_I_(*t*_*ab*_). Similarly, less gene flow is expected (with 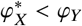) if 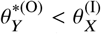.

Eq. 1 predicts 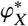 to be 0.31, 0.35, 0.44 and 0 22 for cases **a**–**d**, respectively, compared with the inferred values 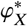 of 0.27, 0.30, 0.98 and 0.17 (table S1). The approximation is reasonably good except for case **c**, where 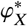 was very high. We discuss these cases further when describing simulation results below.

#### Simulation results under the true models I and B: parameter estimates have drastically different precisions

To verify and extend our theoretical analysis, we simulated datasets under model I (fig. 1**a**) and analyzed them under models I, O, and B (fig. 1**a**-**c**) using four sets of parameter values. Each dataset consists of *L* = 250, 1000 or 4000 loci, with *n*_*A*_ = *n*_*B*_ = 4 sequences sampled per species per locus and *N* = 500 sites in the sequence. Posterior means and 95% highest-probability-density (HPD) credibility intervals (CIs) are plotted in figure 3 (see also table S1 for *L* = 4000).

**Figure 3:**
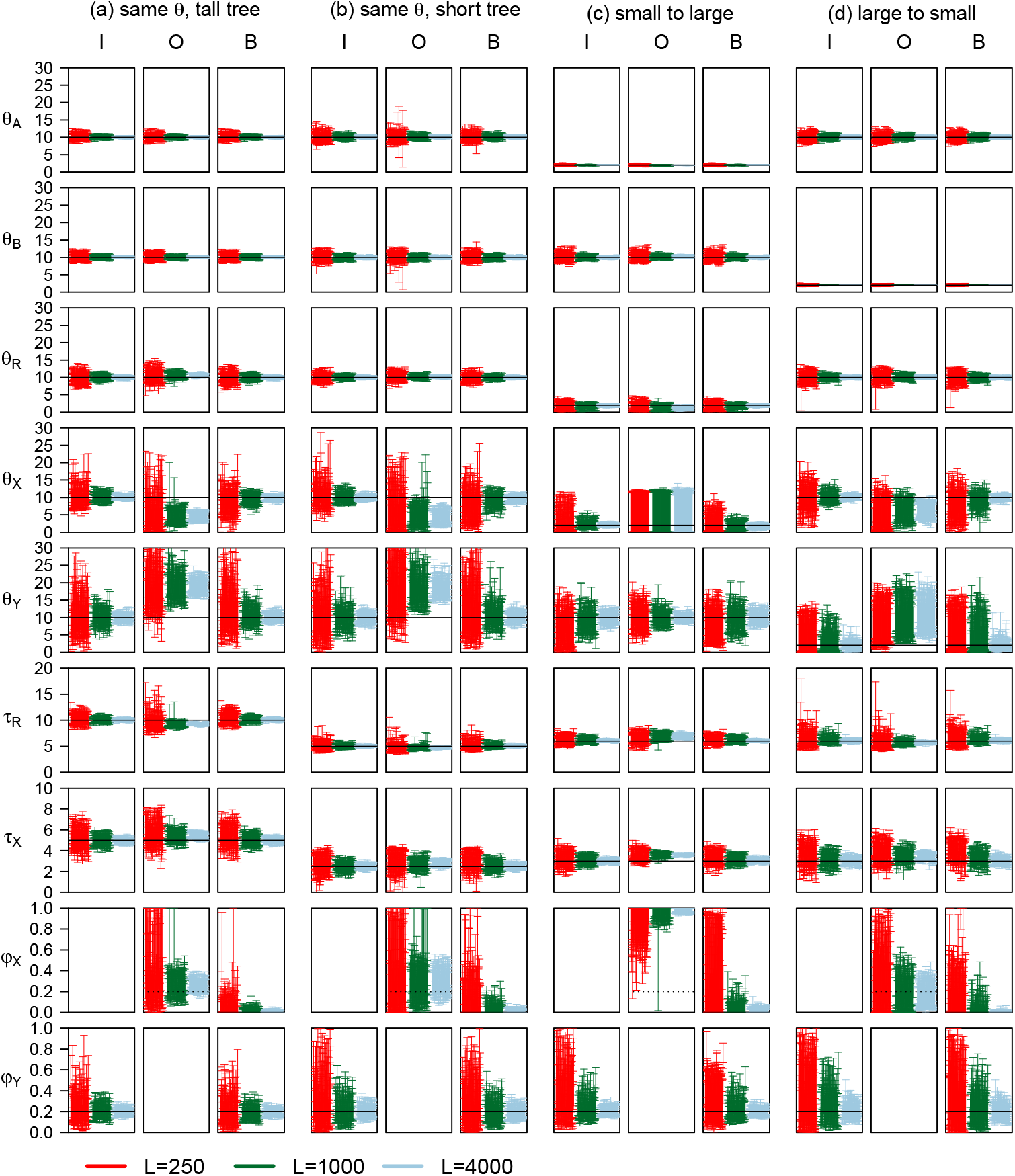
The 95% HPD CIs for parameters in 100 replicate datasets (each of *L* loci) simulated under model I and analyzed under models I, O, and B of figure 1**a**–**c**. Four sets of parameter values are used (cases **a**–**d**) (table S1). Parameters *θ*s and τs are multiplied by 10^3^. Black solid lines indicate the true values. Dotted lines for *φ*_*X*_ in model O indicate the true value of *φ*_*Y*_ in model I.

Model I is the true model, so that the performance under this model constitutes the best-case scenario. Indeed all parameters are well estimated, with the posterior means approaching true values and the CI width approaching 0 when the amount of data *L* → ∞ (fig. 3 and table S1, cases **a**-**d**, model I). However, the amount of information in the data varies hugely for different parameters, as reflected in the relative error, measured for example by the CI width divided by the true value. Population sizes for extant species (*θ*_*A*_, *θ*_*B*_) are much better estimated than those for ancestral species (*θ*_*X*_, *θ*_*Y*_). Divergence times (τ_*R*_, τ_*X*_) are well estimated as well. Introgression probability (*φ*_*Y*_) has substantial uncertainties with wide CIs but with *L* = 4000 loci in the data, the estimates are fairly precise, suggesting that thousands of loci are necessary to estimate introgression probability precisely. The results parallel those found in a previous simulation examining the impact of data size (such as the number of loci, the number sequences per species, and the number of sites) on inference under the MSci model (Huang *et al*., 2020).

Model B allows bidirectional introgression and thus is a correct model although it is over-parametrized with an extra parameter *φ*_*X*_. As the amount of data increases, 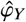 should converge to the true value while 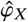 to 0. Estimates of other parameters are very similar to those under model I, and the CI widths under models I and B are also very similar. In particular, *φ*_*Y*_ is estimated with similar precision in the two models. In large datasets of *L* = 4000 loci, the average CI width is 0.07, 0.12, 0.08, and 0.16 for cases **a**–**d** under model I, compared with 0.07, 0.12, 0.09, 0.17 under model B. Even in small or intermediate datasets with *L* = 250 or 1000 loci, the CIs for *φ*_*Y*_ are similar between the two models. Thus over-parametrization incurred little cost to statistical performance of model B. This might seem surprising, because, given the difficulty of inferring introgression direction, one might expect the assumed incorrect *B* → *A* introgression in model B would interfere with estimation of *φ*_*Y*_ in the correct direction, so that 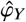 would have a much larger variance under model B than under model I. However, information concerning *φ*_*Y*_ is largely determined by (i) the number of sequences reaching the hybridization node *Y* and (ii) the ease with which one can tell the parental path taken by each *B* sequence at *Y* (see the next subsection for detailed discussions). Thus there may be little difference in information content about *φ*_*Y*_ between models I and B. Computationally, model B is much more expensive than model I due to sampling an extra parameter in the Markov chain Monte Carlo (MCMC) algorithm and to MCMC mixing issues (Yang and Flouri, 2022).

#### Information content for estimating introgression probability under the true model

Here we consider estimation of introgression probability *φ*_*Y*_ in model I in the four cases (fig. 3, cases **a**–**d**, model I). We characterize the amount of information concerning *φ*_*Y*_ when the correct model is assumed, and explain why *φ*_*Y*_ was much better estimated in case **a** (same *θ* tall tree) than in case **b** (same *θ* short tree), and in case **c** (small to large) than in case **d** (large to small) (fig. 3, table S1: cases **a**–**d**, model I), even though the data size is the same and the true *φ*_*Y*_ is the same (0.2) in all cases. The theory is also useful for understanding later simulation results for larger species trees.

Consider tracing the genealogical history of sequences at a locus backwards in time. When sequences from *B* reach the hybridization node *Y* (fig. 1a), there is a binomial sampling process, with each sequence taking the horizontal (introgression) parental path (into *RX*) with probability *φ*_*Y*_ and the vertical parental path (into *RY*) with 1 − *φ*_*Y*_. However there are two differences from a typical binomial sampling. First, the number of *B* sequences reaching node *Y* is a random variable. Second, the outcome of the sampling process (i.e., the parental path taken by the sequence) is not observed but instead reflected in the gene tree and coalescent times (and thus in mutations in the sequences). Using a coin-tossing analogy, the number of coin tosses is random, and the outcome of the toss is visible only probabilistically. If a *B* sequence coalesces with an *A* sequence during the time interval (τ_*X*_, τ_*R*_), it will be clear that the *B* sequence has taken the introgression parental path.

Thus the amount of information in the data concerning *φ*_*Y*_ is determined by two factors: (i) the number of *B* sequences reaching *Y* and (ii) the ease with which one can tell the parental path taken by each *B* sequence at *Y*. The number of *B* sequences reaching *Y* at the locus is given as *n*_*B*_ − *c*_*B*_, where *n*_*B*_ is the number of *B* sequences sampled at the locus and *c*_*B*_ is the number of coalescent events among them in *B* before reaching *Y*. The distribution of *n*_*B*_ − *c*_*B*_ can be easily calculated as a function of *n*_*B*_ and 2τ_*Y*_ /*θ*_*B*_, the length of branch *B* measured in coalescent units (Tavaré, 1984: eqs. 6.1 & 6.2; Wakeley, 2009: eqs. 3.39 & 3.41). More *B* sequences will reach *Y* the larger *n*_*B*_ is and the smaller 2τ_*Y*_ /*θ*_*B*_ is. As a result, it will be harder to estimate *φ*_*Y*_ if introgression is older (larger τ_*Y*_).

The second factor — the ease with which one can tell the parental path taken by each *B* sequence at *Y* — concerns the probability that two sequences entering *X* coalesce in *X* before reaching *R*; there is more information about *φ*_*Y*_ the longer the internal branch *RX* is or the smaller the population size *θ*_*X*_ is (fig. 1**a**). This may be seen by considering the special case where the data consist of one sequence per species per locus and where the true coalescent time (*t*_*ab*_) is available at each locus. Then the information content for estimating *φ*_*Y*_ may be measured by the Fisher information, given by

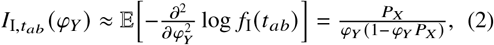

where the expectation is with respect to *t*_*ab*_ (eq. A3), and where 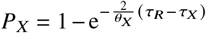 is the probability that two sequences (*a, b*) entering population *X* coalesce in *X*. The asymptotic variance of the estimate 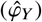 is

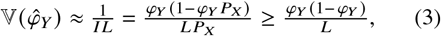

with equality holding if *P*_*X*_ = 1. There is thus more information for estimating *φ*_*Y*_ the closer *P*_*X*_ is to 1, or in other words if the branch length in coalescent units, 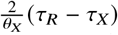, is greater. Increasing the number of sequences reaching *Y* per locus (*n*_*B*_ − *c*_*B*_) may be expected to have a similar effect to increasing the number of loci (*L*) as both increases the binomial sample size. Eq. 3 thus suggests that increasing *P*_*X*_ is more effective in reducing 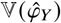 than increasing the number of loci (*L*) by the same factor, which is in turn more effective than increasing the number of sampled sequences per locus (*n*_*B*_) by the same factor. For example, doubling *n*_*B*_ − *c*_*B*_ reduces the variance for 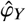 by a half, but doubling *P*_*X*_ reduces the variance by more than a half.

In our simulation (fig. 3, model I), the introgression probability *φ*_*Y*_ was better estimated in case **a** (same *θ* tall tree) than in case **b** (same *θ* short tree). At *L* = 4000, the 95% HPD CI width was 0.07 for case **a**, and 0.12 for case **b**. Consider the two factors. First, in case **a** (tall tree), branch *Y B* is longer, with length 2τ_*Y*_ /*θ*_*B*_ in coalescent units, with a smaller number of sequences reaching *Y* than in case **b** (short tree). Indeed, given *n*_*B*_ = 4 sequences from *B*, the probability that *n*_*B*_−*c*_*B*_ = 1, 2, 3, and 4 sequences remain by time τ_*Y*_ is 0.388, 0.515, 0.095, and 0.002, respectively in case **a**, with an average of 1.71 (fig. S1). For the short tree of case **b**, the corresponding probabilities are 0.122, 0.481, 0.347, and 0.050, with average 2.32. The average number of sequences reaching *Y* differ by a factor 1.36. Second, in case **a** (tall tree), any *B* sequence reaching *Y* and taking the left parental path is more likely to coalesce with *A* sequences in *X* than in case **b** (short tree), with *P*_*X*_ = 1 − e^−1^ = 0.632 in case **a** and *P*_*X*_ = 1 − e^−0.5^ = 0.393 in case **b**, differing by a factor of 1.61. As increasing *P*_*X*_ is more effective than increasing *n*_*B*_ − *c*_*B*_ (eq. 3), *φ*_*Y*_ was more precisely estimated (with smaller variance) in case **a** than in **b** (fig. 3, table S1).

The difference between case **c** (small to large) and case **d** (large to small) was even greater, with *φ*_*Y*_ much better estimated in **c** (fig. 3). At *L* = 4000, the CI width was 0.08 for case **c** and 0.16 for case **d** (table S1). In case **c** more *B* sequences reach *Y* because of the large *θ*_*B*_ than in case **d**. Furthermore *B* sequences reaching *Y* into *X* have a high chance of coalescence with other sequences in population *X*. Both effects make it easier to estimate *φ*_*Y*_ in case **c** than in case **d** (eq. 3). It is thus easier to estimate *φ*_*Y*_ if introgression is from a small population to a large one than in the opposite direction (fig. S2). Note that *φ*_*Y*_ is the proportion of immigrants in the recipient population, so that with the same *φ*_*Y*_, there are many more migrants in case **c** than in **d**.

#### Parameter estimation under misspecified introgression direction

When model O was used to analyze data simulated under model I (fig. 1), the introgression direction is misspecified. As discussed above (table 1), species divergence and introgression times (τ_*R*_, τ_*X*_) are well estimated despite misspecification, as are population sizes for extant species and for the root (*θ*_*A*_, *θ*_*B*_, *θ*_*R*_). Indeed those parameters are estimated with the same precision under models O and I (fig. 3).

Here we focus on parameters *φ*_*X*_, *θ*_*X*_, *θ*_*Y*_ (fig. 3, model O). Our arguments from the asymptotic analysis (table 1) also apply, although in simulations the results are affected by random sampling errors due to finite data size.

In cases **a & b**, all populations have the same size. Biases in parameter estimates under model O are well predicted by the theory (table 1): based on coalescent times *t*_*aa*_, *t*_*bb*_, and *t*_*ab*_, we expect 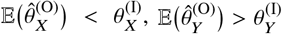,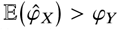

In case **c** (small to large), introgression is from a small population to a large one. As the coalescent rate for sequences *a* and *b* over (τ_*X*_, τ_*R*_) is much slower in the fitting model than in the true model, consideration of *t*_*ab*_ predicts a large 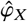 or a small 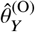 (table 1). Consideration of *t*_*bb*_ suggests 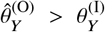 will compensate for reduced coalescence between *B* sequences caused by the *A* → *B* introgression (table 1). Thus predictions about 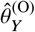 based on *t*_*ab*_ and *t*_*bb*_ are somewhat conflicting. In the simulation, 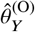, is close to 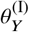, much larger than 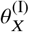. The estimate is 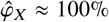 (table S1). The extreme estimate causes small biases in τ_*R*_ and τ_*X*_ and poor estimates of 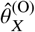 (fig. 3).

Case **d** (large to small) assumes introgression from a large population to a small one (fig. 1**a**). We expect 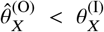 based on *t*_*aa*_, and 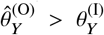 based on *t*_*bb*_ (table 1). Moreover, the larger source population in the true model 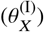 means *t*_*ab*_ is less common in (τ_*X*_, τ_*R*_), with most coalescence occurring in the common ancestor *R*. Thus based on *t*_*ab*_ we predict a larger 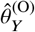 or a smaller 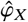 to reduce the amount of coalescence in (τ_*X*_, τ_*R*_) in the fitting model (eq. 1). Thus considerations of both *t*_*bb*_ and *t*_*ab*_ suggest 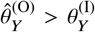 Depending on whether 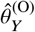 is smaller or greater than 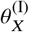, the introgression probability 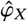 may be greater or smaller than the true *φ*_*Y*_, according to eq. 1. In our setting, 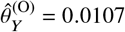, slightly greater than 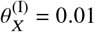, and 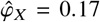, slightly smaller than *φ*_Y_ = 0.2 (table S1).

#### Bayesian test of introgression: power and false positive rate

We applied the Bayesian test of gene flow (Ji *et al*., 2023) to the data analyzed in figure 3. We are interested in the power of the test under the correct model I. Also we ask how often the test is significant if it is conducted under model O, with introgression direction misspecified.

Note that the behavior of the test or the asymptotic behavior of posterior probabilities of the compared models is determined by the parameter values in the limit of *L* → ∞ (Yang and Zhu, 2018). If data are simulated under model I (with *φ*_*Y*_ > 0) and analyzed under model I, the posterior probability for the true model I should approach 1, the Bayes factor in support of model I against model Ø of no gene flow (fig. 1**d**) *B*_IØ_ → ∞, and the power of the test should approach 100%, when the data size *L* → ∞ (Yang and Zhu, 2018). If the data are simulated under model I and analyzed under model B, the power for testing *φ*_*Y*_ (which has the true value *φ*_*Y*_ > 0) should approach 100%, and the false positive rate for testing *φ*_*X*_ (which has the true value *φ*_*X*_ = 0) should approach 0, when the data size *L* → ∞.

If the data are generated under model I and analyzed under model O, both the null and alternative models are incorrect. According to our analysis 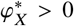, and model O is a ‘less wrong’ model than model Ø, judged by the Kullback-Leibler divergence (Yang and Zhu, 2018). Thus when *L* → ∞, *B*_Oø_ → ∞, and the probability of rejecting *H*_ø_ : *φ*_*X*_ = 0 will approach 100%. Here the biological interpretation of test results is somewhat ambiguous. If one emphasizes the fact that model O allows gene flow while model Ø does not, detecting gene flow may be considered a correct result. However, if one emphasizes misspecification of introgression direction in model O, accepting model O may be considered a rather severe false positive error. In this paper, we use the second interpretation.

The MCMC samples generated in bpp runs of figures 3 were processed to calculate the Bayes factor *B*_10_ in favor of the introgression model (*H*_1_, fig. 1**a**-**c**) against the null MSC model of no gene flow (*H*_0_, fig. 1**d**) via the Savage-Dickey density ratio (see Methods). The results are summarized in figure S2, where a 1% significance level was used (i.e., the test is significant if *B*_10_ > 100). When the data were simulated and analyzed under model I and with *L* = 250 loci in the data, power was between 60–100% (fig. S2, cases **a**-**d**, model I). In such small datasets, *φ*_*Y*_ was poorly estimated with extremely wide CIs (fig. 3, cases **a**–**d**, model I). At *L* = 1000 loci, power was 100% in all four cases. It is thus easier to detect gene flow than to estimate its magnitude reliably. As with our findings on estimation of *φ*_*Y*_, it is easier to detect gene flow in case **a** (tall tree) than in case **b** (short tree), and in case **c** (small → large) than in case **d** (large → small) (fig. S2).

When the data are analyzed under model O, with the introgression direction misspecified, the false positive error is comparable to the power in the analysis under true model I (fig. S2, cases **a**-**d**, model O). When the data are analyzed under model B, power to detect the *A* → *B* introgression is slightly lower than under model I, also reaching 100% at *L* = 1000, while the false positive rate for detecting the non-existent *B* → *A* introgression is low, below the nominal 1%.

### Additional information that results from including a third species

Given two species (*A, B*) with introgression from *A* → *B* at the rate of *φ* (fig. 1**a**), we consider the information gain for estimating *φ* from including a third species (*C*). There are five branches on the two-species tree onto which *C* can be attached (fig. 4**a**–**e**): (**a**) the root population, (**b, c**) the source and target populations before gene flow, and (**d, e**) the source and target populations after gene flow. Case **c** is one of ‘inflow’, with gene flow from the outgroup species (*A*) into one of the ingroup species (*B*), while **b** represents ‘outflow’, with gene flow from an ingroup species (*A*) into the outgroup (*B*). Note that in all cases the correct MSci model is used in the analysis, so that the estimate (posterior mean) of *φ* will converge to the true value (which is 0.2). However, the information content may differ among the five cases. As in the case of two species, the amount of information concerning *φ* is determined by two factors: (i) the number of sequences reaching the hybridization node and (ii) the ease with which one can tell the parental path taken by each sequence at the hybridization node. When introgression is between nonsister species, information concerning the parental path taken by each sequence may be in the change of gene-tree topology rather than in the change of between-species coalescent time.

**Figure 4:**
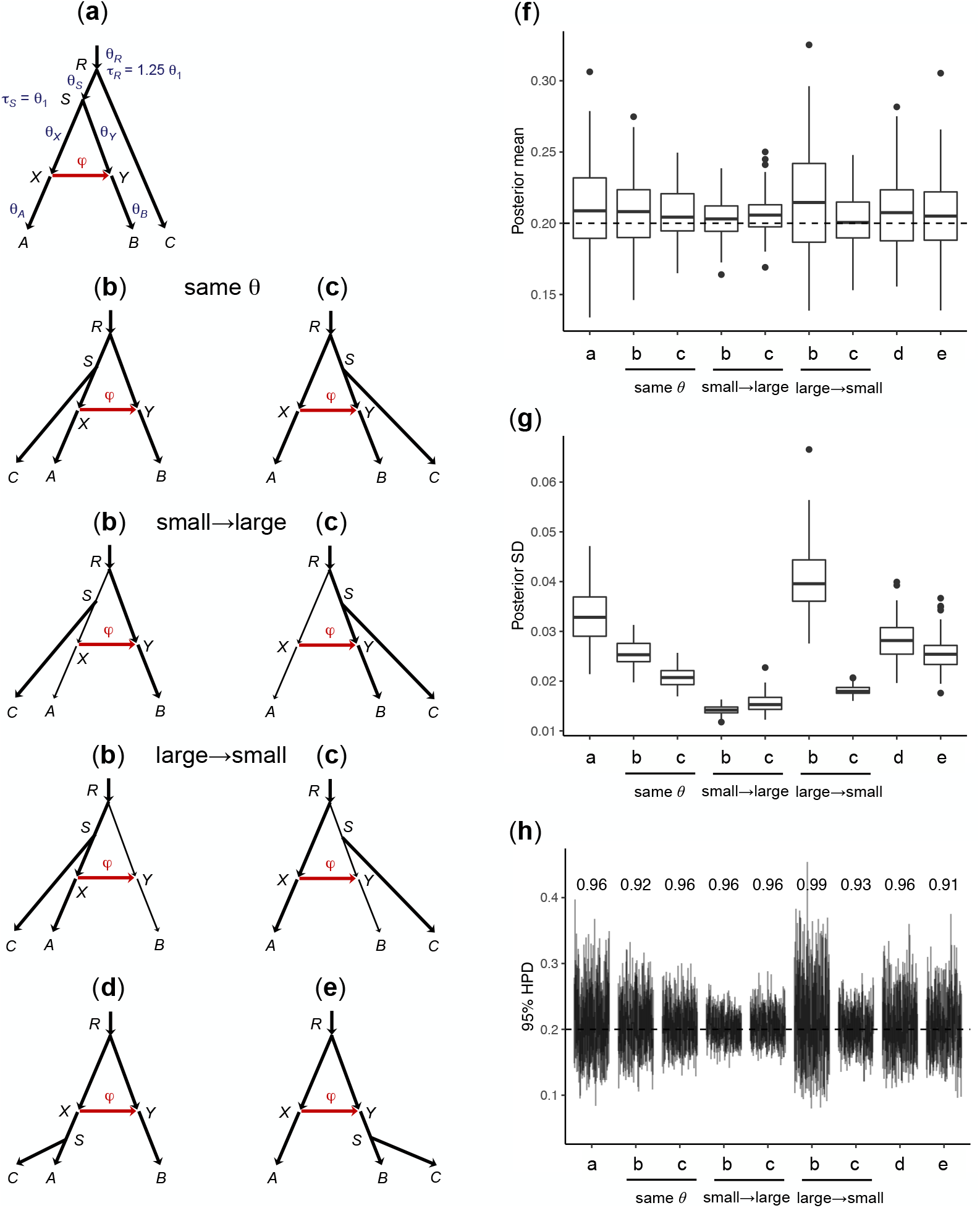
(**a**–**e**) MSci models for three species (*A, B, C*), with introgression from *A* to *B*, obtained by adding a third species *C* onto the two-species tree of figure 1**a** at five possible locations: (**a**) root population, (**b, c**) source and target populations before gene flow, and (**d, e**) source and target populations after gene flow. (**f**) Box plots of the posterior means for *φ* among 100 replicate datasets simulated under each of the five cases (**a**–**e**). The dashed line indicates the true value (*φ* = 0.2). (**g**) Box plots of the posterior SD for *φ*. (**h**) 95% HPD CIs for *φ*, with the CI coverage above the CI bars. See figure S3 for CIs for other parameters.

We assumed the same population size *θ*_1_ = 0.01 for all populations, but examined the impact of different population sizes in cases **b** and **c**. We simulated 100 replicate datesets in each case. The posterior means, the posterior standard deviation (SD), and the width of the HPD CI for *φ* are summarized in figure 4**f**-**h**. The 95% CIs for other parameters are shown in figure S3.

#### Equal population sizes on the species tree

If all populations on the species tree have the same size (*θ*), we expect the amount of information for estimating *φ* to be in the order **a** *≺* **d** *≺* (**b, e**) *≺* **c**, with the order of **b** and **e** undecided (fig. 4**f**–**h**).

First, **a** *≺* **d**. Cases **a** and **d** are the least informative. Adding an outgroup species *C* in case **a** adds little information about *φ*. In **d**, the *C* sequences may reach node *X* and coalesce with a *B* sequence in *RX*, providing information about whether sequences from *B* take the introgression parental path at node *Y*. Thus we expect more information in the data in **d** than in **a**.

Next, **d** *≺* **b**. The number of *B* sequences reaching node *Y* is the same in the two cases, so the only difference is in the difficulty of inferring the parental path taken by *B* sequences at *Y*. In case **b**, coalescence of a *B* sequence with an *A* sequence causes a change to gene tree topology. In case **d**, introgression does not cause such topological change to the gene tree. The information content may thus be higher in **b** than in **d**.

Next, **d** *≺* **e**. In case **e**, sequences from both *B* and *C* may reach the hybridization node *Y* while in **d** only sequences from *B* may reach *Y*, so that the sample size at node *Y* is larger (less than twice as large) in **e** than in **d**. In **d**, more sequences enter population *RX*, increasing slightly the probability of coalescence for any *B* sequence that takes the introgression parental path at *Y*, but this effect may be less important than that of increased sample size in **e**.

Next, **b** *≺* **c** (i.e., it is easier to infer inflow than outflow). In both cases, the number of *B* sequences reaching node *Y* or the sample size at *Y* is the same. However, the two cases differ in the ease with which one can tell the parental path taken by each *B* sequence at *Y*. In **c**, coalescence of a *B* sequence with an *A* sequence over (τ_*X*_, τ_*R*_) causes a change to gene tree topology. In **b** such topology change occurs only if the coalescence occurs in the shorter time interval (τ_*X*_, τ_*S*_), and the resulting gene tree is harder to infer because of the shorter internal branch. It is thus harder to resolve the parental path taken by each *B* sequence at *Y* in **b** than in **c**, and the data are less informative about *φ* in **b**. It is harder to infer outflow than inflow.

Finally, **e** *≺* **c**. In case **c**, introgression leads to changes in gene tree topology whereas in **e**, more sequences reach *Y* with a larger sample size. The relative effects depend on the parameter values. In the simulation here, the increased sample size was less effective than the gene tree topology change (fig. 4**g & h**, case **c** same-*θ* vs. case **e**). Note that in **e** the data are more informative about *φ* the closer τ_*S*_ is to τ_*Y*_, and in both **c** and **e** the data are more informative the smaller τ_*X*_ is.

#### Different population sizes on the species tree

For cases **b** (outflow) and **c** (inflow), we also consider different population sizes. The results are shown in figure 4**f**–**h**.

First, in case **b**, *φ* is most poorly estimated in the large→small setting, much better estimated in the same-*θ* (or large→large) setting, and best in the small→large setting. This can be explained easily by the theory we developed in analysis of the two species case: a large recipient population means many sequences reaching the hybridization node *Y* and a large sample size, while a small donor species (*θ*_*X*_) means fast coalescence and easy determination of the parental path taken at node *Y*. For example, the probability that more than one *B* sequence reaches *Y* is 0.613 in case **b** (same *θ* or small→large), and 0.012 in case **b** (large→small), with a large difference in the sample size.

Similarly in case **c** (inflow), *φ* is more poorly estimated in the large→small and same-*θ* (large→large) settings, and was better in the small→large setting. The differences among the three settings are much smaller than in case **b**.

Although case **b** outflow is less informative about *φ* than **c** inflow in the case of same-*θ*, the order is reversed in the small→large setting (fig. 4). The same number of *B* sequences reaches node *Y* in both cases, so the difference must be due to the different levels of difficulty by which one can tell the parental paths taken by *B* sequences at node *Y*. In **b**, *B* sequences taking the introgression parental path go through the small population *SX* and may coalesce at a high rate with sequences from *A* (which lead to changes to the gene tree topology informative about introgression), and with sequences from both *A* and *C* in population *RS*. In **c**, *B* sequences taking the vertical parental path may coalesce in population *RS* with *C* sequences, but given that both populations *SY* and *RS* are large, this effect may be expected to be minor. While multiple factors can have opposing effects on the relative information content concerning *φ* in cases **b** versus **c** small→large, the data are more informative in case **b** than in **c** overall.

### Simulation results in the case of four species

We conducted simulations under the MSci models of figure 5 for four species on the species tree ((*A*, (*B, C*)), *D*), with introgression between nonsister species *A* and *B* in different directions: inflow (I), outflow (O), and bidirectional introgression (B). Either the same population size was assumed for all species on the species tree or different population sizes were assumed. The simulated data were analyzed under the same three models (I, O, B), resulting in nine combinations. Posterior means and 95% HPD CIs are summarized in figure S4 for the case of equal population sizes and in figure S5 for different population sizes. The results for the large datasets of *L* = 4000 are summarized in tables S2 & S3. We also applied the Bayesian test of introgression (Ji *et al*., 2023) to the simulated data. The results are summarized in figures S6 & S7.

**Figure 5:**
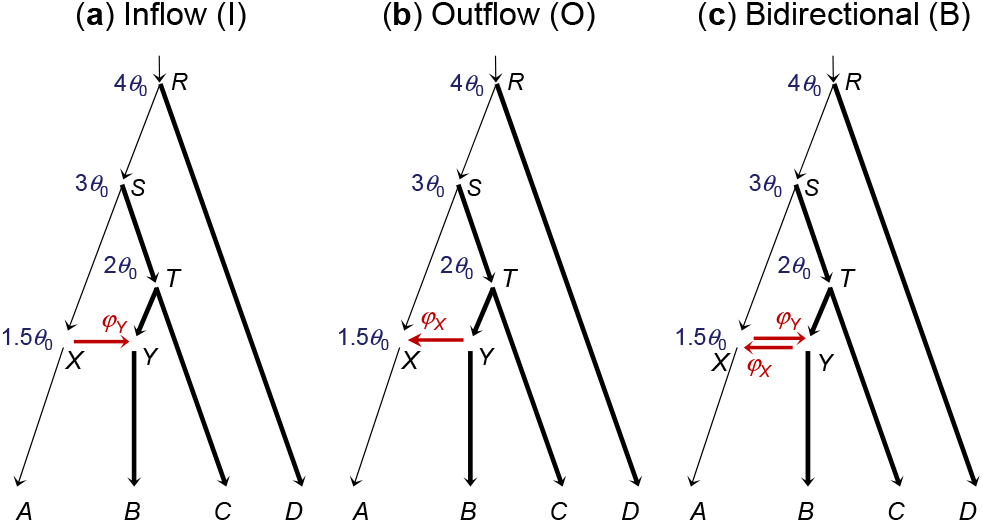
**(4s-trees)** Three MSci models for four species differing in introgression direction assumed to simulatie and analyze data: (**a**) inflow from *A* to *B* (I); (**b**) outflow from *B* to *A* (O); and (**c**) bidirectional introgression between *A* and *B* (B). Divergence times used are shown next to the nodes: τ_*R*_ = 4*θ*_0_, τ_*S*_ = 3*θ*_0_, τ_*T*_ = 2*θ*_0_, and τ_*X*_ = τ_*Y*_ = 1.5*θ*_0_, with population sizes *θ*_0_ = 0.002 for the thin branches and *θ*_1_ = 0.01 for the thick branches. We also used a setting in which all populations on the species tree have the same size, with *θ*_0_ = *θ*_1_ = 0.01. Introgression probabilities are *φ*_*X*_ = *φ*_*Y*_ = 0.2. Data simulated under models I, O, and B are analyzed under models I, O, and B, resulting in nine combinations, with parameter estimates summarized in figures S4 (for the same population size) and S5 (for different population sizes), while results of the Bayesian test are presented in figures S6 (for the same population size) and S7 (for different population sizes).

Overall, the results parallel those for the cases of two and three species discussed above. See the SI text “Simulation in the case of four species” for detailed descriptions.

### Analysis of Heliconius genomic datasets to infer the direction of introgression

#### Overview

To assess the applicability of our results from the asymptotic analysis and computer simulation to empirical datasets and the statistical and computational feasibility of inferring the direction of gene flow using genomic sequence data, we analyzed data from *Heliconius cydno* (*C*), *H. melpomene* (*M*), and *H. hecale* (*H*) (fig. 6). Gene flow is known to occur between *H. cydno* and *H. melpomene*, whereas *H. hecale* is more distantly related, and is here treated as an outgroup, and is assumed not to have had introgression with the other two (Martin *et al*., 2013). We analyzed coding and noncoding loci on each chromosome as separate datasets (see table S4 for the numbers of loci). We fitted four models: (Ø) MSC with no gene flow, (I) MSci with *C* → *M* introgression, (O) MSci with *M* → *C* introgression, and (B) MSci with *C ⇆ M* bidirectional introgression (see fig. 1). We ran the MCMC algorithm in bpp to generate the posterior estimates of parameters in each model (Flouri *et al*., 2020) and conducted the Bayesian test of introgression (Ji *et al*., 2023). We describe the results for the coding and noncoding datasets from chromosome 1 (tables 2 & 3) in detail before discussing results for the other chromosomes.

**Table 2.** Posterior means and 95% HPD CIs for parameters in bpp analyses of two datasets of noncoding and coding loci on chromosome 1 from *Heliconius* butterflies (fig. 6) under four models with different introgression directions.

| | model ∅ (no gene flow) | Model I ($C \rightarrow M$) | Model O ($M \rightarrow C$) | Model B ($C \rightleftharpoons M$) |
| --- | --- | --- | --- | --- |
| **Noncoding loci** ($L = 5,341$ loci) | | | | |
| $\theta_H$ | 0.0131 (0.0127, 0.0136) | 0.0134 (0.0129, 0.0139) | 0.0134 (0.0129, 0.0138) | 0.0134 (0.0129, 0.0139) |
| $\theta_C$ | 0.0407 (0.0329, 0.0496) | 0.0500 (0.0274, 0.0759) | 0.0231 (0.0070, 0.0415) | 0.0499 (0.0267, 0.0759) |
| $\theta_M$ | 0.0026 (0.0021, 0.0031) | 0.0003 (0.0002, 0.0005) | 0.0001 (0.0000, 0.0002) | 0.0003 (0.0002, 0.0005) |
| $\theta_r$ | 0.0124 (0.0119, 0.0128) | 0.0123 (0.0118, 0.0127) | 0.0122 (0.0118, 0.0127) | 0.0123 (0.0118, 0.0127) |
| $\theta_s$ | 0.0343 (0.0328, 0.0358) | 0.0152 (0.0141, 0.0162) | 0.0185 (0.0175, 0.0194) | 0.0152 (0.0141, 0.0162) |
| $\theta_c$ | n/a | 0.0256 (0.0241, 0.0271) | 0.0230 (0.0206, 0.0254) | 0.0255 (0.0240, 0.0270) |
| $\theta_m$ | n/a | 0.0188 (0.0162, 0.0214) | 0.0294 (0.0262, 0.0327) | 0.0189 (0.0164, 0.0215) |
| $\tau_r$ | 0.0116 (0.0114, 0.0117) | 0.0118 (0.0116, 0.0120) | 0.0118 (0.0116, 0.0120) | 0.0118 (0.0116, 0.0120) |
| $\tau_s$ | 0.0010 (0.0008, 0.0012) | 0.0068 (0.0064, 0.0072) | 0.0051 (0.0048, 0.0053) | 0.0068 (0.0064, 0.0071) |
| $\tau_c = \tau_m$ | n/a | 0.0001 (0.0001, 0.0002) | 0.0000 (0.0000, 0.0001) | 0.0001 (0.0001, 0.0002) |
| $\varphi_c$ | n/a | n/a | 0.1744 (0.1458, 0.2038) | 0.0019 (0.0000, 0.0057) |
| $\varphi_m$ | n/a | 0.2830 (0.2565, 0.3090) | n/a | 0.2802 (0.2530, 0.3067) |
| **Coding loci** ($L = 4,942$ loci) | | | | |
| $\theta_H$ | 0.0055 (0.0053, 0.0058) | 0.0055 (0.0053, 0.0058) | 0.0055 (0.0052, 0.0057) | 0.0055 (0.0053, 0.0058) |
| $\theta_C$ | 0.0054 (0.0048, 0.0060) | 0.0361 (0.0203, 0.0545) | 0.0307 (0.0133, 0.0513) | 0.0363 (0.0204, 0.0553) |
| $\theta_M$ | 0.0016 (0.0015, 0.0018) | 0.0010 (0.0008, 0.0011) | 0.0005 (0.0003, 0.0008) | 0.0010 (0.0008, 0.0011) |
| $\theta_r$ | 0.0092 (0.0088, 0.0096) | 0.0092 (0.0088, 0.0096) | 0.0094 (0.0090, 0.0098) | 0.0092 (0.0088, 0.0096) |
| $\theta_s$ | 0.0117 (0.0111, 0.0124) | 0.0027 (0.0004, 0.0054) | 0.0092 (0.0084, 0.0100) | 0.0027 (0.0004, 0.0053) |
| $\theta_c$ | n/a | 0.0059 (0.0055, 0.0063) | 0.0044 (0.0032, 0.0055) | 0.0058 (0.0053, 0.0062) |
| $\theta_m$ | n/a | 0.0119 (0.0076, 0.0168) | 0.0105 (0.0072, 0.0144) | 0.0129 (0.0077, 0.0189) |
| $\tau_r$ | 0.0049 (0.0047, 0.0050) | 0.0049 (0.0047, 0.0050) | 0.0048 (0.0047, 0.0050) | 0.0049 (0.0047, 0.0050) |
| $\tau_s$ | 0.0009 (0.0008, 0.0010) | 0.0047 (0.0045, 0.0049) | 0.0017 (0.0015, 0.0019) | 0.0047 (0.0045, 0.0049) |
| $\tau_c = \tau_m$ | n/a | 0.0005 (0.0004, 0.0006) | 0.0002 (0.0001, 0.0003) | 0.0005 (0.0004, 0.0006) |
| $\varphi_c$ | n/a | n/a | 0.1360 (0.0783, 0.1959) | 0.0073 (0.0000, 0.0194) |
| $\varphi_m$ | n/a | 0.5119 (0.4780, 0.5451) | n/a | 0.5064 (0.4722, 0.5412) |

Note.— Results for the other chromosomes are summarized in figure S8.

**Table 3.** Bayes factors for comparing four introgression models for the *Heliconius* datasets (fig. 6, table 2), calculated using thermodynamic integration with 32 or 64 Gaussian quadrature points and Savage-Dickey density ratio with threshold ϵ = 1%, 0.1%, or 0.01%.

| $B_{ij}$ (null hypothesis tested, $H_0$) | Thermodynamic integration | | Savage-Dickey density ratio | | |
| --- | --- | --- | --- | --- | --- |
| | 32 points | 64 points | $\epsilon = 1\%$ | $\epsilon = 0.1\%$ | $\epsilon = 0.01\%$ |
| **Noncoding loci** ($L = 5,341$ loci) | | | | | |
| $B_{I\emptyset}$ ($H_0: \varphi_{C \rightarrow M} = 0$) | $e^{1087.1}$ | $e^{1082.5}$ | ∞ | ∞ | ∞ |
| $B_{O\emptyset}$ ($H_0: \varphi_{M \rightarrow C} = 0$) | $e^{946.9}$ | $e^{904.9}$ | ∞ | ∞ | ∞ |
| $B_{BI}$ ($H_0: \varphi_{M \rightarrow C} = 0$) | $e^{-5.6}$ | $e^{-9.9}$ | 0.0101 | 0.0025 | 0.0020 |
| $B_{BO}$ ($H_0: \varphi_{C \rightarrow M} = 0$) | $e^{134.6}$ | $e^{167.8}$ | ∞ | ∞ | ∞ |
| $B_{IO}$ ($H_0: \varphi_{C \rightarrow M} = 0$ vs. $\varphi_{M \rightarrow C} = 0$) | $e^{140.2}$ | $e^{177.6}$ | n/a | n/a | n/a |
| $B_{B\emptyset}$ ($H_0: \varphi_{C \rightarrow M} = 0$ and $\varphi_{M \rightarrow C} = 0$) | $e^{1081.6}$ | $e^{1072.6}$ | ∞ | ∞ | ∞ |
| **Coding loci** ($L = 4,942$ loci) | | | | | |
| $B_{I\emptyset}$ ($H_0: \varphi_{C \rightarrow M} = 0$) | $e^{359.9}$ | $e^{358.5}$ | ∞ | ∞ | ∞ |
| $B_{O\emptyset}$ ($H_0: \varphi_{M \rightarrow C} = 0$) | $e^{128.0}$ | $e^{147.6}$ | ∞ | ∞ | ∞ |
| $B_{BI}$ ($H_0: \varphi_{M \rightarrow C} = 0$) | $e^{-13.0}$ | $e^{-8.6}$ | 0.0136 | 0.0090 | 0.0073 |
| $B_{BO}$ ($H_0: \varphi_{C \rightarrow M} = 0$) | $e^{218.9}$ | $e^{202.3}$ | ∞ | ∞ | ∞ |
| $B_{IO}$ ($H_0: \varphi_{C \rightarrow M} = 0$ vs. $\varphi_{M \rightarrow C} = 0$) | $e^{231.9}$ | $e^{210.9}$ | n/a | n/a | n/a |
| $B_{B\emptyset}$ ($H_0: \varphi_{C \rightarrow M} = 0$ and $\varphi_{M \rightarrow C} = 0$) | $e^{346.8}$ | $e^{349.9}$ | ∞ | ∞ | ∞ |

Note.— The four models are ( ∅ ) MSC with no gene flow, (I) $C \rightarrow M$ introgression (I), (O) $M \rightarrow C$ introgression, and (B) $C \rightleftharpoons M$ bidirectional introgression (table 2). Bayes factor $B_{ij}$ represents the evidence in favour of model $i$ against model $j$ . We use a cut-off of 1%, so that $B_{ij} > 100$ means strong support for model $i$ and rejection of model $j$ , $B_{ij} < 0.01$ means strong support for model $j$ and rejection of model $i$ , while $0.01 < B_{ij} < 100$ means no strong preference for either model. The approach based on Savage-Dickey density ratio is inapplicable for $B_{IO}$ as models I and O are not nested. Also it produces $B = \infty$ if all values of $\varphi$ in the MCMC sample are $> \epsilon$ . Results for the other chromosomes are shown in table S5.

**Figure 6:**
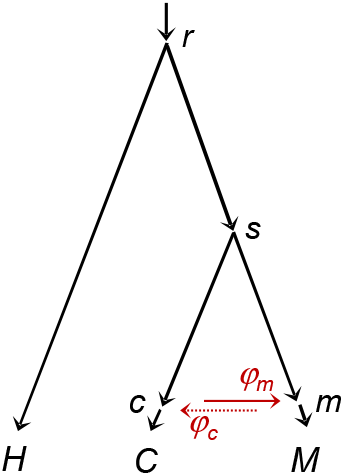
Species tree for *Heliconius hecale* (*H*), *H. cydno* (*C*), and *H. melpomene* (*M*), with introgression between *H. cydno* and *H. melpomene*, used to analyze genomic sequence data. Parameters in the MSci model include species divergence and introgression times (τ_*r*_, τ_*S*_, τ_*c*_ = τ_*m*_), population sizes for branches on the species tree (e.g., *θ*_*C*_ for branch *C* and *θ*_*c*_ for branch *Sc*), as well as introgression probabilities (*φ*_*m*_ ≡ *φ*_*C*→*M*_ and *φ*_*c*_ ≡ *φ*_*M*→*C*_). The data support the *C* → *M* introgression but not the *M* → *C* introgression, with *φ*_*m*_ > 0 and *φ*_*c*_ ≈ 0 (tables 2 and S4, fig. S8).

#### Bayesian test of introgression for chromosome 1

Results of the Bayesian test are summarized in table 3. To compare the four different models, we calculated Bayes factors using two approaches: thermodynamic integration with Gaussian quadrature (Lartillot and Philippe, 2006; Rannala and Yang, 2017) and Savage-Dickey density ratio (Ji *et al*., 2023); see Materials and Methods. The calculated values of the Bayes factor for the same test varied depending on the number of quadrature points in the thermodynamic-integration approach and on the threshold parameter in the Savage-Dickey density ratio, reflecting the challenges of calculating the marginal likelihoods or Bayes factors reliably in large datasets (Rannala and Yang, 2017). For example log *B*_Iø_ for comparison of model I (*C* → *M* introgression) against model Ø (no gene flow) was 1087.1 and 1082.5, respectively, when *K* = 32 and 64 quadrature points were used in Gaussian quadrature. This difference is mainly due to the difficulty of calculating the power posterior rather than the use of too few quadrature points (Rannala and Yang, 2017). Nevertheless, both values are far greater than the cut-off of 4.6 (= log 100). Similarly the Savage-Dickey density ratio approach estimates *B*_Iø_ to be ∞ at all three threshold values (ϵ = 1%, 0.1%, 0.01%). Both approaches thus strongly support model I with *C* → *M* introgression and reject model Ø with no gene flow.

For both datasets from chromosome 1, the two approaches to Bayes factor calculation lead to the same conclusion, as do the three threshold values for the Savage-Dickey density ratio (ϵ = 1%, 0.1%, 0.01%). The null hypothesis *φ*_*C*→*M*_ = 0 is rejected in the I-Ø and B-O comparisons, with strong support for the *C* → *M* introgression, whether or not the *M* → *C* introgression is accommodated in the model.

The B-I comparison tests the null hypothesis *φ*_*M*→*C*_ = 0 when both the null and alternative models accommodate the *C* → *M* introgression. This test leads to strong support for the null model I, with *B*_BI_ < 0.01. With *C* → *M* introgression accommodated, the data strongly support the absence of *M* → *C* introgression. Unlike Frequentist hypothesis testing, which can never support the null hypothesis strongly, here the Bayesian test strongly favors the null model I, rejecting the more general alternative model B.

However, the test of *φ*_*M*→*C*_ = 0 is significant in the O-Ø comparison when the *C* → *M* introgression is not accommodated in the null and alternative models. This result mimics our computer simulation, in which the test of gene flow is often significant if the assumed gene flow is in the wrong direction (figs. S2, S6 & S7).

Models I and O are not nested, but the Bayes factor can be used to compare them. *B*_IO_ suggests strong preference for model I (*C* → *M* gene flow) over model O (*M* → *C* gene flow).

Thus all tests have led to the same conclusions. Both the coding and noncoding datasets strongly support the presence of *H. cydno* → *H. melpomene* introgression, and both strongly support the absence of the *H. melpomene* → *H. cydno* introgression.

#### Parameter estimation for chromosome 1

Bayesian parameter estimates under the four models are summarized in table 2. Consistent with the results of the Bayesian test above, estimates of *φ* under model B suggest that gene flow is unidirectional. The estimates for the noncoding data are 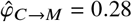 (95% HPD CI: 0.25–0.31) and 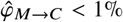 in the opposite direction, while for the coding data, they are 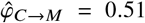 (95% HPD CI: 0.47–0.54) and 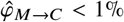 (table 2). The reasons for the higher rate 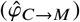 for the coding than the noncoding data are unknown. One intriguing possibility is that introgression is mostly adaptive, driven by natural selection, and that coding loci are under stronger selection. The time of introgression is nearly zero, suggesting that gene flow may be ongoing. Estimates under model I are nearly identical to those under model B. In model O where only *M* → *C* gene flow is allowed, the introgression probability is estimated to be 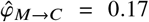 (0.15,0.20) for the noncoding data, and 0.14 (0.08, 0.20) for the coding data. Those rates are substantial, consistent with the significant test results (*B*_OØ_). Even if gene flow is unidirectional from *C* to *M*, assuming introgression in the opposite (and presumably wrong) direction leads to high estimates of the rate and significant test results. Those results again parallel our simulations (figs. S2, S6 & S7). The misspecified introgression direction in model O causes large estimates of *θ*_*S*_ and reduces τ_*S*_. Those results mimic the behaviors of the misspecified model in the large→small case in our theoretical analysis and simulations (fig. 3, table S1**d** large→small).

We note that the divergence time between *H. cydno* and *H. melpomene* (τ_*S*_) is estimated to be much smaller, and *θ*_*S*_ is much larger under model Ø (no gene flow) than under models I or B. This is because ignoring gene flow when it occurs causes model Ø to misinterpret reduced between-species sequence divergence (due to introgression) as more recent species divergence (Leaché *et al*., 2014; Tiley *et al*., 2023).

#### Parameter estimation for the other autosomes

we analyzed the coding and noncoding data from all chromosomes in the same way, with parameter estimates under the four models (Ø, I, O, B) summarized in figure S8 (see also table S5), while Bayesian test results are in table S6.

There is overall consistency among the autosomes (chromosomes 1-20), although estimates of some parameters from chromosomes 5, 10, 13, 15, and 19 appear as outliers. For example, estimates of *θ*_*C*_ and *θ*_*M*_ are unusually large for chromosomes 5, 15, 19, and 20. A likely explanation is that some individuals are partially inbred, with large variations in heterozygosity across chromosomes. We discuss results for the autosomes first before dealing with chromosome 21 (the Z chromosome).

For the autosomes, there is overall consistency between the coding and noncoding data: divergence times τ_*r*_ and τ_*S*_ and population sizes *θ*_*H*_ and *θ*_*r*_ are larger for the noncoding than coding data, by a similar factor across chromosomes (fig. S8). This can be explained by a reduced effective neutral mutation rate for the coding data, due to purifying selection removing nonsynonymous mutations.

Although model Ø (no gene flow) underestimated the divergence time between the two species involved in gene flow, τ_*S*_ (see above), all four models including model Ø produce nearly identical estimates of τ_*r*_, indicating that the impact of introgression is local on the species tree, only affecting estimates of parameters for nodes close to the introgression event. Estimates of τ_*S*_ under model O are consistently smaller than under models I and B, especially for the coding data, apparently related to the low estimates of *φ*_*M*→*C*_ for the coding data under model O. Introgression time τ_*c*_ = τ_*m*_ is nearly zero for most chromosomes under models I, O, and B, indicating that gene flow may be ongoing (Huang *et al*., 2022).

Estimates of introgression probability *φ*_*C*→*M*_ are very similar between models I and B, and they are consistently larger for coding than noncoding data. Estimates of *φ*_*M*→*C*_ under model B are consistently ≈ 0, suggesting the absence of M → C gene flow. Estimates of *φ*_*M*→*C*_ under model O, assuming introgression in the wrong direction, are always larger than estimates under model B, but vary among chromosomes. These results are consistent with our simulations (e.g., figs. 3, cases **a**–**d**), where estimates of introgression probability *φ*_*X*_ in model O vary, even though the true rate in the opposite direction is fixed (*φ*_*Y*_ = 0.2), influenced by estimates of population sizes such as *θ*_*X*_ and *θ*_*Y*_.

#### Bayesian test of introgression for the autosomes

Bayes factors calculated via the Savage-Dickey density ratio are presented in table S6. The results are similar to those for chromosome 1, with overwhelming evidence for the C → M introgression and no evidence for M → C introgression. For some datasets, *B*_OØ_ < 100, so that the test of gene flow (*H*_0_ : *φ*_*M*→*C*_ = 0) is not significant when introgression was assumed to be in the wrong direction.

#### Unidentifiability issues for the haploid sex chromosome

Results for chromosome 21 (the Z chromosome) show very different patterns from the autosomes (fig. S8), because we have only one haploid sequence per species in the data: both *H. cydno* and *H. melpomene* samples are hemizygous females, i.e ZW. For such data, some parameters are unidentifiable in any of the four models, such as *θ*_*C*_, *θ*_*M*_, *θ*_*H*_ for the extant species. As discussed before, models I and O are unidentifiable, with the parameter mapping 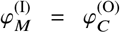 and 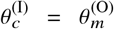

Thus those parameters should have exactly the same posterior. This is a case of cross-model unidentifiability. Model B applied to data from the Z chromosome (with one sequence per species per locus) poses an even more complex unidentifiability issue. As discussed later in Discussion, there are four unidentifiable towers in the posterior surface (fig. 7**b**). Due to the symmetry of the posterior surface, the marginal posteriors for *φ*_*M*_ and *φ*_*C*_ are identical, as are the posteriors for *φ*_*M*_ and 1−*φ*_*M*_; as a result, the posterior means of *φ*_*M*_ and *φ*_*C*_ are both 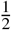 (fig. S8). Similarly the posteriors for *θ*_*c*_ and *θ*_*m*_ are identical. Nevertheless, parameters not involved in the unidentifiability (such as τ_*r*_, τ_*c*_, *θ*_*r*_) are well estimated. In theory, the four towers represent unidentifiability of the label-switching type, and a relabeling algorithm can be used to process the MCMC samples to map the parameter values onto one of the four towers, as in Yang and Flouri (2022). This is not pursued here. Instead our objective here is to provide an explanations for the results of figure S8 (chromosome 21, model B). We recommend that multiple samples per species per locus (in particular from the recipient species) should be used to estimate introgression probabilities. Note that one diploid individual is equivalent to two haploid sequences.

**Figure 7:**
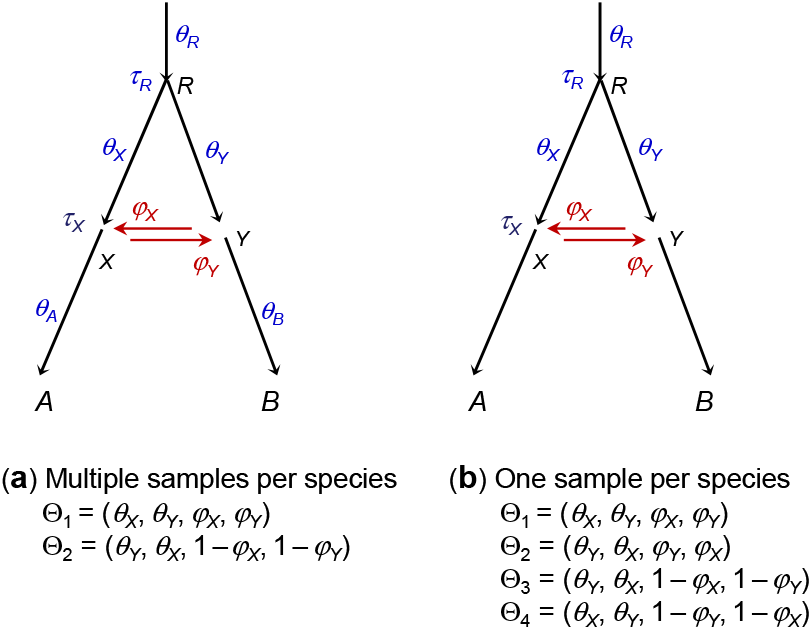
(**a**) When multiple sequences are sampled per species per locus, the MSci model with bidirectional introgression between sister lineages has two unidentifiable modes in the posterior (Θ_1_, Θ_2_) (Yang and Flouri, 2022). Population size parameters for extant species (*θ*_*A*_, *θ*_*B*_) are identifiable. (**b**) When one sequence is sampled per species per locus, the same model shows four unidentifiable modes (Θ_1_, Θ_2_, Θ_3_, Θ_4_). Also *θ*_*A*_ and *θ*_*B*_ are unidentifiable and are not parameters in the model.

## Discussion

### Unidentifiability of introgression models

Here we review the different types of unidentifiability issues encountered in this study. See Yang and Flouri (2022) for detailed discussions.

Yang and Flouri (2022) distinguished between *within-model* and *cross-model* unidentifiability. If the probability distributions of the data are identical under model *m* with parameters Θ and under model *m*′ with parameters Θ ′, with

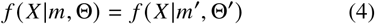

for all possible data *X*, then data *X* cannot identify (*m*, Θ) and (*m′*, Θ *′*). If *m = m′* and Θ ≠ Θ ′, the parameters within the given model are unidentifiable. If *m* ≠ *m′*, the two models are unidentifiable (cross-model); in this case there is a parameter mapping from Θ in *m* to Θ *′* in *m′*.

In the case of two species (say *A* and *B*) with one sequence sampled per species per locus, the coalescent time (*t*_*ab*_) between the two sequences (*a, b*) has the same distribution under model I with *A* → *B* introgression and under model O with *B* → *A* introgression (Appendix A). As a result, the two models are unidentifiable, or in other words, the introgression direction is unidentifiable (Yang and Flouri, 2022, fig. 10). This is a case of cross-model unidentifiability. The parameter mapping is 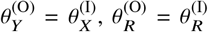 and *φ*_*X*_ = *φ*_*Y*_, with τ_*R*_ and τ_*X*_ being identical between the two models (eq. A3, fig. 1). In the analysis of chromosome 21 from the *Heliconius*, model I and model O are unidentifiable, with 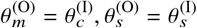 and *φ*_*c*_ = *φ*_*m*_ (fig. S8, table S5).

If the model (model I, say) is given, parameters τ_*R*_, τ_*X*_, *θ*_*X*_, and *φ*_*Y*_ are identifiable even with data of one sequence per species per locus. In the example of chromosome 21 for the *Heliconius* data, parameters τ_*S*_, τ_*c*_, *θ*_*c*_, and *φ*_*m*_ are identifiable (fig. S8, table S5).

In the case of two species, introgression direction becomes identifiable if multiple sequences are sampled per species per locus (Yang and Flouri, 2022). Furthermore if data from other species are available and if gene flow occurs between nonsister species, introgression direction affects the distributions of the gene trees and coalescent times, and is identifiable whether one sequence or multiple sequences are sampled per species per locus (Jiao *et al*., 2021; Hibbins and Hahn, 2022; Yang and Flouri, 2022).

Furthermore, bidirectional introgression (BDI) models have *unidentifiability of the label-switching type* (Yang and Flouri, 2022). The situation is similar to label switching in clustering analysis. Let the parameters in the model be Θ = (*P*_1_, *µ*_1_, *µ*_2_), meaning that the two groups are in proportions *P*_1_ and *P*_2_ = 1−*P*_1_ with means *µ*_1_ and *µ*_2_. Then Θ and Θ ′ = (*P′*_1_, µ′_1_, µ′_2_) = (*P*_2_, µ_2_, µ_1_) are unidentifiable as their only difference is in the labels ‘1’ and ‘2’ for the two groups. Models with unidentifiability of the label-switching type can still be used in inference. If multiple samples are available per species per locus, the BDI model with introgression between sister lineages shows two unidentifiable towers involving the two introgression probabilities and two population size parameters, as discussed by Yang and Flouri (2022): in figure 7**a**, Θ_1_ = (*θ*_*X*_, *θ*_*Y*_, *φ*_*X*_, *φ*_*Y*_) and Θ_2_ = (*θ*_*X*_, *θ*_*Y*_, *φ*_*X*_, *φ*_*Y*_) are unidentifiable. This is a within-model unidentifiability of the label-switching type.

For chromosome 21 in the *Heliconius* genomic data, only one (haploid) sequence is sampled per species per locus (fig. S8). Unidentifiability issues in analysis of such data under the BDI model were not considered by Yang and Flouri (2022). We note that with one sequence per species per locus the BDI model with introgression between sister lineages shows four unidentifiable modes or towers in the posterior: in figure 7**b**, Θ_1_ = (*θ*_*X*_, *θ*_*Y*_, *φ*_*X*_, *φ*_*Y*_), Θ_2_ = (*θ*_*Y*_, *θ*_*X*_, *φ*_*Y*_, *φ*_*X*_), Θ_3_ = (*θ*_*Y*_, *θ*_*X*_, 1 − *φ*_*X*_, 1 − *φ*_*Y*_), and Θ_4_ = (*θ*_*X*_, *θ*_*Y*_, 1 − *φ*_*Y*_, 1 − *φ*_*X*_) are unidentifiable (fig. 7). If introgression is between nonsister lineages, each BDI will create two cross-model towers, whether one sequence or multiple sequences are sampled per species per locus (Yang and Flouri, 2022).

### Asymmetry of gene flow in nature

No systematic studies have examined the frequency of unidirectional versus bidirectional gene flow given that two species are involved in introgression or hybridization. Both scenarios appear to be common. Sometimes gene flow occurs in one direction even though opportunities exist also in the opposite direction. A well-documented example is gene flow in the *Anopheles gambiae* group of mosquitoes in sub-Saharan Africa (della Torre *et al*., 1997; Slotman *et al*., 2005b). Analysis of genomic data provides strong evidence for gene flow from *A. arabiensis* to *A. gambiae* or its sister species *A. coluzzii*, while the rate of gene flow in the opposite direction was estimated to be 0 (Thawornwattana *et al*., 2018; Flouri *et al*., 2020). This result from comparisons of genomic sequences is consistent with crossing experiments which supported introgression of autosomal regions from *A. arabiensis* into *A. coluzzii* but not in the opposite direction (della Torre *et al*., 1997; Slotman *et al*., 2005b). One possible explanation is that the X chromosome from one species may be incompatible with the autosomal background of the other species (Slotman *et al*., 2004, 2005a). The introgression from *A. arabiensis* into the common ancestor of *A. gambiae* and *A. coluzzii* has been hypothesized to have facilitated the range expansion of *A. gambiae* and *A. coluzzii* into the more arid savanna habitats of *A. arabiensis* (Coluzzi *et al*., 1979; Ayala and Coluzzi, 2005).

Note that the rate of gene flow in the MSci model estimated from the genomic sequence data is an ‘effective’ rate, reflecting the combined effects of gene flow and natural selection. Most introgressed alleles are expected to be purged in the recipient species because of incompatibilities with the host genomic background. It seems likely that alleles at introgressed loci from species *A* on the genomic background of species *B* will have different fitnesses than introgressed alleles from *B* on the background of *A*. Another factor is geographic context. If a smaller population of species *A* hybridizes with a larger population of species *B, A* is more likely to be swamped by *B*, making introgression asymmetrical. With all those factors considered, one should expect gene flow to be asymmetrical in most systems, with different rates in the two directions.

### Gene flow in *Heliconius* butterflies

*Heliconius cydno* and *H. melpomene* are broadly sympatric across Central America and northwestern South America, and are known to hybridize in the wild (Mallet *et al*., 2007). Our analysis supports recent unidirectional gene flow from *H. cydno* into *H. melpomene* (fig. 6, tables 2–3, S5, S6), in Panama, where *H. cydno chioneus* and *H. melpomene rosina* are broadly sympatric. In captivity, male F_1_ hybrids are fertile while female F_1_ hybrids are sterile; male hybrids backcross to either parental species much more readily than the pure species mate with one another (Naisbit *et al*., 2001, 2002).

Previous studies used different approaches to estimate gene flow between these two species. Early phylogenetic analyses of multilocus data attributed recent gene flow between *H. cydno chioneus* and *H. melpomene rosina* as a cause for gene tree variation among loci (Beltrán *et al*., 2002). An isolation-with-migration (IM) analysis (Hey and Nielsen, 2004) using a small number of loci yielded an estimated symmetric bidirectional migration rate *m* between the two species of 1.7 × 10^−6^ (95% CI 1.0–45 ×10^−6^) per generation, with *H. cydno chioneus* having a larger effective population size (Bull *et al*., 2006). An IM model allowing for different migration rates in each direction found evidence for unidirectional gene flow from *H. cydno* into *H. melpomene*, with 2*N*_*M*_*m*_*C*→*M*_ = 0.294 (90% HPD CI: 0.116–0.737) whereas 2*N*_*C*_*m*_*M*→*C*_ = 0.000 (0.000, 0.454) (Kronforst *et al*., 2006), consistent with our results. Similar patterns were obtained in a subsequent IMa2 analysis (Hey, 2010) of a larger dataset (Kronforst *et al*., 2013). In a more recent analysis of genome-scale data, Martin *et al*. (2015) estimated a symmetric bidirectional migration rate between *H. c. chioneus* and *H. m. rosina* to be 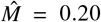 (90% HPD interval: 0.09–0.40) per generation. Lohse *et al*. (2016) compared three models: complete isolation after divergence, and two IM models with unidirectional gene flow, and preferred the model with gene flow from *H. cydno* into *H. melpomene rosina*, with estimated migration rate 4*Nm* = 1.5. Martin *et al*. (2019) used gene tree frequencies to suggest extensive gene flow from *H. cydno* into *H. melpomene* in Panama.

Our estimates are in general consistent among chromosomes and between coding and noncoding data. However, only a few samples per species are included in the genomic data, and some individuals are from inbred lines (selected for sequencing because of easy assembly). These features of the data may have affected our estimates, and may account for the outlier estimates observed for a few chromosomes (fig. S8). Overall, our analyses of genomic data are consistent with previous estimates.

We note that the null model Ø in the Bayesian test used in this study constrains the population sizes (*θ*_*C*_ = *θ*_*c*_ and *θ*_*M*_ = *θ*_*m*_) as well as the introgression probability (*φ* = 0), compared with the alternative model (models I, O, or B) (Ji *et al*., 2023). Rejection of the null model may in theory be due to either introgression or inequality of population sizes, or both. A sharper test may use an alternative model with the same constraints on the population sizes as in the null model (*θ*_*C*_ = *θ*_*c*_, *θ*_*M*_ = *θ*_*m*_) so that the two models under comparison has the only difference concerning the introgression probability (*φ* = 0 vs. *φ* > 0); this is test 2 in Ji *et al*. (2023, fig. 3). For the *Heliconius* data, we note that the CIs for *φ*_*c*_ excludes the null value *φ*_*c*_ = 0 for every autosome (fig. S8), providing strong evidence for introgression. We leave it to future work to use more genomic data and more focused test to infer gene flow in this group of *Heliconius* butterflies.

### Inferring the direction of gene flow using genomic data

Our asymptotic analysis, computer simulation and real data analysis have produced highly consistent results. We have illustrated that one may gain much insight into the workings of likelihood-based inference under the MSci model by simply considering the coalescent times for sequence pairs (*t*_*aa*_, *t*_*ab*_, *t*_*bb*_) even though these may appear to be naively simple summaries of the original data of multilocus sequence alignments (table 1). Knowledge of important features in the data that drive the estimation of model parameters, such as the introgression time and introgression probability, is very useful when we interpret results from analysis of real datasets. For example, we found that if introgression is assumed to occur in the wrong direction, Bayesian test of gene flow will often be significant, and Bayesian estimates of introgression rate will typically be non-zero and may even be greater than the true rate in the correct direction. Thus neither a significant test nor a high rate estimate is reliable evidence that introgression occurred in the specified direction. This result may seem surprising and disturbing given that introgression in the specified direction is non-existent. Our analyses of both simulated and real data suggest that application of the bidirectional model may be advisable if there is uncertainty concerning the direction of gene flow. If gene flow is truly unidirectional, over-parametrization of the bidirectional model appear to incur little cost in statistical performance even though it does add to computational cost: posterior CIs and power to detect gene flow under the bidirectional model are very similar to those under the true unidirectional model.

In the future, we hope to implement efficient cross-model MCMC algorithms to search for the best MSci model in the model space. Currently MCMC algorithms that move between models (Zhang *et al*., 2018; Wen and Nakhleh, 2018) propose changes to the MSci model when the gene trees at all loci are fixed. Such algorithms have poor mixing properties as the proposed new model is very likely to be in conflict with at least some gene trees, leading to rejection of the proposal. The algorithms do not appear to be feasible for analyzing even small datasets with 100 loci (Zhang *et al*., 2018; Wen and Nakhleh, 2018). However, thousands of loci are often needed to provide precise and reliable inference of introgression between species. Smart MCMC moves that make coordinated changes to the gene trees when the chain moves from one model to another, similar to the algorithms developed under the MSC model with no gene flow for updating species divergence times (the rubber-band algorithm, Rannala and Yang, 2003) or species phylogenies (the species-tree NNI or SPR moves, Yang and Rannala, 2014; Rannala and Yang, 2017) may offer significant improvements even though they are more challenging to develop. Similarly most approximate methods for detecting gene flow are based on species triplets or quartets and use summaries of sequence data such as genome-wide site-pattern counts (e.g., the *D*-statistic, Green *et al*., 2010; Durand *et al*., 2011 and HyDe, Blischak *et al*., 2018) or gene-tree frequencies (as in SNaQ, Solis-Lemus and Ane, 2016). The triplet or quartet trees are then assembled into a large species tree with introgression events (Solis-Lemus *et al*., 2017). The maximum likelihood method of Yu *et al*. (2014) uses estimated gene trees from multiple loci as input data to infer the introgression model, but does not utilize information in coalescent times or account for uncertainties in estimated gene-tree topologies. These approximate methods do not make an efficient use of information in the multilocus sequence data, and offer ample chances for improvements (Degnan, 2018; Elworth *et al*., 2019; Jiao *et al*., 2021).

Lastly, likelihood implementations under a fixed MSci model already offer powerful tools for inferring gene flow using genomic sequence data (Flouri *et al*., 2020). Our analyses of both simulated and real data have demonstrated that typical genomic datasets may be very informative about the direction, timing and strength of interspecific introgression, and that current Bayesian implementations of the MSci model can accommodate thousands of genomic loci to allow precise and accurate estimation of introgression times and introgression probabilities (Thawornwattana *et al*., 2022; Ji *et al*., 2023).

## Materials and Methods

### Asymptotic analysis and simulation in the case of two species

We examined the distributions of coalescent times and conducted computer simulations under model I of figure 1**a**, with *A* → *B* introgression. We used four sets of parameter values.

a. same *θ* tall tree: all populations have the same size with *θ* = 0.01. The other parameters are τ_*R*_ = *θ*, τ_*X*_ = 0.5*θ*, and *φ*_*Y*_ = 0.2.
b. same *θ* short tree: *θ* = 0.01 for all populations, τ_*R*_ = 0.5*θ*, τ_*X*_ = 0.25*θ*, and *φ*_*Y*_ = 0.2.
c. small to large: different species on the species tree have different population sizes, with *θ*_*A*_ = *θ*_*X*_ = *θ*_*R*_ = *θ*_0_ = 0.002 on the left of the tree and *θ*_*B*_ = *θ*_*Y*_ = *θ*_1_ = 0.01 on the right, with introgression from a small population to a large one (fig. 1**a**). Other parameters are τ_*R*_ = 3*θ*_0_, τ_*X*_ = 1.5*θ*_0_ and *φ*_*Y*_ = 0.2.
d. large to small: This is the same as case (**c**) except that *θ*_*A*_ = *θ*_*X*_ = *θ*_*R*_ = *θ*_0_ = 0.01 on the left of the tree and *θ*_*B*_ = *θ*_*Y*_ = *θ*_1_ = 0.002 on the right, so that introgression is from a large population to a small one.

We simulated multilocus sequence datasets under model I (fig. 1**a**) and analyzed them under models I, O, and B (fig. 1**a**–**c**). Each replicate dataset consists of *L* = 250, 1000 or 4000 loci, with *n* = 4 sequences sampled per species per locus. The sequence length is *N* = 500 sites. The simulate option of bpp (Flouri *et al*., 2018) was used to simulate gene trees with coalescent times and to ‘evolve’ sequences along the gene tree under the JC model (Jukes and Cantor, 1969). Sequences at the tips of the gene tree constitute the data. The number of replicates was 100.

Each replicate dataset was then analyzed using bpp (Flouri *et al*., 2018, 2020) under models I, O, and B of figure 1**a**–**c**. This setting in which the model is fixed corresponds to the A00 analysis of (Yang, 2015). The JC model was assumed in the analysis. Gamma priors were assigned to the age of the root of the species tree (τ_*R*_) and to population size parameters (*θ*), with the shape parameter *α* = 2 so that the prior was diffuse and with the rate parameter *β* chosen so that the prior mean was close to the true values. We used τ_*R*_ ∼ G(2, 200) and *θ* ∼ G(2, 200) for case **a** “same *θ* tall tree”; τ_*R*_ ∼ G(2, 400) and *θ* ∼ G(2, 200) for case **b** “same *θ* short tree”; τ_*R*_ ∼ G(2, 400) and *θ* ∼ G(2, 400) for case **c** “small to large” and **d** “large to small”. Introgression probability *φ* was assigned the beta prior beta(1, 1), which is 𝕌(0, 1).

MCMC settings were chosen by performing pilot runs, with MCMC convergence assessed by verifying consistency between replicate runs for the same analysis. The same setting was then used to analyze all replicate datasets. We used 16,000 MCMC iterations as burnin, and then took 10^5^ samples, sampling every 2 iterations. Running time for analyzing one replicate dataset was ∼ 45mins for *L* = 250 loci or ∼ 3hrs for *L* = 1000 using one thread, and ∼ 12hrs for *L* = 4000 using two threads.

### Simulation to evaluate the gain in information for estimating *φ* by adding a third species

Given the introgression model for two species (*A, B*) of figure 1**a**, with *A* → *B* introgression, we added a third species (*C*) and assessed the gain in information for estimating *φ*. There are five branches on the two-species tree, to which the third species could be attached (fig. 4**a**–**e**): (**a**) the root population, (**b, c**) the source and target populations before gene flow, and (**d, e**) the source and target populations after gene flow. In all cases *φ* = 0.2. The original two-species tree had τ_*R*_ = *θ*_1_ and τ_*X*_ = *θ*_1_/2. In cases **b**–**e**, species *C* was attached to the midpoint of the target branch, while in **a**, the new root was 1.25× as old as the old root. For models **a, d & e**, all populations on the species tree had the same size, with *θ*_1_ = 0.01. For cases **b** and **c**, three scenarios are considered: (i) equal population size, with *θ*_1_ = 0.01 for all populations; (ii) from small to large, with *θ*_*A*_ = *θ*_*X*_ = *θ*_*S*_ = *θ*_0_ = 0.002 for the thin branches in case **b** and *θ*_*A*_ = *θ*_*X*_ = *θ*_0_ = 0.002 in case **c** and with *θ*_1_ = 0.01 for all other branches; and (iii) from large to small, with *θ*_*B*_ = *θ*_*Y*_ = *θ*_0_ = 0.002 in case **b** and *θ*_*B*_ = *θ*_*Y*_ = *θ*_*S*_ = *θ*_0_ = 0.002 in case **c** and with *θ*_1_ = 0.01 for all other branches. For each parameter setting, we simulated 100 replicate datesets. Each dataset consisted of *L* = 1000 loci, with *n*_*A*_ = *n*_*B*_ = 4 sequences per species per locus and *N* = 500 sites in the sequence. Each dataset was analyzed using bpp to estimate the parameters in the MSci model (fig. 4**a**–**e**). Gamma priors were assigned to τ_*R*_ and *θ*: τ_*R*_ ∼ G(2, 200) and *θ* ∼ G(2, 200), while *φ*_*A*→*B*_ ∼ 𝕌(0, 1). We used 32,000 MCMC iterations as burnin, and then took 10^6^ samples, sampling every 10 iterations. Running time for analyzing one dataset using one thread was ∼30 hrs.

### Simulation in the case of four species: inflow versus outflow

We simulated data under the three MSci models (I, O, B) of figure 5**a**-**c**, with introgression between nonsister species *A* and *B* on a four-species tree ((*A*, (*B, C*)), *D*). The three models differ in the assumed direction of gene flow, with I for inflow from *A* to *B*, O for outflow from *B* to *A*, and B for bidirectional introgression between *A* and *B*. We used two sets of parameter values. In the first set (same-*θ*), all species on the tree had the same population size, with *θ*_0_ = *θ*_1_ = 0.01. In the second set (different-*θ*), the thin branches had *θ*_0_ = 0.002 while the thick branches had *θ*_1_ = 0.01 (fig. 5**a**-**c**). Other parameters were the same in the two settings, with τ_*R*_ = 4*θ*_0_, τ_*S*_ = 3*θ*_0_, τ_*T*_ = 2*θ*_0_, and τ_*X*_ = τ_*Y*_ = 1.5*θ*_0_, and the introgression probabilities were *φ*_*X*_ = *φ*_*Y*_ = 0.2.

Each dataset consists of *L* = 250, 1000, or 4000 loci, with *n* = 4 sequences per species per locus and with *N* = 500 sites in the sequence. The number of replicates was 100. With three MSci models (I, O, B), two population-size settings (same-*θ* vs. different-*θ*), and three data sizes (*L*), a total of 3 × 2 × 3 × 100 = 1800 datasets were generated. Each dataset was analyzed under the three models (I, O, B). Gamma priors were assigned to τ_*R*_ and *θ*: τ_*R*_ ∼ G(2, 200) and *θ* ∼ G(2, 400), while *φ* ∼ 𝕌(0, 1). We used 32,000 MCMC iterations as burnin, and took 2 × 10^5^ samples, sampling every 5 iterations. Running time for analyzing one dataset was ∼12hrs for small datasets of *L* = 250 loci and 60hrs for *L* = 1000 using one thread, and ∼120hrs for *L* = 4000 using two threads.

### Analysis of the *Heliconius* butterfly dataset

We processed the raw genomic sequencing data of Edelman *et al*. (2019) from three species of *Heliconius* butterflies, *H. hecale* (*H*), *H. cydno* (*C*), and *H. melpomene* (*M*), to retrieve coding and noncoding loci for each chromosome, following the procedure of Thawornwattana *et al*. (2022). See table S4 for the number of loci in each of the 22 datasets. Each locus consisted of one unphased diploid sequence per species, except the Z chromosome 21 for which only a haploid sequence is available per species (from ZW females). Heterozygote phase in the diploid sequence was resolved using an analytical integration algorithm in the likelihood calculation in bpp (Gronau *et al*., 2011; Flouri *et al*., 2018; Huang *et al*., 2022). We fitted four MSci models with different introgression directions: (Ø) MSC with no gene flow, (I) *C* → *M* introgression, (O) *M* → *C* introgression, and (B) *C ⇆ M* bidirectional introgression.

We assigned priors τ_*r*_ ∼ G(4, 200), *θ* ∼ G(2, 200), and *φ* ∼ 𝕌(0, 1). We used 10^5^ MCMC iterations for burnin, and recorded 10^4^ samples, sampling every 100 iterations. For each model, we performed ten independent runs to confirm consistency between runs. The resulting MCMC samples were combined to produce final posterior estimates. Each run took ∼100hrs.

### Bayesian test of introgression

We applied the Bayesian test of introgression (Ji *et al*., 2023) to data for two species simulated under the models of figure 1**a**-**c**, the data for four species simulated under models I, O, and B of figure 5, and the *Heliconius* datasets (fig. 6).

Bayesian model selection is used to compare the null model of no gene flow *H*_0_ : *φ* = 0 and the alternative model of introgression *H*_1_ : *φ* > 0. The Bayes factor 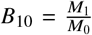, where *M*_0_ and *M*_1_ are marginal likelihood values under *H*_0_ and *H*_1_. If the prior model probabilities are π_0_ and π_1_, *B*_10_ can be converted into posterior model probabilities as 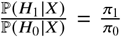 *B*_10_. If π_0_ = π_1_, *B*_10_ = 100 will translate to the posterior probability ℙ(*H*_0_| *X*) ≈ 1%. Thus *B*_10_ > 100 may be considered strong evidence in support of *H*_1_ over *H*_0_, while *B*_10_ < 0.01 is strong evidence in favor of *H*_0_ over *H*_1_.

As *H*_0_ and *H*_1_ are nested, *B*_10_ can be calculated using the Savage-Dickey density ratio (Dickey, 1971), by using an MCMC sample under *H*_1_ (Ji *et al*., 2023). Define an interval of null effects, ø: *φ* < ϵ, inside which the introgression probability is so small that introgression may be considered nonexistent. The Bayes factor in favor of *H*_1_ over *H*_0_ is then

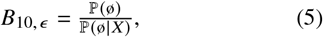

where ℙ(ø) is the prior probability of the null interval, while ℙ(ø|*X*) is the posterior probability, both calculated under *H*_1_ (Ji *et al*., 2023). Note that ℙ(ø) = ℙ(*φ* < ϵ) = ϵ if the prior is *φ* ∼ 𝕌(0, 1). When ϵ → 0, *B*_10,ϵ_ → *B*_10_ (Ji *et al*., 2023). We used a few values for ϵ in the range 0.01%–1% to assess its effect. This approach has a computational advantage as it requires running the MCMC under *H*_1_ only and avoids trans-model MCMC algorithms or calculation of marginal likelihood values.

For the *Heliconius* datasets, we in addition used thermodynamic integration combined with Gaussian quadrature to calculate the marginal likelihood under each model, using 32 or 64 quadrature points (Lartillot and Philippe, 2006; Rannala and Yang, 2017). This approach applies even if the compared models are non-nested, and was used to conduct pairwise comparisons among all four models fitted to the *Heliconius* data.

## Supplementary Material

Supplementary data are available at Molecular Biology and Evolution online.

## Acknowledgments

We thank two reviewers for many suggestions and constructive criticisms. This study is supported by Biotechnology and Biological Sciences Research Council grants (BB/T003502/1, BB/R01356X/1) to Z.Y., start-up funds from Harvard University to J.M., a studentship from the Organismic and Evolutionary Biology Department, Harvard University, to Y.T., and a Natural Science Foundation of China grant (32200490) to J.H.

## Appendix A. The distribution of coalescent times under the MSci model for two species

Here we gave the probability densities of coalescent times (*t*_*aa*_, *t*_*ab*_, *t*_*bb*_) between two sequences sampled from species *A* and *B* under the MSci models I, O, and B of figure 1**a**–**c**. These are simple cases of the gene-tree densities given by, e.g., Yu *et al*. (2014; see also Lohse and Frantz, 2014). Example densities under models I and O are plotted in figure 2 for four sets of parameter values.

Under model I,

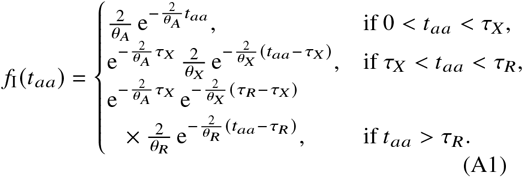

This is a function of τ_*R*_, τ_*X*_, *θ*_*A*_, *θ*_*X*_, *θ*_*R*_, independent of *θ*_*B*_, *θ*_*Y*_, *φ*_*Y*_. From the viewpoint of the two *A* sequences, there are demographic changes in population size with *θ*_*A*_, *θ*_*X*_, and *θ*_*R*_, respectively, for the three time segments (0, τ_*X*_), (τ_*X*_, τ_*R*_), and (τ_*R*_, ∞).

The coalescent time between two sequences sampled from species *B* has the distribution

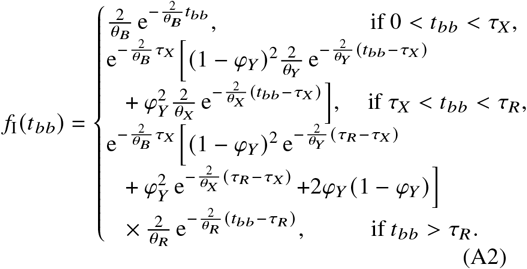

This is a function of τ_*R*_, τ_*X*_, *θ*_*B*_, *θ*_*X*_, *θ*_*Y*_, *θ*_*R*_, *φ*_*Y*_, and is independent of *θ*_*A*_. In the time interval (0, τ_*X*_), the two *B* sequences coalesce at the rate 2/*θ*_*B*_, as in the case of no gene flow. Coalescence during the time interval τ_*X*_ < *t*_*bb*_ < τ_*R*_ can occur in either *X* or *Y*. The former occurs if both *B* sequences migrate to *X* (which occurs with probability 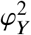) and then coalesce in *X* at the rate 2/*θ*_*X*_, whereas the latter occurs when both *B* sequences fail to migrate and thus stay in *Y* (with probability (1 − *φ*_*Y*_)^2^) and then coalesce in *Y* at the rate 2/*θ*_*Y*_. If one of the *B* sequences migrates to *X* and the other stays in *Y*, coalescence will be impossible, resulting in a suppression of coalescent events in this time interval (see fig. 2 for *f* (*t*_*bb*_)). If the two *B* sequences do not coalesce in *B*, and they do not coalesce in either *X* or *Y*, they will coalesce in species *R* (with *t*_*bb*_ > τ_*R*_), at the rate 2/*θ*_*R*_.

Finally

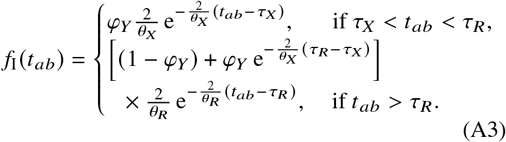

This is a function of τ_*R*_, τ_*X*_, *θ*_*X*_, *θ*_*R*_, and *φ*_*Y*_, and is independent of *θ*_*A*_, *θ*_*B*_, *θ*_*Y*_. Coalescence between *a* and *b* may occur during (τ_*X*_, τ_*R*_) at the rate 2/*θ*_*X*_ if the *B* sequence migrates into *X* (with probability *φ*_*Y*_).

Under model O with *B* → *A* introgression (fig. 1**b**), *f*_O_(*t*_*aa*_) and *f*_O_(*t*_*bb*_) are given by *f*_I_(*t*_*bb*_) and *f*_I_(*t*_*aa*_) with a change of symbols. In particular, 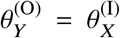 and *φ*_*X*_ = *φ*_*Y*_, with τ_*R*_, τ_*X*_, *θ*_*R*_ being identical between the two models.

Under model B with both *A* → *B* and *B* → *A* introgressions (fig. 1**c**), *f*_B_(*t*_*aa*_) = *f*_O_(*t*_*aa*_) and *f*_B_(*t*_*bb*_) = *f*_I_(*t*_*bb*_), while

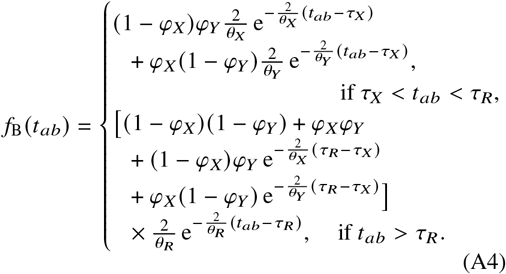

## References

Arnold, M. L. and Kunte, K. 2017. Adaptive genetic exchange: A tangled history of admixture and evolutionary innovation. Trends Ecol. Evol., 32(8): 601–611.

Ayala, F. J. and Coluzzi, M. 2005. Chromosome speciation: Humans, Drosophila, and mosquitoes. Proc. Natl. Acad. Sci. USA, 102(suppl 1): 6535–6542.

Barton, N. and Bengtsson, B. O. 1986. The barrier to genetic exchange between hybridising populations. Heredity, 57(3): 357–376.

Beltrán, M., Jiggins, C. D., Bull, V., Linares, M., Mallet, J., McMillan, W. O., and Bermingham, E. 2002. Phylogenetic discordance at the species boundary: Comparative gene genealogies among rapidly radiating Heliconius butterflies. Mol. Biol. Evol., 19(12): 2176–2190.

Blischak, P. D., Chifman, J., Wolfe, A. D., and Kubatko, L. S. 2018. HyDe: A Python package for genome-scale hybridization detection. Syst. Biol., 67(5): 821–829.

Bull, V., Beltrán, M., Jiggins, C. D., McMillan, W. O., Bermingham, E., and Mallet, J. 2006. Polyphyly and gene flow between non-sibling Heliconius species. BMC Biol., 4(1): 11.

Campbell, C. R., Poelstra, J. W., and Yoder, A. D. 2018. What is speciation genomics? The roles of ecology, gene flow, and genomic architecture in the formation of species. Biol. J. Linn. Soc., 124(4): 561–583.

Coluzzi, M., Sabatini, A., Petrarca, V., and Di Deco, M. A. 1979. Chromosomal differentiation and adaptation to human environments in the Anopheles gambiae complex. Trans. R. Soc. Trop. Med. Hyg., 73(5): 483–497.

Coyne, J. A. and Orr, H. A. 2004. Speciation. Sinauer Assoc., Sunderland, Massachusetts.

Degnan, J. H. 2018. Modeling hybridization under the network multispecies coalescent. Syst. Biol., 67(5): 786–799.

della Torre, A., Merzagora, L., Powell, J., and Coluzzi, M. 1997. Selective introgression of paracentric inversions between two sibling species of the Anopheles gambiae complex. Genetics, 146(1): 239–244.

Dickey, J. M. 1971. The weighted likelihood ratio, linear hypotheses on normal location parameters. Ann. Math. Statist., 42(1): 204–223.

Durand, E. Y., Patterson, N., Reich, D., and Slatkin, M. 2011. Testing for ancient admixture between closely related populations. Mol. Biol. Evol., 28: 2239–2252.

Edelman, N. and Mallet, J. 2021. Prevalence and adaptive impact of introgression. Annu. Rev. Genet., 55(1): 265–283.

Edelman, N. B., Frandsen, P. B., Miyagi, M., Clavijo, B., and Davey, J. e. a. 2019. Genomic architecture and introgression shape a butterfly radiation. Science, 366(6465): 594–599.

Elworth, R. A. L., Ogilvie, H. A., Zhu, J., and Nakhleh, L. 2019. Advances in computational methods for phylogenetic networks in the presence of hybridization. Bioinformatics and Phylogenetics, 29: 317–360.

Feurtey, A. and Stukenbrock, E. H. 2018. Interspecific gene exchange as a driver of adaptive evolution in fungi. Annu. Rev. Microbiol., 72: 377–398.

Flouri, T., Jiao, X., Rannala, B., and Yang, Z. 2018. Species tree inference with bpp using genomic sequences and the multispecies coalescent. Mol. Biol. Evol., 35(10): 2585–2593.

Flouri, T., Jiao, X., Rannala, B., and Yang, Z. 2020. A Bayesian implementation of the multispecies coalescent model with introgression for phylogenomic analysis. Mol. Biol. Evol., 37(4): 1211–1223.

Green, R. E., Krause, J., Briggs, A. W., Maricic, T., and Stenzel, U. e. a. 2010. A draft sequence of the Neandertal genome. Science, 328: 710–722.

Gronau, I., Hubisz, M. J., Gulko, B., Danko, C. G., and Siepel, A. 2011. Bayesian inference of ancient human demography from individual genome sequences. Nature Genet., 43: 1031–1034.

Hey, J. 2010. Isolation with migration models for more than two populations. Mol. Biol. Evol., 27: 905–920.

Hey, J. and Nielsen, R. 2004. Multilocus methods for estimating population sizes, migration rates and divergence time, with applications to the divergence of Drosophila pseudoobscura and D. persimilis. Genetics, 167: 747–760.

Hey, J., Chung, Y., Sethuraman, A., Lachance, J., Tishkoff, S., Sousa, V. C., and Wang, Y. 2018. Phylogeny estimation by integration over isolation with migration models. Mol. Biol. Evol., 35(11): 2805–2818.

Hibbins, M. S. and Hahn, M. W. 2022. Phylogenomic approaches to detecting and characterizing introgression. Genetics, page 10.1093/genetics/iyab173.

Huang, J., Flouri, T., and Yang, Z. 2020. A simulation study to examine the information content in phylogenomic datasets under the multispecies coalescent model. Mol. Biol. Evol., 37(11): 3211–3224.

Huang, J., Thawornwattana, Y., Flour, T., Mallet, J., and Yang, Z. 2022. Inference of gene flow between species under misspecified models. Mol. Biol. Evol., 39(12): msac237.

Ji, J., Jackson, D. J., Leache, A. D., and Yang, Z. 2023. Power of Bayesian and heuristic tests to detect cross-species introgression with reference to gene flow in the Tamias quadrivittatus group of North American chipmunks. Syst. Biol.

Jiao, X., Flouri, T., and Yang, Z. 2021. Multispecies coalescent and its applications to infer species phylogenies and cross-species gene flow. Nat. Sci. Rev., 8: nwab127 (DOI: 10.1093/nsr/nwab127).

Jukes, T. and Cantor, C. 1969. Evolution of protein molecules. In H. Munro, editor, Mammalian Protein Metabolism, pages 21–123. Academic Press, New York.

Kronforst, M. R., Young, L. G., Blume, L. M., and Gilbert, L. E. 2006. Multilocus analyses of admixture and introgression among hybridizing Heliconius butterflies. Evolution, 60(6): 1254–1268.

Kronforst, M. R., Hansen, M. E., Crawford, N. G., Gallant, J. R., Zhang, W., Kulathinal, R. J., Kapan, D. D., and Mullen, S. P. 2013. Hybridization reveals the evolving genomic architecture of speciation. Cell Reports, 5(3): 666–677.

Lartillot, N. and Philippe, H. 2006. Computing Bayes factors using thermodynamic integration. Syst. Biol., 55: 195–207.

Leaché, A. D., Harris, R. B., Rannala, B., and Yang, Z. 2014. The influence of gene flow on bayesian species tree estimation: A simulation study. Syst. Biol., 63(1): 17–30.

Lohse, K. and Frantz, L. A. F. 2014. Neandertal admixture in Eurasia confirmed by maximum likelihood analysis of three genomes. Genetics, 196(4): 1241–1251.

Lohse, K., Chmelik, M., Martin, S. H., and Barton, N. H. 2016. Efficient strategies for calculating blockwise likelihoods under the coalescent. Genetics, 202(2): 775–786.

Mallet, J., Beltrán, M., Neukirchen, W., and Linares, M. 2007. Natural hybridization in heliconiine butterflies: the species boundary as a continuum. BMC Evol. Biol., 7(1): 28.

Marques, D. A., Meier, J. I., and Seehausen, O. 2019. A combinatorial view on speciation and adaptive radiation. Trends Ecol. Evol., 34(6): 531–544.

Martin, S. H. and Jiggins, C. D. 2017. Interpreting the genomic landscape of introgression. Curr. Opin. Genet. Dev., 47: 69–74.

Martin, S. H., Dasmahapatra, K. K., Nadeau, N. J., Salazar, C., Walters, J. R., Simpson, F., Blaxter, M., Manica, A., Mallet, J., and Jiggins, C. D. 2013. Genome-wide evidence for speciation with gene flow in Heliconius butterflies. Genome Res., 23(11): 1817–1828.

Martin, S. H., Eriksson, A., Kozak, K. M., Manica, A., and Jiggins, C. D. 2015. Speciation in Heliconius butterflies: Minimal contact followed by millions of generations of hybridisation. bioRxiv.

Martin, S. H., Davey, J. W., Salazar, C., and Jiggins, C. D. 2019. Recombination rate variation shapes barriers to introgression across butterfly genomes. PLoS Biol., 17(2): e2006288.

Moran, B. M., Payne, C., Langdon, Q., Powell, D. L., Brandvain, Y., and Schumer, M. 2021. The genomic consequences of hybridization. eLife, 10: e69016.

Naisbit, R. E., Jiggins, C. D., and Mallet, J. 2001. Disruptive sexual selection against hybrids contributes to speciation between Heliconius cydno and Heliconius melpomene. Proc. R. Soc. Lond. B, 268(1478): 1849–1854.

Naisbit, R. E., Jiggins, C. D., Linares, M., Salazar, C., and Mallet, J. 2002. Hybrid sterility, Haldane’s rule and speciation in Heliconius cydno and H. melpomene. Genetics, 161(4): 1517–1526.

Nielsen, R. and Wakeley, J. 2001. Distinguishing migration from isolation: a Markov chain Monte Carlo approach. Genetics, 158: 885–896.

Pease, J. B. and Hahn, M. W. 2015. Detection and polarization of introgression in a five-taxon phylogeny. Syst. Biol., 64(4): 651–662.

Peters, K. J., Myers, S. A., Dudaniec, R. Y., O’Connor, J. A., and Kleindorfer, S. 2017. Females drive asymmetrical introgression from rare to common species in Darwin’s tree finches. J. Evol. Biol., 30(11): 1940–1952.

Petry, D. 1983. The effect on neutral gene flow of selection at a linked locus. Theor. Popul. Biol., 23: 300–313.

Rannala, B. and Yang, Z. 2003. Bayes estimation of species divergence times and ancestral population sizes using DNA sequences from multiple loci. Genetics, 164(4): 1645–1656.

Rannala, B. and Yang, Z. 2017. Efficient Bayesian species tree inference under the multispecies coalescent. Syst. Biol., 66: 823–842.

Slotman, M., Torre, A. d., and Powell, J. R. 2004. The genetics of inviability and male sterility in hybrids between Anopheles gambiae and An. arabiensis. Genetics, 167(1): 275–287.

Slotman, M., della Torre, A., and Powell, J. R. 2005a. Female sterility in hybrids between Anopheles gambiae and A. arabiensis, and the causes of Haldane’s rule. Evolution, 59(5): 1016–1026.

Slotman, M. A., della Torre, A., Calzetta, M., and Powell, J. R. 2005b. Differential introgression of chromosomal regions between Anopheles gambiae and An. arabiensis. Am. J. Trop. Med. Hyg., 73(2): 326–335.

Solis-Lemus, C. and Ane, C. 2016. Inferring phylogenetic networks with maximum pseudolikelihood under incomplete lineage sorting. PLoS Genet., 12(3): e1005896.

Solis-Lemus, C., Bastide, P., and Ane, C. 2017. PhyloNetworks: A package for phylogenetic networks. Mol. Biol. Evol., 34(12): 3292–3298.

Tavaré, S. 1984. Lines of descent and genealogical processes, and their applications in population genetics models. Theor. Popul. Biol., 26: 119–164.

Thawornwattana, Y., Dalquen, D., and Yang, Z. 2018. Coalescent analysis of phylogenomic data confidently resolves the species relationships in the Anopheles gambiae species complex. Mol. Biol. Evol., 35(10): 2512–2527.

Thawornwattana, Y., Seixas, F. A., Mallet, J., and Yang, Z. 2022. Full-likelihood genomic analysis clarifies a complex history of species divergence and introgression: the example of the erato-sara group of Heliconius butterflies. Syst. Biol., 71(5): 1159–1177.

Tiley, G. P., Flouri, T., Jiao, X., Poelstra, J. P., Xu, B., Zhu, T., Rannala, B., Yoder, A. D., and Yang, Z. 2023. Estimation of species divergence times in presence of cross-species gene flow. Syst. Biol.

Wakeley, J. 2009. Coalescent Theory: An Introduction. Roberts and Company, Greenwood Village, Colorado.

Wen, D. and Nakhleh, L. 2018. Coestimating reticulate phylogenies and gene trees from multilocus sequence data. Syst. Biol., 67(3): 439–457.

Yang, Z. 2014. Molecular Evolution: A Statistical Approach. Oxford University Press, Oxford, England.

Yang, Z. 2015. The bpp program for species tree estimation and species delimitation. Curr. Zool., 61: 854–865.

Yang, Z. and Flouri, T. 2022. Estimation of cross-species introgression rates using genomic data despite model unidentifiability. Mol. Biol. Evol., 39(5). msac083.

Yang, Z. and Rannala, B. 2014. Unguided species delimitation using DNA sequence data from multiple loci. Mol. Biol. Evol., 31(12): 3125–3135.

Yang, Z. and Zhu, T. 2018. Bayesian selection of misspecified models is overconfident and may cause spurious posterior probabilities for phylogenetic trees. Proc. Natl. Acad. Sci. U.S.A., 115(8): 1854–1859.

Yu, Y., Degnan, J. H., and Nakhleh, L. 2012. The probability of a gene tree topology within a phylogenetic network with applications to hybridization detection. PLoS Genet., 8(4): e1002660.

Yu, Y., Dong, J., Liu, K. J., and Nakhleh, L. 2014. Maximum likelihood inference of reticulate evolutionary histories. Proc. Natl. Acad. Sci. U.S.A., 111(46): 16448–16453.

Zhang, C., Ogilvie, H. A., Drummond, A. J., and Stadler, T. 2018. Bayesian inference of species networks from multilocus sequence data. Mol. Biol. Evol., 35: 504–517.

Zhu, T., Flouri, T., and Yang, Z. 2022. A simulation study to examine the impact of recombination on phylogenomic inferences under the multispecies coalescent model. Mol. Ecol., 31: 2814–2829.

