## supplemental information for "Inferring the direction of introgression using genomic sequence data"

**Supplemental Information for**
**Inferring the direction of introgression using genomic sequence data**

Yuttapong Thawornwattana, Jun Huang, Tomáš Flouri, James Mallet, and Ziheng Yang

### SI text: Simulation results in the case of four species

We simulated datasets under the three MSci models of figure 5 for four species on the species tree $((A, (B, C)), D)$, with introgression between non-sister species $A$ and $B$ in different directions: inflow (I), outflow (O), and bidirectional introgression (B). The data were analyzed under the same three models (I, O, B), resulting in nine combinations.

#### *Four species, equal population sizes*

Figure S4 shows average posterior means and 95% HPD CIs in analysis of data simulated assuming the same population size ($\theta_0 = \theta_1 = 0.01$). The results for the large datasets of $L = 4000$ are summarized in table S2. Parameters shared among all three models (I, O, B) are well estimated, with no discernible impact of model misspecification. These include species divergence and introgression times ($\tau_R, \tau_S, \tau_T$, and $\tau_X = \tau_Y$), population sizes for the extant species ($\theta_A, \theta_B, \theta_C, \theta_D$) and for ancestral species not involved in gene flow ($\theta_R, \theta_S$). As discussed in the case of two species, introgression time is well estimated even if the introgression direction is misspecified (e.g., $\tau_X$ in the I-O and O-I settings), because the estimate is dominated by the minimum sequence divergence between species involved in introgression ($t_{ab}$) (table 1). Ancestral population sizes ($\theta_R, \theta_S$) are slightly less well estimated but appear to converge to the correct values in all settings when the number of loci $L \to \infty$ (fig. S4). Below we focus on introgression probabilities ($\varphi_X, \varphi_Y$) and population sizes $\theta_X$, $\theta_Y$, and $\theta_T$.

In the I-I, O-O, and B-B settings, the true model is assumed in the analysis, and the results provide a reference for comparison. The introgression probability $\varphi_Y$ in the I-I setting is more precisely estimated than $\varphi_X$ in the O-O setting, with narrower CIs. Inflow is easier to infer than outflow, as observed in our simulations for the three-species case (fig. 4**b&c**, same $\theta$). Similarly in the B-B setting, the inflow probability $\varphi_Y$ is better estimated than the outflow probability $\varphi_X$. The B-B setting had slightly wider CIs for $\varphi_X$ and $\varphi_Y$ than in the I-I and O-O settings, due to more parameters in model B (fig. S4, table S2). Overall the introgression probabilities are well estimated under all three settings, although thousands of loci appear necessary to obtain precise estimates. Population size $\theta_X$ is better estimated in the I-I setting than in the O-O and B-B settings, because estimation is affected by uncertainties in $\varphi_X$ in model O and B. Similarly $\theta_Y$ is better estimated in O-O than in I-I and B-B settings.

In the I-B and O-B settings, gene flow is unidirectional but the model assumes bidirectional gene flow. The model B is over-parametrized but not misspecified. As Bayesian estimation under the correct model is consistent, the introgression probability for the nonexistent introgression ($\varphi_X$ in I-B, $\varphi_Y$ in O-B) should converge to 0 when the data size approaches $\infty$. Results in both the I-B and O-B settings are consistent with this expectation (fig. S4). Other parameters are well-estimated, with CI widths indistinguishable from those in the I-I and O-O settings. Over-parametrization in model B incurs little cost to the statistical performance of the method, as in the case of two species (fig. 3).

In the I-O and O-I settings, introgression is assumed to occur in the wrong (opposite) direction. According to our analysis of the two-species case (table 1), this misspecification should only affect estimation of the introgression probability and population sizes $\theta_X, \theta_Y$, and $\theta_T$, while other parameters including introgression time should be correctly estimated. This is indeed the case (fig. S4). In the I-O setting, we expect $\theta_X$ to be underestimated, $\theta_Y$ and $\theta_T$ to be overestimated, and the introgression probability $\varphi_X$ may be larger or smaller than $\varphi_Y$ depending on how the estimates of $\theta_Y$ and $\theta_T$ compare with the true value of $\theta_X$ (table 1). Simulation results confirm these predictions (fig. S4). In the O-I setting, the effects are the opposite: $\theta_X$ was overestimated while $\theta_Y$ and $\theta_T$ were underestimated. As expected, $\varphi_Y$ was estimated to be smaller than $\varphi_X$.

Finally, in the B-O and B-I settings, introgression occurs in both directions but is assumed to occur in only one direction. The estimates of $\theta_X, \theta_Y$, and $\theta_T$ follow the same pattern as in I-O and O-I, respectively. The introgression probability ($\varphi_X$ in B-O, $\varphi_Y$ in B-I) is larger and less well-estimated than when the true model has unidirectional gene flow ($\varphi_X$ in O-I, $\varphi_Y$ in I-O). This positive bias may be explained by the fact that gene flow in the two directions in the true model B have an accumulative effect on the distribution of the coalescent times between species ($t_{ab}$) (see eq. A4). For instance, in the B-I setting, $B \to A$ introgression in the true model B is expected to increase the chance of coalescence during $\tau_X < t_{ab} < \tau_S$, and such introgression events may be recognized and misinterpreted as extra $A \to B$ introgression in the fitting model I, leading to $\hat{\varphi}_Y^{(\mathrm{I})} > \varphi_Y^{(\mathrm{B})}$.

#### *Four species, different population sizes*

Results for cases with different population sizes are summarized in figure S5 and table S3. In our setting, model I assumes inflow from a small population to a large one, model O assumes outflow from a large population to a small one, while model B assumes both inflow from small population to large one as well as outflow from large to small (fig. 5).

As in the case of equal population sizes, species

divergence and introgression times ($\tau_R, \tau_S, \tau_T$, and $\tau_X = \tau_Y$) and population sizes for extant species ($\theta_A, \theta_B, \theta_C, \theta_D$) and for common ancestors $R$ and $S$ ($\theta_R, \theta_S$) are all well estimated, in spite of model misspecification. Thus we focus on introgression probabilities ($\varphi_X, \varphi_Y$) and population sizes $\theta_X$, $\theta_Y$, and $\theta_T$.

In the I-I, O-O, and B-B settings, the correct model is assumed in the analysis. Introgression probability $\varphi_Y$ in model I is far more precisely estimated than $\varphi_X$ in model O. At $L = 4000$ loci, the average 95% CI is 0.19-0.22 for I-I and 0.16-0.24 for O-O (table S3). The difference is far greater than in the case of equal population sizes where the inflow probability $\varphi_Y$ in model I is slightly better estimated than the outflow probability $\varphi_X$ in model O (table S2). It is easier to estimate the introgression probability from a small population to a large one (model I) than in the opposite direction, as discussed before. Similarly in the B-B setting, the inflow probability of small → large introgression ($\varphi_Y$) is much better estimated than the outflow probability of large → small introgression ($\varphi_X$): for $L = 4000$, the average 95% CIs are 0.18-0.23 for $\varphi_Y$ and 0.15-0.24 for $\varphi_X$ (table S3). In the I-B and O-B settings, model B is over-parameterized. Performance is very similar to that in the I-I and O-O settings, respectively, with $\varphi$ for the nonexistent migration approaching 0 with the increase of data size (fig. S5, table S3).

In the I-O and O-I settings, the introgression direction is misspecified. In the I-O setting, $\hat{\varphi}_X$ is much greater than in the case of equal population sizes. The extremely large $\hat{\varphi}_X$ mimics the extreme estimate in the two-species case (fig. 3**c** small→large, model O). In the O-I setting, gene flow is from a large population into a small one, and the donor population size $\theta_Y$ is grossly underestimated while the recipient population size $\theta_X$ is overestimated when introgression direction is misspecified. These patterns are similar to those in the two-species analysis (fig. 3**d** large→small, model O). The estimate $\hat{\varphi}_Y$ is much lower than the true introgression probability $\varphi_X = 0.2$ in the opposite direction.

The B-I and B-O settings show a cumulative effect in the estimates of the migration rate: $\hat{\varphi}_Y$ is greater in the B-I setting than in the I-I setting, and $\hat{\varphi}_X$ is greater in the B-O setting than in the O-O setting. This is the same pattern as found in the case of equal population sizes (fig. S4).

#### *Bayesian test of introgression*

We apply the Bayesian test of introgression (Ji *et al.* (2023)) to the data analyzed in figures S4&S5, with results summarized in figures S6&S7. In the I-I, O-O, and B-O settings, where the correct model is assumed, the power of the test is high, reaching $\sim 100\%$ at $L \geq 1000$ loci (figs. S6&S7). In the I-B and O-B settings, the power of detecting introgression in the direction that exists in the true model (B) is high, while the false positive rate for detecting non-existent introgression in the incorrect direction is low, below the nominal 1%. In the I-O and O-I settings, where introgression direction is misspecified, the false positive rate is very high, comparable to the power in the analysis under the correct model. Overall, the results are similar to those for the two-species simulations (fig. S2).

### References


Ji, J., Jackson, D. J., Leache, A. D., and Yang, Z. 2023. Power of Bayesian and heuristic tests to detect cross-species introgression with reference to gene flow in the *Tamias quadrivittatus* group of North American chipmunks. *Syst. Biol.*

Tavaré, S. 1984. Lines of descent and genealogical processes, and their applications in population genetics models. *Theor. Popul. Biol.*, 26: 119–164.

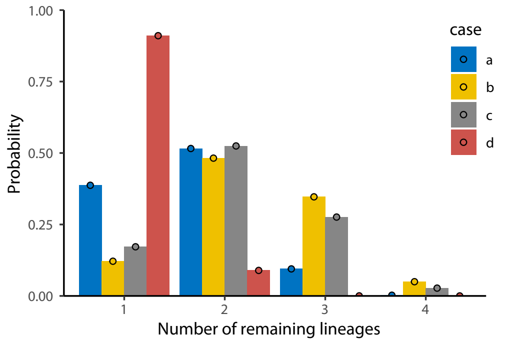


Figure S1: Probability distribution of the number of lineages remaining at time $T$ (in $2N$ generations) when we trace, backwards in time, the genealogy for a sample of $n$ sequences randomly sampled from a diploid population of size $N$. The four cases are for $n = 4$ and $T = 1, 0.5, 0.6$, and 3, for cases **a**–**d** of figure 3. The bars represent estimates from $10^7$ simulations while circles are from eqs. 6.1 & 6.2 in Tavaré (1984).

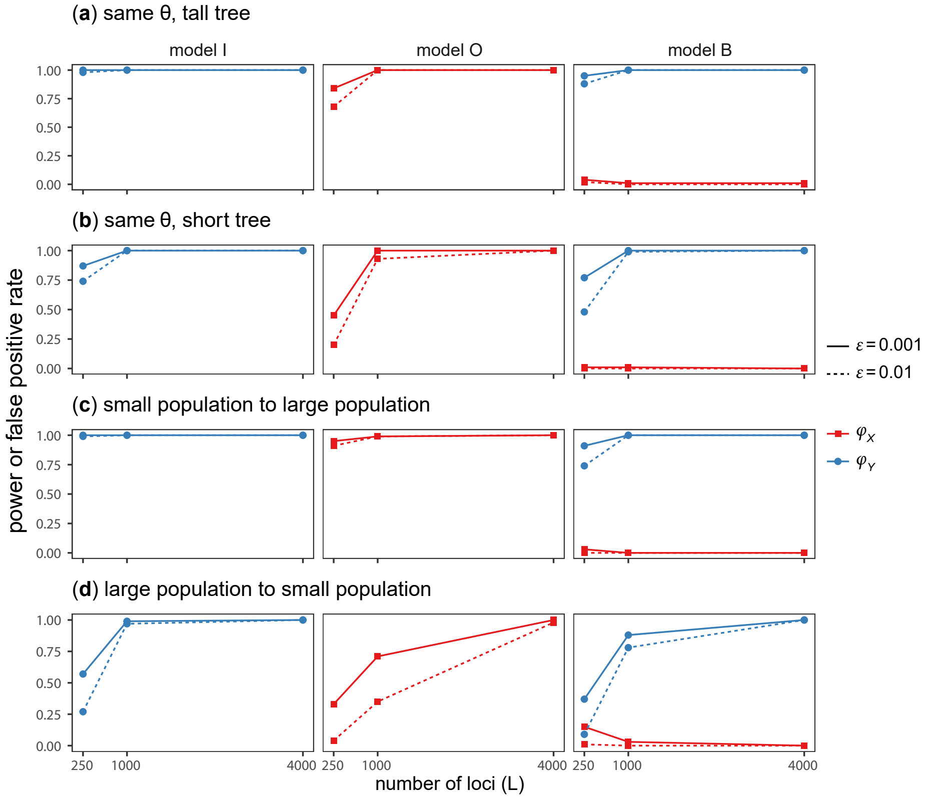


Figure S2: (**2s-power**) Power (blue) and false positive rate (red) of Bayesian test for introgression applied to the simulated data with two species under model I of figure 1**a** using four sets of parameter values (cases **a**–**d**). Bayesian test is conducted using a cut-off of 100 for the Bayes factor, calculated using the Savage-Dickey density ratio with the small value for null effect ($\epsilon = 0.001$ or $0.01$). Note that model I is the true model with $A \rightarrow B$ introgression with probability $\varphi_Y$. A significant result for testing the null $H_0$: $\varphi_Y = 0$ under model I or B is considered a true positive, whereas a significant result for testing the null $H_0$: $\varphi_X = 0$ under model O or B is considered a false positive. Parameter estimates from those data are summarized in figure 3.

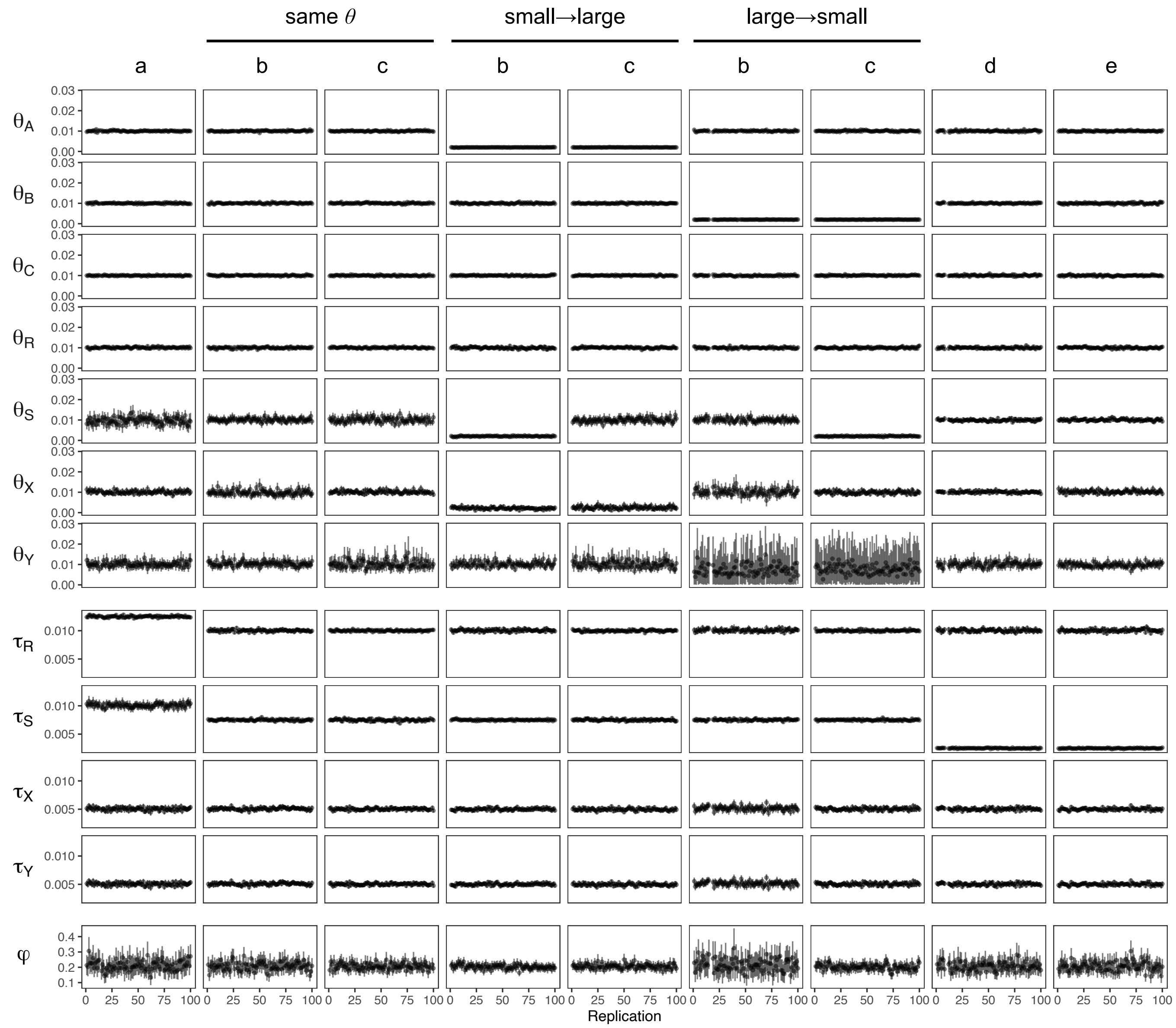


Figure S3: Posterior means and 95% HPD CIs for parameters in 100 replicate datasets simulated and analyzed under the models of figure 4**a**–**e**. Results for $\varphi$ are shown also in figure 4**h**.

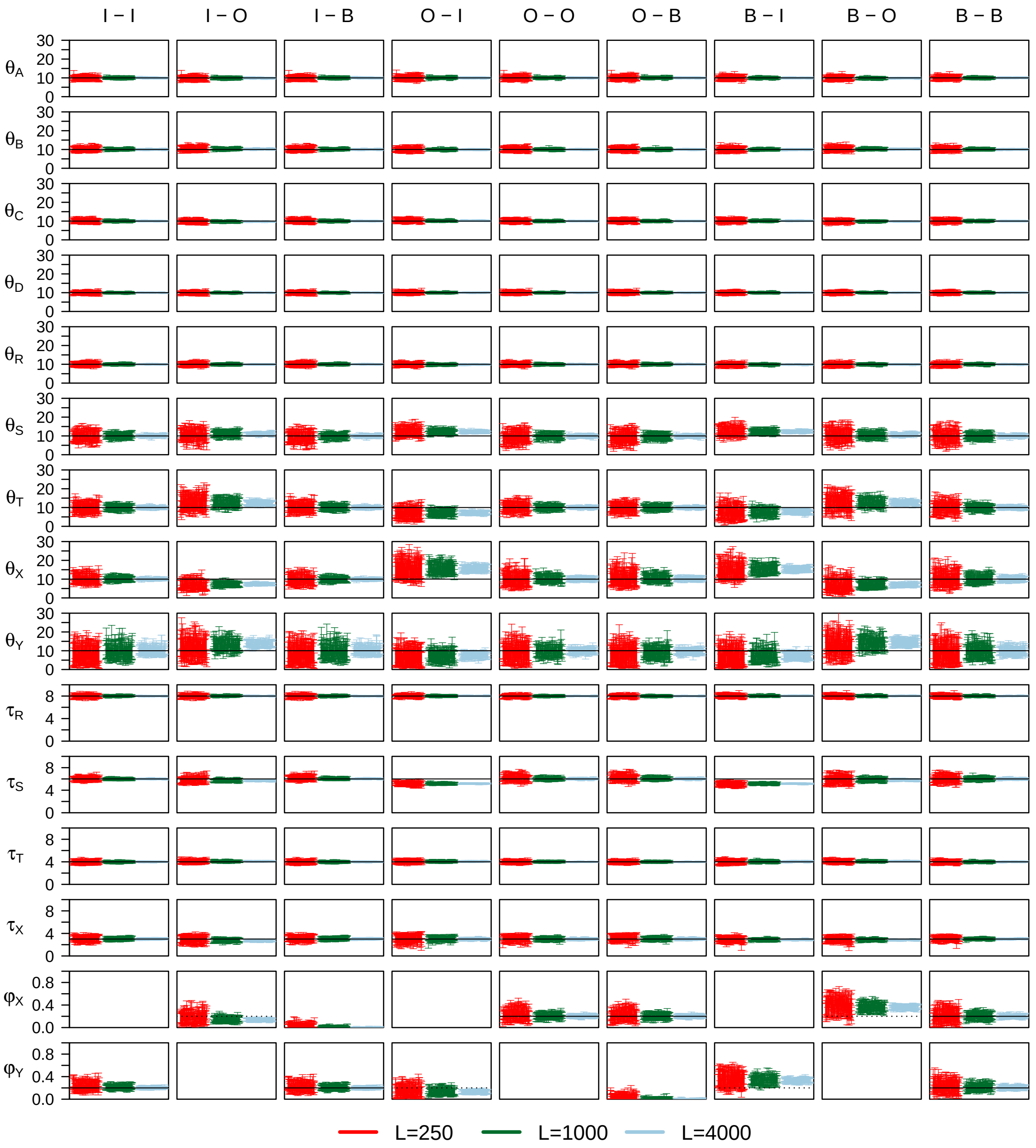


Figure S4: **(4s-same-$\theta$)** The 95% HPD CIs for parameters in 100 replicate datasets simulated and analyzed under models I, O, and B of figure 5, with all species on the species tree having the same population size ($\theta_0 = \theta_1 = 0.01$) when the data are generated. The nine settings are specified in the simulation-analysis format; i.e., 'I-O' means that data were simulated under model I and analyzed under model O. Parameters $\theta$ and $\tau$ are multiplied by $10^3$. Horizontal solid lines indicate the true values. In the I-O setting, the dotted line for $\varphi_X$ in model O indicates the true value of $\varphi_Y$ in model I assumed in the simulation, while in the O-I setting, the dotted line for $\varphi_Y$ indicates the true value of $\varphi_X$ in the assumed O model. In the B-I and B-O settings, two introgression probabilities exist in the simulation model ($\varphi_X, \varphi_Y$) but only one is assumed in the analysis model, and the dotted line indicates its true value.

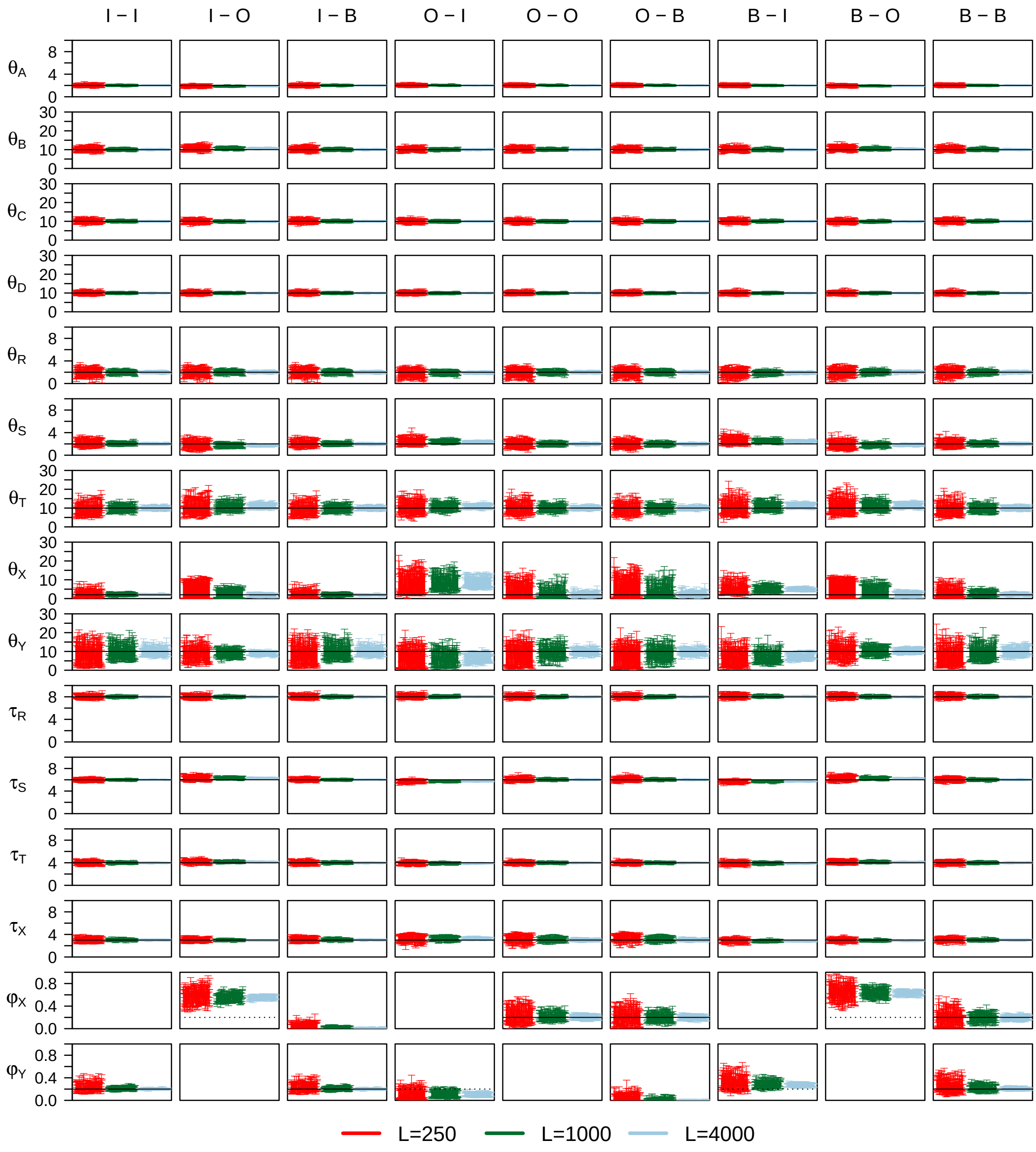


Figure S5: **(4s-diff-$\theta$)** The 95% HPD CIs for parameters in 100 replicate datasets simulated and analyzed under models I, O, and B of figure 5, with $\theta_0$ = 0.002 for the thin branches and $\theta_1$ = 0.01 for the thick branches in the species tree. Other details as in figure S4.

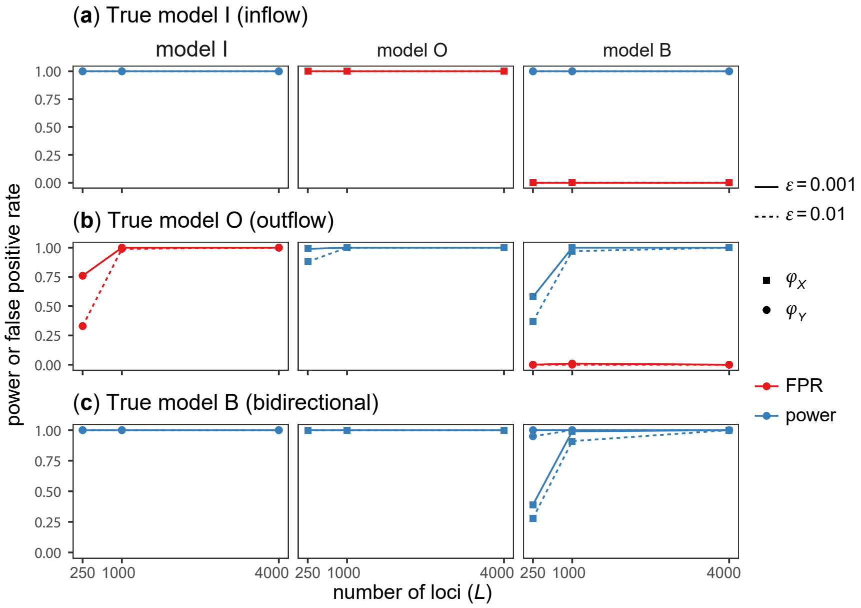


Figure S6: (**4s-same-$\theta$-power**) Power (blue) and false positive rate (red) of Bayesian test for introgression applied to data of four species simulated under the (**a**) inflow (I), (**b**) outflow (O), and (**c**) bidirectional (B) models of figure 5**a**–**c**, assuming the same $\theta$ for all populations. The data were analyzed under the same I, O, and B models, resulting in nine combinations. Parameter estimates from those data are summarized in figure S4.

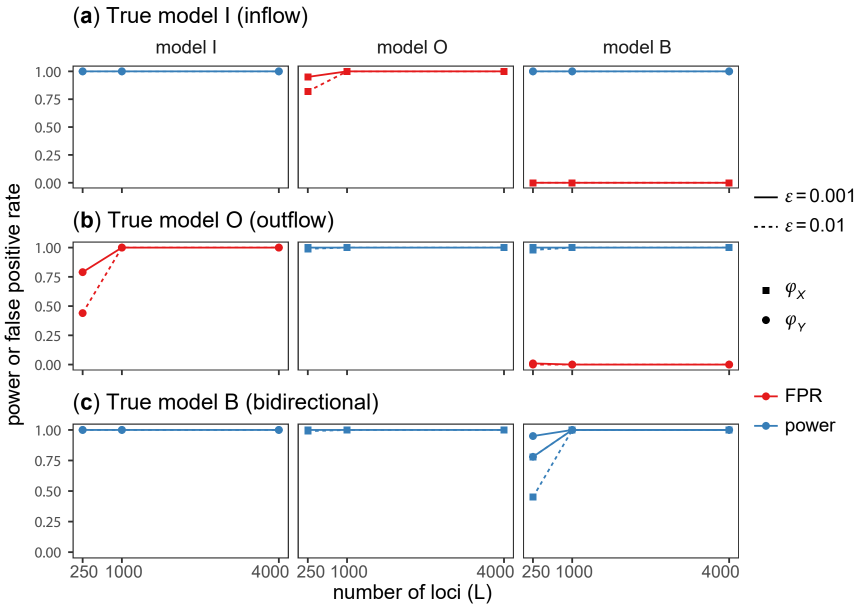


Figure S7: (**4s-diff-$\theta$-power**) Power (blue) and false positive rate (red) of Bayesian test for introgression applied to data of four species simulated under the I, O, and B models of figure 5**a**–**c**, assuming different $\theta$ for populations on the species tree. Parameter estimates from those data are summarized in figure S5. Other details as in figure S6.

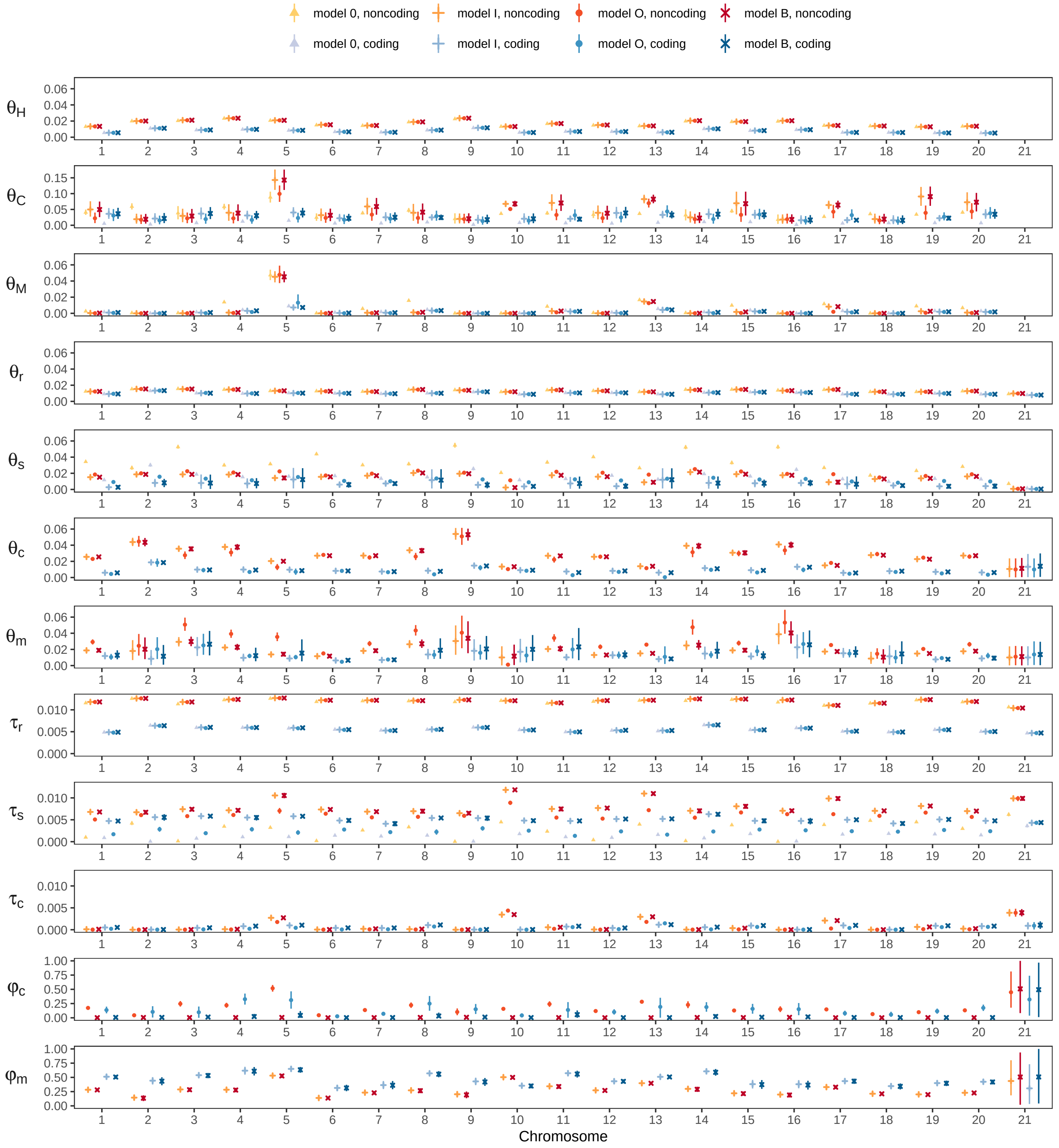


Figure S8: Posterior means and 95% HPD CIs for parameters in BPP analyses of coding and noncoding data from the different chromosomes of *H. cydno* (C), *H. melpomene* (M) and *H. hecale* (*H*) (fig. 5). The four models are (Ø) MSC with no gene flow, (I) MSci with $C \to M$ introgression, (O) MSci with $M \to C$ introgression, and (B) MSci with $C \leftrightarrows M$ bidirectional introgression (see table 2). Results for chromosome 1 are also shown in table 2. In model Ø with no gene flow, branches $C$ and $c$ are assigned the same $\theta_C$, and branches $M$ and $m$ are assigned the same $\theta_M$ (fig. 6). For the sex chromosome (chr 21), there is only one sequence per species per locus, so that the population sizes for extant species ($\theta_C, \theta_M, \theta_H$) are unidentifiable/inestimable in all four models. Furthermore, $\theta_m$ in model I and $\theta_c$ in model O are unidentifiable, and models I and O are unidentifiable (with the parameter mapping $\varphi_m^{(\mathrm{I})} = \varphi_c^{(\mathrm{O})}$ and $\theta_c^{(\mathrm{I})} = \theta_m^{(\mathrm{O})}$). See Discussion for a discussion of unidentifiability issues for data of chromosome 21 (the X chromosome).

**Table S1. (2s-4 cases) Average posterior means and 95% HPD CIs for parameters over 100 replicate datasets of $L$ = 4000 loci simulated under model I (inflow) and analyzed under model I (inflow), model O (outflow) and model B (bidirectional) (fig. 1) using four sets of parameter values (cases a–d)**

| | (**a**) Same $\theta$ tall tree | | | | (**b**) Same $\theta$ short tree | | | | (**c**) Small to large | | | | (**d**) Large to small | | | |
|---|---|---|---|---|---|---|---|---|---|---|---|---|---|---|---|---|
| | $\Theta_{\mathrm{I}}$ | $\hat{\Theta}_{\mathrm{I}}$ | $\hat{\Theta}_{\mathrm{O}}$ | $\hat{\Theta}_{\mathrm{B}}$ | $\Theta_{\mathrm{I}}$ | $\hat{\Theta}_{\mathrm{I}}$ | $\hat{\Theta}_{\mathrm{O}}$ | $\hat{\Theta}_{\mathrm{B}}$ | $\Theta_{\mathrm{I}}$ | $\hat{\Theta}_{\mathrm{I}}$ | $\hat{\Theta}_{\mathrm{O}}$ | $\hat{\Theta}_{\mathrm{B}}$ | $\Theta_{\mathrm{I}}$ | $\hat{\Theta}_{\mathrm{I}}$ | $\hat{\Theta}_{\mathrm{O}}$ | $\hat{\Theta}_{\mathrm{B}}$ |
| $\theta_A$ | 1.0 | 1.00 (0.97, 1.03) | 1.00 (0.97, 1.03) | 1.00 (0.97, 1.03) | 1.00 | 1.01 (0.97, 1.04) | 1.01 (0.97, 1.04) | 1.01 (0.97, 1.04) | 0.2 | 0.20 (0.19, 0.21) | 0.19 (0.19, 0.20) | 0.20 (0.19, 0.21) | 1.0 | 1.00 (0.97, 1.03) | 1.00 (0.97, 1.03) | 1.00 (0.97, 1.03) |
| $\theta_B$ | 1.0 | 1.00 (0.97, 1.03) | 1.00 (0.97, 1.03) | 1.00 (0.97, 1.03) | 1.00 | 1.00 (0.97, 1.04) | 1.00 (0.97, 1.04) | 1.00 (0.97, 1.04) | 1.0 | 1.00 (0.97, 1.03) | 1.02 (0.99, 1.05) | 1.00 (0.97, 1.03) | 0.2 | 0.20 (0.19, 0.21) | 0.20 (0.19, 0.21) | 0.20 (0.19, 0.21) |
| $\theta_R$ | 1.0 | 1.00 (0.96, 1.04) | 1.06 (1.02, 1.10) | 1.00 (0.95, 1.04) | 1.00 | 1.00 (0.97, 1.03) | 1.02 (0.99, 1.05) | 1.00 (0.97, 1.03) | 0.2 | 0.20 (0.16, 0.23) | 0.14 (0.09, 0.19) | 0.19 (0.16, 0.23) | 1.0 | 1.00 (0.96, 1.04) | 1.02 (0.98, 1.05) | 1.00 (0.96, 1.04) |
| $\theta_X$ | 1.0 | 1.01 (0.94, 1.08) | 0.45 (0.33, 0.58) | 0.98 (0.90, 1.06) | 1.00 | 1.00 (0.93, 1.08) | 0.44 (0.24, 0.65) | 0.97 (0.88, 1.05) | 0.2 | 0.20 (0.16, 0.25) | 0.50 (0.01, 1.19) | 0.18 (0.11, 0.24) | 1.0 | 1.00 (0.93, 1.08) | 0.65 (0.40, 0.88) | 0.99 (0.91, 1.07) |
| $\theta_Y$ | 1.0 | 1.00 (0.88, 1.12) | 1.91 (1.69, 2.14) | 1.02 (0.90, 1.15) | 1.00 | 1.00 (0.85, 1.14) | 1.84 (1.56, 2.14) | 1.02 (0.87, 1.17) | 1.0 | 0.98 (0.81, 1.15) | 1.00 (0.88, 1.12) | 1.01 (0.83, 1.19) | 0.2 | 0.21 (0.09, 0.34) | 1.07 (0.56, 1.65) | 0.24 (0.10, 0.39) |
| $\tau_R$ | 1.0 | 1.00 (0.97, 1.03) | 0.93 (0.90, 0.95) | 1.00 (0.98, 1.03) | 0.50 | 0.50 (0.48, 0.52) | 0.47 (0.45, 0.49) | 0.50 (0.48, 0.52) | 0.6 | 0.60 (0.58, 0.63) | 0.69 (0.65, 0.73) | 0.60 (0.58, 0.63) | 0.6 | 0.60 (0.57, 0.63) | 0.57 (0.54, 0.59) | 0.61 (0.57, 0.64) |
| $\tau_X$ | 0.5 | 0.50 (0.47, 0.53) | 0.55 (0.52, 0.57) | 0.50 (0.47, 0.53) | 0.25 | 0.25 (0.23, 0.28) | 0.28 (0.25, 0.31) | 0.25 (0.23, 0.28) | 0.3 | 0.30 (0.28, 0.32) | 0.36 (0.34, 0.37) | 0.31 (0.29, 0.33) | 0.3 | 0.30 (0.27, 0.34) | 0.35 (0.31, 0.38) | 0.31 (0.27, 0.34) |
| $\varphi_X$ | n/a | n/a | 0.27 (0.20, 0.33) | 0.01 (0.00, 0.02) | n/a | n/a | 0.30 (0.18, 0.44) | 0.01 (0.00, 0.04) | n/a | n/a | 0.98 (0.96, 1.00) | 0.02 (0.00, 0.05) | n/a | n/a | 0.17 (0.06, 0.31) | 0.01 (0.00, 0.02) |
| $\varphi_Y$ | 0.2 | 0.20 (0.17, 0.24) | n/a | 0.19 (0.16, 0.23) | 0.20 | 0.21 (0.15, 0.27) | n/a | 0.20 (0.14, 0.26) | 0.2 | 0.21 (0.17, 0.25) | n/a | 0.20 (0.16, 0.25) | 0.2 | 0.21 (0.13, 0.29) | n/a | 0.20 (0.12, 0.29) |

Note.— $\Theta_{\mathrm{I}}$ denotes the true parameter values, while $\hat{\Theta}_{\mathrm{I}}$, $\hat{\Theta}_{\mathrm{O}}$, and $\hat{\Theta}_{\mathrm{B}}$ are estimates under models I, O and B, respectively (fig. 1). There are $n$ = 4 sequences per species per locus and $N$ = 500 sites in the sequence. Values of $\tau$ and $\theta$ are multiplied by 100. Estimates for individual datasets and for all data sizes ($L$ = 250, 1000, 4000) are plotted in figure 3.

**Table S2. (4s-same-$\theta$) Average posterior means and 95% HPD CIs for parameters over 100 replicate datasets of $L$ = 4000 loci simulated and analyzed under the inflow (I), outflow (O) and bidirectional (B) models of figure 5 with the same population size ($\theta$) for all species**

| | True model I | | | | True model O | | | | True model B | | | |
|---|---|---|---|---|---|---|---|---|---|---|---|---|
| | $\Theta_{\mathrm{I}}$ | $\hat{\Theta}_{\mathrm{I}}$ | $\hat{\Theta}_{\mathrm{O}}$ | $\hat{\Theta}_{\mathrm{B}}$ | $\Theta_{\mathrm{O}}$ | $\hat{\Theta}_{\mathrm{I}}$ | $\hat{\Theta}_{\mathrm{O}}$ | $\hat{\Theta}_{\mathrm{B}}$ | $\Theta_{\mathrm{B}}$ | $\hat{\Theta}_{\mathrm{I}}$ | $\hat{\Theta}_{\mathrm{O}}$ | $\hat{\Theta}_{\mathrm{B}}$ |
| $\theta_A$ | 1.0 | 1.00 (0.97, 1.03) | 0.98 (0.95, 1.02) | 1.00 (0.97, 1.03) | 1.0 | 1.00 (0.97, 1.03) | 1.00 (0.97, 1.03) | 1.00 (0.97, 1.03) | 1.0 | 1.00 (0.96, 1.03) | 0.98 (0.95, 1.01) | 1.00 (0.97, 1.03) |
| $\theta_B$ | 1.0 | 1.00 (0.97, 1.03) | 1.02 (0.99, 1.06) | 1.00 (0.97, 1.03) | 1.0 | 0.99 (0.96, 1.03) | 1.00 (0.97, 1.03) | 1.00 (0.97, 1.03) | 1.0 | 0.99 (0.96, 1.03) | 1.02 (0.98, 1.05) | 1.00 (0.97, 1.03) |
| $\theta_C$ | 1.0 | 1.00 (0.97, 1.03) | 0.97 (0.95, 1.00) | 1.00 (0.97, 1.03) | 1.0 | 1.02 (0.99, 1.05) | 1.00 (0.97, 1.03) | 1.00 (0.97, 1.03) | 1.0 | 1.01 (0.98, 1.04) | 0.98 (0.95, 1.00) | 1.00 (0.97, 1.03) |
| $\theta_D$ | 1.0 | 1.00 (0.98, 1.02) | 1.00 (0.98, 1.02) | 1.00 (0.98, 1.02) | 1.0 | 1.00 (0.98, 1.02) | 1.00 (0.98, 1.02) | 1.00 (0.98, 1.02) | 1.0 | 1.00 (0.98, 1.02) | 1.00 (0.98, 1.02) | 1.00 (0.98, 1.02) |
| $\theta_R$ | 1.0 | 1.00 (0.97, 1.03) | 0.99 (0.96, 1.02) | 1.00 (0.97, 1.03) | 1.0 | 0.98 (0.96, 1.01) | 1.00 (0.97, 1.03) | 1.00 (0.97, 1.03) | 1.0 | 0.98 (0.95, 1.01) | 0.99 (0.96, 1.02) | 1.00 (0.97, 1.02) |
| $\theta_S$ | 1.0 | 1.00 (0.93, 1.08) | 1.11 (1.03, 1.19) | 0.99 (0.92, 1.07) | 1.0 | 1.22 (1.16, 1.29) | 0.99 (0.91, 1.07) | 0.99 (0.91, 1.07) | 1.0 | 1.23 (1.17, 1.30) | 1.07 (0.99, 1.16) | 1.00 (0.91, 1.08) |
| $\theta_T$ | 1.0 | 1.00 (0.92, 1.08) | 1.24 (1.12, 1.36) | 1.00 (0.92, 1.08) | 1.0 | 0.71 (0.62, 0.81) | 1.00 (0.93, 1.08) | 1.00 (0.92, 1.07) | 1.0 | 0.77 (0.66, 0.88) | 1.26 (1.15, 1.38) | 1.00 (0.91, 1.08) |
| $\theta_X$ | 1.0 | 1.00 (0.94, 1.07) | 0.74 (0.68, 0.80) | 0.99 (0.92, 1.06) | 1.0 | 1.55 (1.39, 1.72) | 1.00 (0.90, 1.11) | 1.01 (0.91, 1.12) | 1.0 | 1.52 (1.39, 1.66) | 0.69 (0.59, 0.79) | 1.01 (0.88, 1.13) |
| $\theta_Y$ | 1.0 | 1.00 (0.77, 1.26) | 1.36 (1.19, 1.54) | 1.01 (0.77, 1.28) | 1.0 | 0.74 (0.58, 0.91) | 0.98 (0.82, 1.14) | 0.96 (0.80, 1.12) | 1.0 | 0.70 (0.52, 0.90) | 1.44 (1.26, 1.63) | 0.98 (0.76, 1.21) |
| $\tau_R$ | 8.0 | 0.80 (0.79, 0.81) | 0.80 (0.79, 0.81) | 0.80 (0.79, 0.81) | 8.0 | 0.80 (0.79, 0.81) | 0.80 (0.79, 0.81) | 0.80 (0.79, 0.81) | 8.0 | 0.80 (0.80, 0.81) | 0.80 (0.79, 0.81) | 0.80 (0.79, 0.81) |
| $\tau_S$ | 6.0 | 0.60 (0.59, 0.61) | 0.57 (0.56, 0.58) | 0.60 (0.59, 0.61) | 6.0 | 0.52 (0.51, 0.53) | 0.60 (0.59, 0.62) | 0.60 (0.59, 0.62) | 6.0 | 0.52 (0.51, 0.52) | 0.58 (0.56, 0.60) | 0.60 (0.58, 0.62) |
| $\tau_T$ | 4.0 | 0.40 (0.39, 0.41) | 0.41 (0.40, 0.42) | 0.40 (0.39, 0.41) | 4.0 | 0.41 (0.40, 0.42) | 0.40 (0.39, 0.41) | 0.40 (0.39, 0.41) | 4.0 | 0.40 (0.39, 0.41) | 0.41 (0.40, 0.42) | 0.40 (0.39, 0.41) |
| $\tau_X$ | 3.0 | 0.30 (0.29, 0.32) | 0.27 (0.26, 0.29) | 0.30 (0.29, 0.32) | 3.0 | 0.30 (0.28, 0.32) | 0.30 (0.28, 0.32) | 0.30 (0.29, 0.32) | 3.0 | 0.29 (0.28, 0.30) | 0.28 (0.27, 0.30) | 0.30 (0.29, 0.31) |
| $\varphi_X$ | n/a | n/a | 0.13 (0.11, 0.16) | 0.01 (0.00, 0.01) | 0.2 | n/a | 0.20 (0.17, 0.23) | 0.20 (0.17, 0.23) | 0.2 | n/a | 0.36 (0.32, 0.40) | 0.20 (0.17, 0.23) |
| $\varphi_Y$ | 0.2 | 0.20 (0.18, 0.23) | n/a | 0.20 (0.18, 0.22) | n/a | 0.13 (0.10, 0.16) | n/a | 0.01 (0.00, 0.01) | 0.2 | 0.33 (0.29, 0.37) | n/a | 0.21 (0.17, 0.24) |

Note.— $\Theta_{\mathrm{I}}$, $\Theta_{\mathrm{O}}$, and $\Theta_{\mathrm{B}}$ denote the true parameter values in the true model, while $\hat{\Theta}_{\mathrm{I}}$, $\hat{\Theta}_{\mathrm{O}}$ and $\hat{\Theta}_{\mathrm{B}}$ are estimates (fig. 5). Each dataset consists of $L$ = 4000 loci, with $n$ = 4 sequences per species per locus and $N$ = 500 sites in the sequence. Values of $\tau$ and $\theta$ are multiplied by 100. Results for all data sizes with $L$ = 250, 1000 or 4000 loci are shown in figure S4.

**Table S3. (4s-diff-$\theta$) Average posterior means and 95% HPD CIs for parameters over 100 replicate datasets of $L$ = 4000 loci simulated and analyzed under the I, O, and B models of figure 5 with different population sizes**

| | Model I | | | | Model O | | | | Model B | | | |
|---|---|---|---|---|---|---|---|---|---|---|---|---|
| | $\Theta_I$ | $\hat{\Theta}_I$ | $\hat{\Theta}_O$ | $\hat{\Theta}_B$ | $\Theta_O$ | $\hat{\Theta}_I$ | $\hat{\Theta}_O$ | $\hat{\Theta}_B$ | $\Theta_B$ | $\hat{\Theta}_I$ | $\hat{\Theta}_O$ | $\hat{\Theta}_B$ |
| $\theta_A$ | 0.2 | 0.20 (0.19, 0.21) | 0.19 (0.18, 0.19) | 0.20 (0.19, 0.21) | 0.2 | 0.20 (0.19, 0.21) | 0.20 (0.19, 0.21) | 0.20 (0.19, 0.21) | 0.2 | 0.20 (0.19, 0.20) | 0.19 (0.18, 0.20) | 0.20 (0.19, 0.21) |
| $\theta_B$ | 1.0 | 1.00 (0.97, 1.03) | 1.06 (1.03, 1.10) | 1.00 (0.97, 1.03) | 1.0 | 1.00 (0.97, 1.03) | 1.00 (0.97, 1.03) | 1.00 (0.97, 1.03) | 1.0 | 1.00 (0.96, 1.03) | 1.04 (1.01, 1.07) | 1.00 (0.97, 1.03) |
| $\theta_C$ | 1.0 | 1.00 (0.97, 1.03) | 0.99 (0.96, 1.01) | 1.00 (0.97, 1.03) | 1.0 | 1.00 (0.97, 1.03) | 1.00 (0.97, 1.03) | 1.00 (0.97, 1.03) | 1.0 | 1.00 (0.97, 1.03) | 0.99 (0.96, 1.01) | 1.00 (0.97, 1.03) |
| $\theta_D$ | 1.0 | 1.00 (0.97, 1.02) | 1.00 (0.97, 1.02) | 1.00 (0.97, 1.02) | 1.0 | 1.00 (0.97, 1.02) | 1.00 (0.98, 1.02) | 1.00 (0.98, 1.02) | 1.0 | 1.00 (0.97, 1.02) | 1.00 (0.97, 1.02) | 1.00 (0.97, 1.02) |
| $\theta_R$ | 0.2 | 0.20 (0.18, 0.22) | 0.20 (0.19, 0.22) | 0.20 (0.18, 0.22) | 0.2 | 0.19 (0.18, 0.21) | 0.20 (0.18, 0.22) | 0.20 (0.18, 0.22) | 0.2 | 0.19 (0.17, 0.21) | 0.20 (0.18, 0.22) | 0.20 (0.18, 0.22) |
| $\theta_S$ | 0.2 | 0.20 (0.19, 0.21) | 0.17 (0.16, 0.19) | 0.20 (0.18, 0.21) | 0.2 | 0.24 (0.22, 0.25) | 0.20 (0.18, 0.22) | 0.20 (0.18, 0.22) | 0.2 | 0.25 (0.23, 0.26) | 0.18 (0.16, 0.20) | 0.20 (0.18, 0.22) |
| $\theta_T$ | 1.0 | 1.00 (0.91, 1.09) | 1.11 (0.99, 1.23) | 0.99 (0.90, 1.08) | 1.0 | 1.08 (0.97, 1.19) | 1.00 (0.91, 1.10) | 1.00 (0.90, 1.09) | 1.0 | 1.12 (1.00, 1.25) | 1.14 (1.01, 1.27) | 1.00 (0.90, 1.11) |
| $\theta_X$ | 0.2 | 0.20 (0.17, 0.24) | 0.17 (0.07, 0.28) | 0.19 (0.16, 0.23) | 0.2 | 0.85 (0.59, 1.14) | 0.22 (0.11, 0.33) | 0.24 (0.12, 0.37) | 0.2 | 0.48 (0.41, 0.56) | 0.20 (0.05, 0.36) | 0.22 (0.15, 0.29) |
| $\theta_Y$ | 1.0 | 0.99 (0.78, 1.23) | 0.89 (0.78, 1.00) | 1.01 (0.78, 1.25) | 1.0 | 0.64 (0.41, 0.86) | 0.99 (0.82, 1.17) | 0.98 (0.80, 1.16) | 1.0 | 0.71 (0.56, 0.86) | 1.03 (0.91, 1.15) | 0.96 (0.75, 1.19) |
| $\tau_R$ | 8.0 | 0.80 (0.79, 0.81) | 0.80 (0.79, 0.81) | 0.80 (0.79, 0.81) | 8.0 | 0.80 (0.79, 0.81) | 0.80 (0.79, 0.81) | 0.80 (0.79, 0.81) | 8.0 | 0.81 (0.80, 0.82) | 0.80 (0.79, 0.81) | 0.80 (0.79, 0.81) |
| $\tau_S$ | 6.0 | 0.60 (0.59, 0.61) | 0.63 (0.62, 0.64) | 0.60 (0.59, 0.61) | 6.0 | 0.57 (0.56, 0.58) | 0.60 (0.59, 0.61) | 0.60 (0.59, 0.61) | 6.0 | 0.57 (0.56, 0.58) | 0.62 (0.61, 0.63) | 0.60 (0.59, 0.61) |
| $\tau_T$ | 4.0 | 0.40 (0.39, 0.41) | 0.41 (0.40, 0.42) | 0.40 (0.39, 0.41) | 4.0 | 0.39 (0.38, 0.40) | 0.40 (0.39, 0.41) | 0.40 (0.39, 0.41) | 4.0 | 0.39 (0.38, 0.40) | 0.41 (0.40, 0.42) | 0.40 (0.39, 0.41) |
| $\tau_X$ | 3.0 | 0.30 (0.29, 0.31) | 0.30 (0.29, 0.30) | 0.30 (0.29, 0.31) | 3.0 | 0.33 (0.31, 0.35) | 0.30 (0.28, 0.32) | 0.30 (0.28, 0.33) | 3.0 | 0.28 (0.27, 0.29) | 0.29 (0.28, 0.30) | 0.30 (0.29, 0.31) |
| $\varphi_X$ | n/a | n/a | 0.55 (0.51, 0.59) | 0.01 (0.00, 0.02) | 0.2 | n/a | 0.20 (0.16, 0.24) | 0.20 (0.15, 0.24) | 0.2 | n/a | 0.63 (0.59, 0.67) | 0.19 (0.15, 0.24) |
| $\varphi_Y$ | 0.2 | 0.20 (0.19, 0.22) | n/a | 0.20 (0.18, 0.22) | n/a | 0.10 (0.07, 0.14) | n/a | 0.00 (0.00, 0.01) | 0.2 | 0.27 (0.24, 0.30) | n/a | 0.20 (0.18, 0.23) |

Note.— Results for all data sizes with $L$ = 250, 1000 or 4000 loci are in figure S5. See legend to table S2.

**Table S4. Numbers of coding and noncoding loci on each chromosome from the genomic data of *Heliconius* butterflies (fig. 6)**

| | Number of loci | | | Number of loci | |
|---|---|---|---|---|---|
| Chr | Coding | Noncoding | Chr | Coding | Noncoding |
| 1 | 4,942 | 5,341 | 12 | 5,242 | 4,962 |
| 2 | 2,902 | 2,534 | 13 | 4,201 | 5,361 |
| 3 | 2,349 | 3,113 | 14 | 2,480 | 2,629 |
| 4 | 1,998 | 2,839 | 15 | 2,719 | 2,920 |
| 5 | 2,316 | 2,901 | 16 | 2,126 | 3,002 |
| 6 | 3,922 | 4,167 | 17 | 4,380 | 4,390 |
| 7 | 3,717 | 4,301 | 18 | 3,601 | 5,068 |
| 8 | 2,877 | 2,706 | 19 | 4,519 | 4,929 |
| 9 | 1,957 | 2,554 | 20 | 4,659 | 4,505 |
| 10 | 5,172 | 5,540 | 21 | 3,078 | 3,275 |
| 11 | 4,074 | 3,541 | | | |

**Table S5. Posterior means and 95% HPD CIs for parameters in analyses of the coding and noncoding data on each on each chromosome from Heliconius under four models (I, O, B, 0)**

| region | variable | chr | model 0 | model I | model O | model B |
|---|---|---|---|---|---|---|
| noncoding | theta_H | 1 | 0.01313 (0.01268, 0.01357) | 0.01339 (0.01295, 0.01386) | 0.01337 (0.01292, 0.01383) | 0.01339 (0.01294, 0.01385) |
| noncoding | theta_H | 2 | 0.01994 (0.01886, 0.02106) | 0.02003 (0.01892, 0.02112) | 0.02003 (0.01893, 0.02114) | 0.02003 (0.01894, 0.02115) |
| noncoding | theta_H | 3 | 0.02039 (0.01940, 0.02141) | 0.02106 (0.02004, 0.02212) | 0.02102 (0.01998, 0.02205) | 0.02107 (0.02001, 0.02208) |
| noncoding | theta_H | 4 | 0.02330 (0.02206, 0.02452) | 0.02351 (0.02227, 0.02476) | 0.02349 (0.02226, 0.02474) | 0.02351 (0.02229, 0.02476) |
| noncoding | theta_H | 5 | 0.02068 (0.01964, 0.02172) | 0.02091 (0.01985, 0.02195) | 0.02088 (0.01985, 0.02196) | 0.02091 (0.01985, 0.02196) |
| noncoding | theta_H | 6 | 0.01500 (0.01439, 0.01559) | 0.01541 (0.01480, 0.01603) | 0.01541 (0.01480, 0.01603) | 0.01541 (0.01479, 0.01603) |
| noncoding | theta_H | 7 | 0.01432 (0.01377, 0.01488) | 0.01449 (0.01394, 0.01506) | 0.01447 (0.01392, 0.01503) | 0.01449 (0.01392, 0.01504) |
| noncoding | theta_H | 8 | 0.01878 (0.01779, 0.01974) | 0.01893 (0.01794, 0.01992) | 0.01891 (0.01793, 0.01989) | 0.01893 (0.01796, 0.01993) |
| noncoding | theta_H | 9 | 0.02261 (0.02134, 0.02389) | 0.02352 (0.02222, 0.02487) | 0.02351 (0.02220, 0.02485) | 0.02352 (0.02222, 0.02488) |
| noncoding | theta_H | 10 | 0.01319 (0.01275, 0.01363) | 0.01320 (0.01276, 0.01364) | 0.01321 (0.01277, 0.01366) | 0.01320 (0.01275, 0.01364) |
| noncoding | theta_H | 11 | 0.01671 (0.01594, 0.01745) | 0.01691 (0.01614, 0.01767) | 0.01688 (0.01613, 0.01766) | 0.01691 (0.01615, 0.01767) |
| noncoding | theta_H | 12 | 0.01478 (0.01424, 0.01531) | 0.01509 (0.01455, 0.01565) | 0.01507 (0.01453, 0.01563) | 0.01509 (0.01455, 0.01565) |
| noncoding | theta_H | 13 | 0.01390 (0.01341, 0.01436) | 0.01396 (0.01349, 0.01444) | 0.01399 (0.01350, 0.01446) | 0.01396 (0.01348, 0.01444) |
| noncoding | theta_H | 14 | 0.01986 (0.01878, 0.02089) | 0.02047 (0.01939, 0.02157) | 0.02042 (0.01934, 0.02152) | 0.02047 (0.01940, 0.02157) |
| noncoding | theta_H | 15 | 0.01912 (0.01817, 0.02008) | 0.01930 (0.01833, 0.02027) | 0.01928 (0.01831, 0.02025) | 0.01930 (0.01832, 0.02026) |
| noncoding | theta_H | 16 | 0.01965 (0.01869, 0.02064) | 0.02040 (0.01939, 0.02142) | 0.02038 (0.01938, 0.02142) | 0.02040 (0.01936, 0.02140) |
| noncoding | theta_H | 17 | 0.01446 (0.01389, 0.01502) | 0.01452 (0.01395, 0.01508) | 0.01454 (0.01396, 0.01510) | 0.01452 (0.01394, 0.01508) |
| noncoding | theta_H | 18 | 0.01384 (0.01336, 0.01433) | 0.01388 (0.01338, 0.01437) | 0.01387 (0.01339, 0.01436) | 0.01388 (0.01339, 0.01436) |
| noncoding | theta_H | 19 | 0.01283 (0.01237, 0.01327) | 0.01291 (0.01245, 0.01336) | 0.01290 (0.01243, 0.01334) | 0.01291 (0.01245, 0.01337) |
| noncoding | theta_H | 20 | 0.01349 (0.01299, 0.01400) | 0.01363 (0.01311, 0.01414) | 0.01361 (0.01309, 0.01411) | 0.01363 (0.01312, 0.01415) |
| noncoding | theta_H | 21 | n/a | n/a | n/a | n/a |
| noncoding | theta_C | 1 | 0.04072 (0.03288, 0.04964) | 0.04990 (0.02739, 0.07593) | 0.02212 (0.00676, 0.04017) | 0.04970 (0.02577, 0.07497) |
| noncoding | theta_C | 2 | 0.05921 (0.04970, 0.06946) | 0.01919 (0.00517, 0.03656) | 0.01726 (0.00406, 0.03363) | 0.01894 (0.00526, 0.03586) |
| noncoding | theta_C | 3 | 0.03868 (0.01875, 0.06060) | 0.02972 (0.00976, 0.05223) | 0.02195 (0.00648, 0.04031) | 0.02907 (0.00787, 0.05161) |
| noncoding | theta_C | 4 | 0.05852 (0.04968, 0.06785) | 0.03962 (0.01404, 0.06652) | 0.02189 (0.00586, 0.04082) | 0.03841 (0.01352, 0.06606) |
| noncoding | theta_C | 5 | 0.08858 (0.07180, 0.10625) | 0.14309 (0.11183, 0.17617) | 0.09934 (0.07394, 0.12586) | 0.14284 (0.11160, 0.17613) |
| noncoding | theta_C | 6 | 0.02472 (0.01393, 0.03756) | 0.03194 (0.01343, 0.05303) | 0.02333 (0.00774, 0.04092) | 0.03154 (0.01308, 0.05263) |
| noncoding | theta_C | 7 | 0.03862 (0.03394, 0.04330) | 0.05934 (0.03429, 0.08696) | 0.03368 (0.01433, 0.05538) | 0.05917 (0.03332, 0.08634) |
| noncoding | theta_C | 8 | 0.04724 (0.04051, 0.05454) | 0.03956 (0.01398, 0.06763) | 0.02310 (0.00501, 0.04300) | 0.04148 (0.01587, 0.06882) |
| noncoding | theta_C | 9 | 0.02027 (0.00546, 0.03780) | 0.02014 (0.00589, 0.03724) | 0.01961 (0.00578, 0.03648) | 0.02062 (0.00668, 0.03780) |
| noncoding | theta_C | 10 | 0.03674 (0.03386, 0.03966) | 0.06767 (0.05861, 0.07738) | 0.05131 (0.04586, 0.05687) | 0.06767 (0.05832, 0.07728) |
| noncoding | theta_C | 11 | 0.03843 (0.03302, 0.04414) | 0.07060 (0.04612, 0.09807) | 0.03271 (0.01600, 0.05133) | 0.07039 (0.04528, 0.09742) |
| noncoding | theta_C | 12 | 0.03379 (0.02286, 0.04622) | 0.03948 (0.01885, 0.06258) | 0.02267 (0.00737, 0.04078) | 0.03805 (0.01769, 0.06180) |
| noncoding | theta_C | 13 | 0.03666 (0.03364, 0.03974) | 0.08261 (0.06932, 0.09652) | 0.06986 (0.05609, 0.08414) | 0.08250 (0.06965, 0.09671) |
| noncoding | theta_C | 14 | 0.03206 (0.01521, 0.05068) | 0.02477 (0.00849, 0.04387) | 0.01923 (0.00510, 0.03626) | 0.02269 (0.00702, 0.04132) |
| noncoding | theta_C | 15 | 0.04507 (0.03923, 0.05116) | 0.06957 (0.03427, 0.10619) | 0.03299 (0.01071, 0.05760) | 0.06823 (0.03085, 0.10622) |
| noncoding | theta_C | 16 | 0.01781 (0.00436, 0.03419) | 0.01805 (0.00482, 0.03437) | 0.01923 (0.00557, 0.03662) | 0.01830 (0.00511, 0.03454) |
| noncoding | theta_C | 17 | 0.02722 (0.02508, 0.02942) | 0.06394 (0.05126, 0.07772) | 0.04281 (0.02229, 0.06455) | 0.06394 (0.05117, 0.07785) |
| noncoding | theta_C | 18 | 0.03580 (0.03304, 0.03867) | 0.01954 (0.00537, 0.03673) | 0.01528 (0.00336, 0.03018) | 0.01936 (0.00570, 0.03613) |
| noncoding | theta_C | 19 | 0.03424 (0.03154, 0.03699) | 0.09091 (0.06148, 0.12124) | 0.03922 (0.01747, 0.06245) | 0.09059 (0.06150, 0.12248) |
| noncoding | theta_C | 20 | 0.04083 (0.03619, 0.04567) | 0.07281 (0.04470, 0.10424) | 0.04361 (0.01992, 0.06980) | 0.07294 (0.04368, 0.10231) |
| noncoding | theta_C | 21 | n/a | n/a | n/a | n/a |
| noncoding | theta_M | 1 | 0.00257 (0.00208, 0.00307) | 0.00034 (0.00019, 0.00048) | 9e-05 (3e-05, 0.00017) | 0.00033 (0.00017, 0.00049) |
| noncoding | theta_M | 2 | 0.00048 (0.00043, 0.00053) | 1e-05 (0.00000, 1e-05) | 0.00000 (0.00000, 1e-05) | 1e-05 (0.00000, 1e-05) |
| noncoding | theta_M | 3 | 0.00043 (2e-04, 0.00066) | 0.00013 (1e-05, 0.00023) | 8e-05 (2e-05, 0.00016) | 0.00012 (1e-05, 0.00023) |
| noncoding | theta_M | 4 | 0.01388 (0.01275, 0.01505) | 0.00094 (0.00017, 0.00186) | 0.00044 (4e-05, 0.00092) | 0.00088 (0.00012, 0.00180) |
| noncoding | theta_M | 5 | 0.04738 (0.04113, 0.05406) | 0.04521 (0.03834, 0.05215) | 0.04774 (0.03742, 0.05905) | 0.04518 (0.03839, 0.05212) |
| noncoding | theta_M | 6 | 0.00017 (0.00012, 0.00021) | 4e-05 (2e-05, 7e-05) | 3e-05 (1e-05, 5e-05) | 4e-05 (2e-05, 7e-05) |
| noncoding | theta_M | 7 | 0.00573 (0.00529, 0.00617) | 0.00073 (0.00043, 0.00105) | 0.00032 (0.00012, 0.00052) | 0.00072 (0.00042, 0.00102) |
| noncoding | theta_M | 8 | 0.01577 (0.01432, 0.01726) | 0.00116 (0.00014, 0.00240) | 0.00048 (3e-05, 0.00110) | 0.00127 (0.00029, 0.00250) |
| noncoding | theta_M | 9 | 1e-05 (0.00000, 2e-05) | 1e-05 (0.00000, 1e-05) | 1e-05 (0.00000, 1e-05) | 1e-05 (0.00000, 1e-05) |
| noncoding | theta_M | 10 | 4e-05 (3e-05, 5e-05) | 4e-05 (4e-05, 5e-05) | 4e-05 (4e-05, 5e-05) | 4e-05 (4e-05, 5e-05) |
| noncoding | theta_M | 11 | 0.00855 (0.00774, 0.00935) | 0.00280 (0.00197, 0.00363) | 0.00123 (6e-04, 0.00186) | 0.00279 (0.00196, 0.00363) |
| noncoding | theta_M | 12 | 0.00064 (0.00049, 0.00078) | 0.00012 (6e-05, 0.00018) | 5e-05 (2e-05, 9e-05) | 0.00011 (4e-05, 0.00017) |
| noncoding | theta_M | 13 | 0.01665 (0.01574, 0.01759) | 0.01462 (0.01366, 0.01556) | 0.01261 (0.01127, 0.01395) | 0.01462 (0.01367, 0.01557) |
| noncoding | theta_M | 14 | 5e-04 (0.00024, 0.00076) | 0.00011 (4e-05, 0.00021) | 8e-05 (2e-05, 0.00017) | 9e-05 (2e-05, 0.00016) |
| noncoding | theta_M | 15 | 0.00983 (0.00910, 0.01059) | 0.00177 (5e-04, 0.00298) | 0.00044 (4e-05, 0.00089) | 0.00172 (0.00042, 0.00309) |
| noncoding | theta_M | 16 | 1e-05 (0.00000, 3e-05) | 1e-05 (0.00000, 1e-05) | 1e-05 (0.00000, 2e-05) | 1e-05 (0.00000, 1e-05) |
| noncoding | theta_M | 17 | 0.01166 (0.01098, 0.01233) | 0.00825 (0.00733, 0.00918) | 0.00192 (0.00071, 0.00305) | 0.00826 (0.00732, 0.00918) |
| noncoding | theta_M | 18 | 8e-05 (6e-05, 9e-05) | 0.00000 (0.00000, 0.00000) | 0.00000 (0.00000, 0.00000) | 0.00000 (0.00000, 1e-05) |
| noncoding | theta_M | 19 | 0.00894 (0.00852, 0.00937) | 0.00256 (0.00168, 0.00344) | 0.00065 (0.00021, 0.00108) | 0.00254 (0.00167, 0.00348) |
| noncoding | theta_M | 20 | 0.00672 (0.00628, 0.00717) | 0.00093 (0.00056, 0.00130) | 0.00042 (9e-05, 0.00066) | 0.00093 (0.00058, 0.00129) |
| noncoding | theta_M | 21 | n/a | n/a | n/a | n/a |
| noncoding | theta_r | 1 | 0.01237 (0.01192, 0.01282) | 0.01228 (0.01181, 0.01274) | 0.01222 (0.01176, 0.01268) | 0.01228 (0.01181, 0.01274) |
| noncoding | theta_r | 2 | 0.01530 (0.01449, 0.01614) | 0.01539 (0.01455, 0.01623) | 0.01535 (0.01450, 0.01618) | 0.01538 (0.01456, 0.01623) |
| noncoding | theta_r | 3 | 0.01562 (0.01498, 0.01624) | 0.01526 (0.01461, 0.01590) | 0.01522 (0.01456, 0.01585) | 0.01526 (0.01462, 0.01591) |
| noncoding | theta_r | 4 | 0.01464 (0.01397, 0.01530) | 0.01470 (0.01401, 0.01536) | 0.01465 (0.01398, 0.01532) | 0.01469 (0.01402, 0.01537) |
| noncoding | theta_r | 5 | 0.01304 (0.01242, 0.01366) | 0.01309 (0.01245, 0.01372) | 0.01302 (0.01240, 0.01365) | 0.01309 (0.01246, 0.01374) |
| noncoding | theta_r | 6 | 0.01276 (0.01223, 0.01329) | 0.01256 (0.01201, 0.01310) | 0.01253 (0.01199, 0.01308) | 0.01256 (0.01200, 0.01310) |
| noncoding | theta_r | 7 | 0.01215 (0.01164, 0.01265) | 0.01209 (0.01158, 0.01262) | 0.01206 (0.01154, 0.01257) | 0.01208 (0.01156, 0.01260) |
| noncoding | theta_r | 8 | 0.01458 (0.01389, 0.01528) | 0.01466 (0.01394, 0.01536) | 0.01462 (0.01391, 0.01533) | 0.01466 (0.01395, 0.01538) |
| noncoding | theta_r | 9 | 0.01416 (0.01344, 0.01486) | 0.01375 (0.01304, 0.01448) | 0.01373 (0.01300, 0.01444) | 0.01375 (0.01302, 0.01447) |
| noncoding | theta_r | 10 | 0.01172 (0.01125, 0.01216) | 0.01189 (0.01143, 0.01235) | 0.01185 (0.01139, 0.01232) | 0.01189 (0.01144, 0.01236) |
| noncoding | theta_r | 11 | 0.01403 (0.01342, 0.01463) | 0.01411 (0.01347, 0.01472) | 0.01402 (0.01340, 0.01463) | 0.01410 (0.01347, 0.01472) |
| noncoding | theta_r | 12 | 0.01310 (0.01260, 0.01360) | 0.01306 (0.01253, 0.01357) | 0.01297 (0.01246, 0.01348) | 0.01305 (0.01252, 0.01356) |
| noncoding | theta_r | 13 | 0.01173 (0.01130, 0.01215) | 0.01183 (0.01139, 0.01227) | 0.01173 (0.01131, 0.01218) | 0.01183 (0.01138, 0.01227) |
| noncoding | theta_r | 14 | 0.01463 (0.01391, 0.01535) | 0.01425 (0.01350, 0.01498) | 0.01423 (0.01350, 0.01495) | 0.01425 (0.01352, 0.01499) |
| noncoding | theta_r | 15 | 0.01473 (0.01406, 0.01540) | 0.01476 (0.01406, 0.01544) | 0.01473 (0.01407, 0.01544) | 0.01476 (0.01407, 0.01544) |
| noncoding | theta_r | 16 | 0.01370 (0.01309, 0.01430) | 0.01327 (0.01266, 0.01390) | 0.01324 (0.01263, 0.01386) | 0.01327 (0.01263, 0.01387) |
| noncoding | theta_r | 17 | 0.01449 (0.01396, 0.01503) | 0.01468 (0.01412, 0.01521) | 0.01459 (0.01404, 0.01512) | 0.01468 (0.01413, 0.01523) |
| noncoding | theta_r | 18 | 0.01185 (0.01138, 0.01231) | 0.01192 (0.01145, 0.01239) | 0.01190 (0.01143, 0.01236) | 0.01192 (0.01145, 0.01239) |
| noncoding | theta_r | 19 | 0.01184 (0.01136, 0.01231) | 0.01192 (0.01142, 0.01240) | 0.01187 (0.01140, 0.01236) | 0.01192 (0.01143, 0.01241) |
| noncoding | theta_r | 20 | 0.01271 (0.01221, 0.01321) | 0.01278 (0.01225, 0.01329) | 0.01274 (0.01223, 0.01326) | 0.01278 (0.01226, 0.01330) |
| noncoding | theta_r | 21 | 0.00924 (0.00856, 0.00989) | 0.00981 (0.00913, 0.01048) | 0.00981 (0.00914, 0.01048) | 0.00979 (0.00912, 0.01046) |
| noncoding | theta_s | 1 | 0.03431 (0.03283, 0.03578) | 0.01517 (0.01412, 0.01624) | 0.01847 (0.01751, 0.01943) | 0.01518 (0.01413, 0.01627) |
| noncoding | theta_s | 2 | 0.02672 (0.02433, 0.02913) | 0.01865 (0.01680, 0.02046) | 0.01996 (0.01817, 0.02177) | 0.01866 (0.01683, 0.02051) |
| noncoding | theta_s | 3 | 0.05265 (0.05046, 0.05499) | 0.01863 (0.01678, 0.02046) | 0.02258 (0.02075, 0.02429) | 0.01867 (0.01683, 0.02051) |
| noncoding | theta_s | 4 | 0.03005 (0.02822, 0.03188) | 0.01844 (0.01678, 0.02014) | 0.02098 (0.01938, 0.02266) | 0.01847 (0.01677, 0.02014) |
| noncoding | theta_s | 5 | 0.03150 (0.02981, 0.03315) | 0.01420 (0.01131, 0.01708) | 0.02236 (0.02027, 0.02445) | 0.01423 (0.01136, 0.01713) |
| noncoding | theta_s | 6 | 0.04387 (0.04213, 0.04559) | 0.01560 (0.01436, 0.01686) | 0.01727 (0.01606, 0.01848) | 0.01560 (0.01437, 0.01688) |
| noncoding | theta_s | 7 | 0.03024 (0.02876, 0.03175) | 0.01702 (0.01580, 0.01825) | 0.01958 (0.01844, 0.02076) | 0.01703 (0.01579, 0.01824) |
| noncoding | theta_s | 8 | 0.03154 (0.02957, 0.03362) | 0.02040 (0.01839, 0.02238) | 0.02325 (0.02143, 0.02506) | 0.02040 (0.01837, 0.02241) |
| noncoding | theta_s | 9 | 0.05469 (0.05175, 0.05773) | 0.01949 (0.01760, 0.02139) | 0.02074 (0.01888, 0.02262) | 0.01948 (0.01758, 0.02133) |
| noncoding | theta_s | 10 | 0.02071 (0.01968, 0.02173) | 0.00239 (0.00036, 0.00455) | 0.01126 (0.00961, 0.01286) | 0.00231 (0.00033, 0.00446) |
| noncoding | theta_s | 11 | 0.03362 (0.03177, 0.03547) | 0.01754 (0.01563, 0.01937) | 0.02189 (0.02023, 0.02358) | 0.01757 (0.01570, 0.01944) |
| noncoding | theta_s | 12 | 0.04022 (0.03886, 0.04162) | 0.01579 (0.01443, 0.01715) | 0.02070 (0.01946, 0.02192) | 0.01582 (0.01445, 0.01718) |
| noncoding | theta_s | 13 | 0.02653 (0.02546, 0.02764) | 0.00887 (0.00641, 0.01127) | 0.01836 (0.01720, 0.01957) | 0.00890 (0.00643, 0.01125) |
| noncoding | theta_s | 14 | 0.05227 (0.04978, 0.05477) | 0.02144 (0.01926, 0.02356) | 0.02514 (0.02303, 0.02720) | 0.02150 (0.01936, 0.02368) |
| noncoding | theta_s | 15 | 0.03304 (0.03093, 0.03522) | 0.01889 (0.01683, 0.02103) | 0.02222 (0.02043, 0.02407) | 0.01892 (0.01684, 0.02100) |
| noncoding | theta_s | 16 | 0.05265 (0.05016, 0.05515) | 0.01758 (0.01593, 0.01920) | 0.01924 (0.01763, 0.02090) | 0.01759 (0.01592, 0.01921) |
| noncoding | theta_s | 17 | 0.02682 (0.02550, 0.02821) | 0.00901 (0.00595, 0.01200) | 0.01890 (0.01755, 0.02027) | 0.00903 (0.00594, 0.01200) |
| noncoding | theta_s | 18 | 0.01740 (0.01648, 0.01833) | 0.01294 (0.01195, 0.01391) | 0.01488 (0.01390, 0.01585) | 0.01294 (0.01196, 0.01391) |
| noncoding | theta_s | 19 | 0.02314 (0.02206, 0.02424) | 0.01375 (0.01263, 0.01491) | 0.01667 (0.01570, 0.01768) | 0.01372 (0.01259, 0.01487) |
| noncoding | theta_s | 20 | 0.02828 (0.02687, 0.02966) | 0.01610 (0.01488, 0.01730) | 0.01879 (0.01765, 0.01988) | 0.01609 (0.01489, 0.01731) |
| noncoding | theta_s | 21 | 0.00710 (0.00582, 0.00840) | 0.00079 (0.00016, 0.00172) | 0.00079 (8e-05, 0.00167) | 8e-04 (0.00011, 0.00161) |
| noncoding | theta_c | 1 | n/a | 0.00592 (0.00553, 0.00632) | 0.00438 (0.00324, 0.00548) | 0.00577 (0.00528, 0.00624) |
| noncoding | theta_c | 2 | n/a | 0.01875 (0.01714, 0.02040) | 0.01842 (0.01329, 0.02355) | 0.01848 (0.01675, 0.02026) |
| noncoding | theta_c | 3 | n/a | 0.00980 (0.00892, 0.01069) | 0.00918 (0.00644, 0.01181) | 0.00938 (0.00817, 0.01052) |
| noncoding | theta_c | 4 | n/a | 0.00984 (0.00871, 0.01100) | 0.00687 (0.00452, 0.00919) | 0.00925 (0.00750, 0.01087) |
| noncoding | theta_c | 5 | n/a | 0.00966 (0.00875, 0.01057) | 0.00702 (0.00331, 0.01108) | 0.00850 (0.00620, 0.01025) |
| noncoding | theta_c | 6 | n/a | 0.00822 (0.00758, 0.00886) | 0.00821 (0.00748, 0.00892) | 0.00814 (0.00749, 0.00881) |
| noncoding | theta_c | 7 | n/a | 0.00739 (0.00673, 0.00805) | 0.00660 (0.00569, 0.00752) | 0.00728 (0.00658, 0.00794) |
| noncoding | theta_c | 8 | n/a | 0.00843 (0.00765, 0.00925) | 0.00393 (0.00156, 0.00624) | 0.00761 (0.00584, 0.00901) |
| noncoding | theta_c | 9 | n/a | 0.01472 (0.01334, 0.01611) | 0.01211 (0.00888, 0.01527) | 0.01428 (0.01272, 0.01594) |
| noncoding | theta_c | 10 | n/a | 0.00908 (0.00859, 0.00956) | 0.00837 (0.00739, 0.00929) | 0.00899 (0.00847, 0.00952) |
| noncoding | theta_c | 11 | n/a | 0.00741 (0.00680, 0.00803) | 0.00289 (0.00118, 0.00462) | 0.00614 (0.00433, 0.00777) |
| noncoding | theta_c | 12 | n/a | 0.00822 (0.00769, 0.00875) | 0.00696 (0.00577, 0.00807) | 0.00814 (0.00760, 0.00870) |
| noncoding | theta_c | 13 | n/a | 0.00604 (0.00559, 0.00649) | 0.00034 (4e-05, 0.00081) | 0.00596 (0.00549, 0.00644) |
| noncoding | theta_c | 14 | n/a | 0.01163 (0.01063, 0.01269) | 0.00956 (0.00705, 0.01195) | 0.01086 (0.00917, 0.01239) |
| noncoding | theta_c | 15 | n/a | 0.00904 (0.00799, 0.01013) | 0.00625 (0.00403, 0.00845) | 0.00873 (0.00748, 0.00995) |
| noncoding | theta_c | 16 | n/a | 0.01325 (0.01201, 0.01442) | 0.00953 (0.00656, 0.01240) | 0.01271 (0.01113, 0.01423) |
| noncoding | theta_c | 17 | n/a | 0.00577 (0.00531, 0.00624) | 0.00464 (0.00360, 0.00564) | 0.00564 (0.00512, 0.00617) |
| noncoding | theta_c | 18 | n/a | 0.00793 (0.00740, 0.00845) | 0.00688 (0.00586, 0.00785) | 0.00783 (0.00730, 0.00843) |
| noncoding | theta_c | 19 | n/a | 0.00701 (0.00646, 0.00755) | 0.00505 (0.00390, 0.00622) | 0.00692 (0.00637, 0.00751) |
| noncoding | theta_c | 20 | n/a | 0.00622 (0.00575, 0.00671) | 0.00346 (0.00244, 0.00448) | 0.00613 (0.00559, 0.00664) |
| noncoding | theta_c | 21 | n/a | 0.01326 (0.00049, 0.02916) | 0.00994 (2e-04, 0.02363) | 0.01392 (0.00074, 0.02982) |
| noncoding | theta_m | 1 | n/a | 0.01186 (0.00764, 0.01684) | 0.01052 (0.00716, 0.01443) | 0.01285 (0.00774, 0.01890) |
| noncoding | theta_m | 2 | n/a | 0.00837 (0.00067, 0.01928) | 0.02025 (0.00812, 0.03512) | 0.01150 (0.00109, 0.02548) |
| noncoding | theta_m | 3 | n/a | 0.02248 (0.01182, 0.03529) | 0.02500 (0.01333, 0.03935) | 0.02642 (0.01258, 0.04288) |
| noncoding | theta_m | 4 | n/a | 0.00955 (0.00512, 0.01472) | 0.01189 (0.00931, 0.01474) | 0.01224 (0.00532, 0.02211) |
| noncoding | theta_m | 5 | n/a | 0.00878 (0.00483, 0.01332) | 0.01040 (0.00769, 0.01322) | 0.01539 (0.00522, 0.03242) |
| noncoding | theta_m | 6 | n/a | 0.00633 (0.00424, 0.00863) | 0.00466 (0.00378, 0.00561) | 0.00652 (0.00433, 0.00897) |
| noncoding | theta_m | 7 | n/a | 0.00688 (0.00477, 0.00916) | 0.00748 (0.00573, 0.00938) | 0.00711 (0.00486, 0.00957) |
| noncoding | theta_m | 8 | n/a | 0.01368 (0.00779, 0.02062) | 0.01323 (0.00823, 0.01931) | 0.01897 (0.00836, 0.03370) |
| noncoding | theta_m | 9 | n/a | 0.01822 (0.00617, 0.03263) | 0.01581 (0.00742, 0.02590) | 0.02056 (0.00726, 0.03662) |
| noncoding | theta_m | 10 | n/a | 0.01701 (0.00390, 0.03312) | 0.01254 (0.00381, 0.02342) | 0.02008 (0.00570, 0.03780) |
| noncoding | theta_m | 11 | n/a | 0.01017 (0.00639, 0.01451) | 0.01999 (0.00881, 0.03413) | 0.02309 (0.00670, 0.04651) |
| noncoding | theta_m | 12 | n/a | 0.01271 (0.00826, 0.01784) | 0.01266 (0.00861, 0.01736) | 0.01329 (0.00845, 0.01879) |
| noncoding | theta_m | 13 | n/a | 0.00798 (0.00574, 0.01038) | 0.01082 (0.00196, 0.02381) | 0.00826 (0.00590, 0.01084) |
| noncoding | theta_m | 14 | n/a | 0.01480 (0.00756, 0.02350) | 0.01327 (0.00924, 0.01775) | 0.01777 (0.00836, 0.02951) |
| noncoding | theta_m | 15 | n/a | 0.01125 (0.00709, 0.01621) | 0.01775 (0.01172, 0.02492) | 0.01212 (0.00728, 0.01782) |
| noncoding | theta_m | 16 | n/a | 0.02264 (0.00868, 0.03850) | 0.02669 (0.01434, 0.04141) | 0.02575 (0.01028, 0.04349) |

| | | | | | | |
|---|---|---|---|---|---|---|
| noncoding | theta_m | 17 | n/a | 0.01523 (0.00964, 0.02182) | 0.01474 (0.01010, 0.02028) | 0.01623 (0.01000, 0.02383) |
| noncoding | theta_m | 18 | n/a | 0.01152 (0.00097, 0.02512) | 0.00981 (0.00285, 0.01921) | 0.01441 (0.00185, 0.03000) |
| noncoding | theta_m | 19 | n/a | 0.00760 (0.00563, 0.00969) | 0.00927 (0.00716, 0.01153) | 0.00776 (0.00577, 0.00998) |
| noncoding | theta_m | 20 | n/a | 0.00881 (0.00620, 0.01186) | 0.01208 (0.00888, 0.01576) | 0.00914 (0.00624, 0.01230) |
| noncoding | theta_m | 21 | n/a | 0.01000 (0.00026, 0.02389) | 0.01372 (0.00069, 0.02998) | 0.01367 (0.00049, 0.02945) |
| noncoding | tau_r | 1 | 0.00487 (0.00472, 0.00501) | 0.00489 (0.00474, 0.00504) | 0.00480 (0.00465, 0.00495) | 0.00489 (0.00474, 0.00503) |
| noncoding | tau_r | 2 | 0.00644 (0.00620, 0.00667) | 0.00638 (0.00614, 0.00661) | 0.00636 (0.00613, 0.00660) | 0.00638 (0.00613, 0.00660) |
| noncoding | tau_r | 3 | 0.00593 (0.00570, 0.00616) | 0.00600 (0.00576, 0.00623) | 0.00584 (0.00561, 0.00607) | 0.00600 (0.00576, 0.00623) |
| noncoding | tau_r | 4 | 0.00595 (0.00569, 0.00619) | 0.00596 (0.00571, 0.00620) | 0.00590 (0.00564, 0.00614) | 0.00596 (0.00570, 0.00620) |
| noncoding | tau_r | 5 | 0.00579 (0.00557, 0.00602) | 0.00589 (0.00567, 0.00611) | 0.00579 (0.00556, 0.00601) | 0.00589 (0.00567, 0.00611) |
| noncoding | tau_r | 6 | 0.00551 (0.00533, 0.00569) | 0.00548 (0.00529, 0.00565) | 0.00543 (0.00525, 0.00561) | 0.00548 (0.00530, 0.00565) |
| noncoding | tau_r | 7 | 0.00532 (0.00514, 0.00549) | 0.00528 (0.00510, 0.00545) | 0.00526 (0.00508, 0.00543) | 0.00528 (0.00510, 0.00545) |
| noncoding | tau_r | 8 | 0.00549 (0.00528, 0.00569) | 0.00554 (0.00534, 0.00574) | 0.00546 (0.00525, 0.00566) | 0.00554 (0.00533, 0.00573) |
| noncoding | tau_r | 9 | 0.00601 (0.00575, 0.00627) | 0.00597 (0.00572, 0.00623) | 0.00595 (0.00568, 0.00621) | 0.00597 (0.00570, 0.00622) |
| noncoding | tau_r | 10 | 0.00542 (0.00525, 0.00557) | 0.00540 (0.00524, 0.00555) | 0.00535 (0.00519, 0.00551) | 0.00540 (0.00524, 0.00556) |
| noncoding | tau_r | 11 | 0.00494 (0.00478, 0.00511) | 0.00496 (0.00479, 0.00512) | 0.00492 (0.00475, 0.00509) | 0.00496 (0.00478, 0.00512) |
| noncoding | tau_r | 12 | 0.00529 (0.00513, 0.00545) | 0.00533 (0.00517, 0.00549) | 0.00518 (0.00501, 0.00533) | 0.00533 (0.00517, 0.00549) |
| noncoding | tau_r | 13 | 0.00516 (0.00499, 0.00532) | 0.00528 (0.00511, 0.00543) | 0.00516 (0.00499, 0.00532) | 0.00528 (0.00510, 0.00544) |
| noncoding | tau_r | 14 | 0.00654 (0.00630, 0.00677) | 0.00656 (0.00632, 0.00679) | 0.00647 (0.00624, 0.00671) | 0.00655 (0.00631, 0.00678) |
| noncoding | tau_r | 15 | 0.00542 (0.00522, 0.00563) | 0.00542 (0.00521, 0.00562) | 0.00538 (0.00517, 0.00558) | 0.00542 (0.00521, 0.00562) |
| noncoding | tau_r | 16 | 0.00586 (0.00562, 0.00610) | 0.00582 (0.00557, 0.00605) | 0.00581 (0.00557, 0.00605) | 0.00582 (0.00558, 0.00605) |
| noncoding | tau_r | 17 | 0.00507 (0.00492, 0.00523) | 0.00514 (0.00497, 0.00529) | 0.00502 (0.00486, 0.00518) | 0.00514 (0.00497, 0.00529) |
| noncoding | tau_r | 18 | 0.00492 (0.00474, 0.00509) | 0.00493 (0.00475, 0.00510) | 0.00489 (0.00472, 0.00506) | 0.00493 (0.00476, 0.00510) |
| noncoding | tau_r | 19 | 0.00546 (0.00530, 0.00562) | 0.00545 (0.00529, 0.00561) | 0.00543 (0.00526, 0.00559) | 0.00545 (0.00529, 0.00560) |
| noncoding | tau_r | 20 | 0.00503 (0.00488, 0.00519) | 0.00504 (0.00489, 0.00519) | 0.00499 (0.00484, 0.00514) | 0.00504 (0.00489, 0.00519) |
| noncoding | tau_r | 21 | 0.00467 (0.00442, 0.00493) | 0.00470 (0.00447, 0.00493) | 0.00470 (0.00447, 0.00492) | 0.00471 (0.00448, 0.00493) |
| noncoding | tau_s | 1 | 0.00089 (8e-04, 0.00098) | 0.00472 (0.00448, 0.00494) | 0.00170 (0.00149, 0.00191) | 0.00471 (0.00448, 0.00493) |
| noncoding | tau_s | 2 | 6e-05 (2e-05, 1e-04) | 0.00559 (0.00486, 0.00629) | 0.00284 (0.00227, 0.00339) | 0.00556 (0.00479, 0.00629) |
| noncoding | tau_s | 3 | 0.00075 (0.00057, 0.00094) | 0.00583 (0.00546, 0.00618) | 0.00194 (0.00161, 0.00228) | 0.00583 (0.00545, 0.00617) |
| noncoding | tau_s | 4 | 0.00107 (0.00089, 0.00125) | 0.00551 (0.00486, 0.00608) | 0.00282 (0.00232, 0.00334) | 0.00552 (0.00483, 0.00607) |
| noncoding | tau_s | 5 | 0.00112 (0.00095, 0.00128) | 0.00580 (0.00550, 0.00607) | 0.00209 (0.00156, 0.00263) | 0.00580 (0.00551, 0.00607) |
| noncoding | tau_s | 6 | 0.00142 (0.00126, 0.00158) | 0.00483 (0.00431, 0.00532) | 0.00277 (0.00252, 0.00302) | 0.00485 (0.00435, 0.00534) |
| noncoding | tau_s | 7 | 0.00124 (0.00111, 0.00137) | 0.00409 (0.00351, 0.00472) | 0.00218 (0.00196, 0.00240) | 0.00411 (0.00350, 0.00472) |
| noncoding | tau_s | 8 | 0.00144 (0.00127, 0.00160) | 0.00541 (0.00508, 0.00570) | 0.00223 (0.00160, 0.00281) | 0.00541 (0.00508, 0.00571) |
| noncoding | tau_s | 9 | 7e-05 (2e-05, 0.00012) | 0.00538 (0.00472, 0.00596) | 0.00305 (0.00251, 0.00355) | 0.00539 (0.00473, 0.00596) |
| noncoding | tau_s | 10 | 0.00178 (0.00161, 0.00195) | 0.00482 (0.00439, 0.00521) | 0.00252 (0.00226, 0.00277) | 0.00480 (0.00438, 0.00522) |
| noncoding | tau_s | 11 | 0.00109 (0.00097, 0.00121) | 0.00472 (0.00433, 0.00506) | 0.00133 (0.00109, 0.00160) | 0.00474 (0.00436, 0.00507) |
| noncoding | tau_s | 12 | 0.00093 (0.00082, 0.00105) | 0.00518 (0.00492, 0.00540) | 0.00235 (0.00209, 0.00260) | 0.00517 (0.00492, 0.00540) |
| noncoding | tau_s | 13 | 0.00160 (0.00148, 0.00172) | 0.00521 (0.00499, 0.00541) | 0.00161 (0.00147, 0.00177) | 0.00521 (0.00499, 0.00541) |
| noncoding | tau_s | 14 | 0.00081 (0.00065, 0.00098) | 0.00628 (0.00575, 0.00671) | 0.00232 (0.00193, 0.00270) | 0.00627 (0.00571, 0.00671) |
| noncoding | tau_s | 15 | 0.00179 (0.00161, 0.00197) | 0.00476 (0.00418, 0.00531) | 0.00278 (0.00241, 0.00314) | 0.00476 (0.00419, 0.00532) |
| noncoding | tau_s | 16 | 1e-04 (3e-05, 0.00016) | 0.00473 (0.00398, 0.00545) | 0.00260 (0.00216, 0.00303) | 0.00470 (0.00394, 0.00542) |
| noncoding | tau_s | 17 | 0.00168 (0.00155, 0.00181) | 0.00500 (0.00473, 0.00525) | 0.00239 (0.00217, 0.00260) | 0.00500 (0.00473, 0.00525) |
| noncoding | tau_s | 18 | 0.00184 (0.00164, 0.00204) | 0.00416 (0.00369, 0.00462) | 0.00229 (0.00202, 0.00256) | 0.00416 (0.00367, 0.00463) |
| noncoding | tau_s | 19 | 0.00172 (0.00160, 0.00184) | 0.00507 (0.00471, 0.00541) | 0.00266 (0.00240, 0.00292) | 0.00507 (0.00470, 0.00541) |
| noncoding | tau_s | 20 | 0.00150 (0.00139, 0.00162) | 0.00476 (0.00445, 0.00504) | 0.00239 (0.00209, 0.00267) | 0.00476 (0.00445, 0.00504) |
| noncoding | tau_s | 21 | 0.00362 (0.00324, 0.00398) | 0.00430 (0.00392, 0.00464) | 0.00432 (0.00392, 0.00468) | 0.00436 (0.00399, 0.00472) |
| noncoding | tau_c | 1 | n/a | 0.00052 (0.00042, 0.00061) | 0.00021 (8e-05, 0.00033) | 0.00053 (0.00043, 0.00062) |
| noncoding | tau_c | 2 | n/a | 4e-05 (3e-05, 6e-05) | 2e-05 (0.00000, 4e-05) | 4e-05 (2e-05, 7e-05) |
| noncoding | tau_c | 3 | n/a | 0.00043 (0.00028, 0.00058) | 8e-05 (0.00000, 0.00016) | 0.00045 (0.00029, 6e-04) |
| noncoding | tau_c | 4 | n/a | 8e-04 (0.00058, 0.00101) | 0.00022 (4e-05, 0.00042) | 0.00083 (6e-04, 0.00106) |
| noncoding | tau_c | 5 | n/a | 0.00099 (0.00082, 0.00114) | 0.00044 (0.00000, 0.00077) | 0.00104 (0.00086, 0.00123) |
| noncoding | tau_c | 6 | n/a | 0.00042 (0.00029, 0.00055) | 6e-05 (1e-05, 0.00012) | 0.00043 (3e-04, 0.00056) |
| noncoding | tau_c | 7 | n/a | 4e-04 (0.00025, 0.00056) | 1e-04 (2e-05, 0.00019) | 0.00041 (0.00025, 0.00058) |
| noncoding | tau_c | 8 | n/a | 0.00106 (0.00088, 0.00123) | 0.00073 (0.00042, 0.00107) | 0.00110 (0.00091, 0.00128) |
| noncoding | tau_c | 9 | n/a | 3e-05 (2e-05, 5e-05) | 2e-05 (0.00000, 4e-05) | 4e-05 (2e-05, 7e-05) |
| noncoding | tau_c | 10 | n/a | 5e-05 (2e-05, 7e-05) | 1e-05 (0.00000, 3e-05) | 4e-05 (3e-05, 6e-05) |
| noncoding | tau_c | 11 | n/a | 0.00072 (0.00058, 0.00086) | 6e-04 (0.00036, 0.00082) | 0.00081 (0.00062, 0.00099) |
| noncoding | tau_c | 12 | n/a | 0.00041 (3e-04, 5e-04) | 0.00013 (4e-05, 0.00022) | 0.00041 (0.00031, 0.00051) |
| noncoding | tau_c | 13 | n/a | 0.00115 (0.00103, 0.00128) | 0.00147 (0.00131, 0.00160) | 0.00116 (0.00103, 0.00128) |
| noncoding | tau_c | 14 | n/a | 0.00058 (0.00041, 0.00074) | 8e-05 (1e-05, 0.00016) | 6e-04 (0.00042, 0.00077) |
| noncoding | tau_c | 15 | n/a | 0.00094 (0.00069, 0.00118) | 0.00064 (0.00039, 0.00088) | 0.00096 (0.00071, 0.00120) |
| noncoding | tau_c | 16 | n/a | 3e-05 (1e-05, 6e-05) | 2e-05 (0.00000, 4e-05) | 4e-05 (2e-05, 7e-05) |
| noncoding | tau_c | 17 | n/a | 0.00098 (8e-04, 0.00115) | 4e-04 (0.00022, 0.00057) | 0.00100 (0.00082, 0.00117) |
| noncoding | tau_c | 18 | n/a | 3e-05 (1e-05, 5e-05) | 1e-05 (0.00000, 3e-05) | 3e-05 (1e-05, 5e-05) |
| noncoding | tau_c | 19 | n/a | 0.00091 (0.00075, 0.00106) | 6e-04 (4e-04, 8e-04) | 0.00091 (0.00075, 0.00106) |
| noncoding | tau_c | 20 | n/a | 0.00082 (0.00069, 0.00094) | 0.00065 (0.00049, 0.00082) | 0.00082 (0.00069, 0.00095) |
| noncoding | tau_c | 21 | n/a | 9e-04 (6e-05, 0.00172) | 0.00092 (6e-05, 0.00174) | 0.00106 (0.00019, 0.00191) |
| noncoding | phi_c<-m | 1 | n/a | n/a | 0.13595 (0.07830, 0.19593) | 0.00728 (0.00000, 0.01937) |
| noncoding | phi_c<-m | 2 | n/a | n/a | 0.10401 (0.00518, 0.20372) | 0.00698 (0.00000, 0.02301) |
| noncoding | phi_c<-m | 3 | n/a | n/a | 0.09711 (0.00000, 0.19803) | 0.01299 (0.00000, 0.03940) |
| noncoding | phi_c<-m | 4 | n/a | n/a | 0.32695 (0.23205, 0.42542) | 0.02137 (0.00000, 0.06803) |



| | | | | | | |
|---|---|---|---|---|---|---|
| noncoding | phi_c<-m | 5 | n/a | n/a | 0.31102 (0.15996, 0.46466) | 0.04204 (0.00000, 0.11832) |
| noncoding | phi_c<-m | 6 | n/a | n/a | 0.02639 (0.01151, 0.04218) | 0.00282 (0.00000, 0.00841) |
| noncoding | phi_c<-m | 7 | n/a | n/a | 0.07026 (0.03709, 0.10615) | 0.00466 (0.00000, 0.01384) |
| noncoding | phi_c<-m | 8 | n/a | n/a | 0.24908 (0.12407, 0.38234) | 0.03137 (0.00000, 0.09080) |
| noncoding | phi_c<-m | 9 | n/a | n/a | 0.15048 (0.06718, 0.24245) | 0.01214 (0.00000, 0.03687) |
| noncoding | phi_c<-m | 10 | n/a | n/a | 0.04169 (0.00785, 0.07886) | 0.00386 (0.00000, 0.01209) |
| noncoding | phi_c<-m | 11 | n/a | n/a | 0.13794 (0.00000, 0.27567) | 0.05520 (0.00000, 0.12933) |
| noncoding | phi_c<-m | 12 | n/a | n/a | 0.10137 (0.05642, 0.15020) | 0.00281 (0.00000, 0.00859) |
| noncoding | phi_c<-m | 13 | n/a | n/a | 0.19210 (0.00176, 0.35197) | 0.00367 (0.00000, 0.01101) |
| noncoding | phi_c<-m | 14 | n/a | n/a | 0.18839 (0.10709, 0.27442) | 0.02317 (0.00000, 0.06168) |
| noncoding | phi_c<-m | 15 | n/a | n/a | 0.15730 (0.07160, 0.24443) | 0.01095 (0.00000, 0.03343) |
| noncoding | phi_c<-m | 16 | n/a | n/a | 0.15124 (0.04559, 0.25918) | 0.01604 (0.00000, 0.04798) |
| noncoding | phi_c<-m | 17 | n/a | n/a | 0.07873 (0.03629, 0.12391) | 0.00528 (0.00000, 0.01605) |
| noncoding | phi_c<-m | 18 | n/a | n/a | 0.05697 (0.01679, 0.10592) | 0.00448 (0.00000, 0.01414) |
| noncoding | phi_c<-m | 19 | n/a | n/a | 0.11574 (0.07158, 0.16181) | 0.00370 (0.00000, 0.01106) |
| noncoding | phi_c<-m | 20 | n/a | n/a | 0.17410 (0.11990, 0.23178) | 0.00442 (0.00000, 0.01314) |
| noncoding | phi_c<-m | 21 | n/a | n/a | 0.32184 (0.03771, 0.73888) | 0.49378 (0.01240, 0.96903) |
| noncoding | phi_m<-c | 1 | n/a | 0.51186 (0.47796, 0.54512) | n/a | 0.50635 (0.47215, 0.54122) |
| noncoding | phi_m<-c | 2 | n/a | 0.44214 (0.37537, 0.50479) | n/a | 0.43340 (0.36168, 0.50215) |
| noncoding | phi_m<-c | 3 | n/a | 0.53862 (0.49402, 0.58446) | n/a | 0.53205 (0.48471, 0.57871) |
| noncoding | phi_m<-c | 4 | n/a | 0.61804 (0.54779, 0.68624) | n/a | 0.60871 (0.53344, 0.68180) |
| noncoding | phi_m<-c | 5 | n/a | 0.64934 (0.60737, 0.69024) | n/a | 0.63273 (0.58153, 0.68286) |
| noncoding | phi_m<-c | 6 | n/a | 0.31478 (0.25730, 0.37125) | n/a | 0.31434 (0.25751, 0.37057) |
| noncoding | phi_m<-c | 7 | n/a | 0.36471 (0.29479, 0.43596) | n/a | 0.36202 (0.29172, 0.43635) |
| noncoding | phi_m<-c | 8 | n/a | 0.57075 (0.52290, 0.61786) | n/a | 0.55627 (0.50236, 0.61042) |
| noncoding | phi_m<-c | 9 | n/a | 0.42973 (0.36364, 0.49511) | n/a | 0.42014 (0.35121, 0.48919) |
| noncoding | phi_m<-c | 10 | n/a | 0.35381 (0.31389, 0.39544) | n/a | 0.34922 (0.30748, 0.38952) |
| noncoding | phi_m<-c | 11 | n/a | 0.57335 (0.52111, 0.62507) | n/a | 0.55867 (0.49750, 0.61703) |
| noncoding | phi_m<-c | 12 | n/a | 0.43229 (0.39754, 0.46689) | n/a | 0.43008 (0.39587, 0.46511) |
| noncoding | phi_m<-c | 13 | n/a | 0.50966 (0.47918, 0.54042) | n/a | 0.50719 (0.47672, 0.53863) |
| noncoding | phi_m<-c | 14 | n/a | 0.60629 (0.55528, 0.65595) | n/a | 0.58939 (0.53058, 0.64735) |
| noncoding | phi_m<-c | 15 | n/a | 0.38112 (0.30490, 0.45615) | n/a | 0.37351 (0.29823, 0.45334) |
| noncoding | phi_m<-c | 16 | n/a | 0.37992 (0.30787, 0.45231) | n/a | 0.36546 (0.28624, 0.44105) |
| noncoding | phi_m<-c | 17 | n/a | 0.43462 (0.38634, 0.47966) | n/a | 0.43319 (0.38545, 0.47877) |
| noncoding | phi_m<-c | 18 | n/a | 0.34645 (0.29424, 0.39884) | n/a | 0.34238 (0.28776, 0.39390) |
| noncoding | phi_m<-c | 19 | n/a | 0.39945 (0.35239, 0.44644) | n/a | 0.39598 (0.34791, 0.44359) |
| noncoding | phi_m<-c | 20 | n/a | 0.42318 (0.38219, 0.46390) | n/a | 0.41928 (0.37683, 0.46020) |
| noncoding | phi_m<-c | 21 | n/a | 0.30713 (0.03269, 0.73111) | n/a | 0.50759 (0.04163, 0.99984) |
| coding | theta_H | 1 | 0.00550 (0.00525, 0.00575) | 0.00552 (0.00527, 0.00577) | 0.00548 (0.00523, 0.00572) | 0.00552 (0.00527, 0.00577) |
| coding | theta_H | 2 | 0.01101 (0.01035, 0.01169) | 0.01106 (0.01041, 0.01174) | 0.01107 (0.01039, 0.01175) | 0.01106 (0.01039, 0.01173) |
| coding | theta_H | 3 | 0.00890 (0.00830, 0.00949) | 0.00897 (0.00838, 0.00958) | 0.00888 (0.00828, 0.00947) | 0.00897 (0.00837, 0.00956) |
| coding | theta_H | 4 | 0.00967 (0.00895, 0.01037) | 0.00970 (0.00898, 0.01042) | 0.00967 (0.00895, 0.01038) | 0.00969 (0.00898, 0.01040) |
| coding | theta_H | 5 | 0.00839 (0.00784, 0.00895) | 0.00848 (0.00793, 0.00904) | 0.00838 (0.00782, 0.00893) | 0.00848 (0.00792, 0.00904) |
| coding | theta_H | 6 | 0.00668 (0.00635, 0.00702) | 0.00667 (0.00633, 0.00700) | 0.00666 (0.00632, 0.00699) | 0.00667 (0.00634, 0.00701) |
| coding | theta_H | 7 | 0.00615 (0.00583, 0.00646) | 0.00614 (0.00582, 0.00646) | 0.00614 (0.00582, 0.00645) | 0.00614 (0.00582, 0.00645) |
| coding | theta_H | 8 | 0.00872 (0.00819, 0.00927) | 0.00876 (0.00823, 0.00931) | 0.00870 (0.00817, 0.00924) | 0.00877 (0.00823, 0.00931) |
| coding | theta_H | 9 | 0.01156 (0.01070, 0.01245) | 0.01164 (0.01077, 0.01254) | 0.01163 (0.01074, 0.01252) | 0.01164 (0.01077, 0.01255) |
| coding | theta_H | 10 | 0.00582 (0.00556, 0.00607) | 0.00581 (0.00556, 0.00607) | 0.00580 (0.00555, 0.00606) | 0.00581 (0.00556, 0.00607) |
| coding | theta_H | 11 | 0.00704 (0.00668, 0.00741) | 0.00706 (0.00669, 0.00742) | 0.00703 (0.00666, 0.00739) | 0.00706 (0.00670, 0.00743) |
| coding | theta_H | 12 | 0.00691 (0.00660, 0.00722) | 0.00695 (0.00664, 0.00726) | 0.00686 (0.00655, 0.00717) | 0.00695 (0.00664, 0.00726) |
| coding | theta_H | 13 | 0.00606 (0.00576, 0.00636) | 0.00613 (0.00582, 0.00642) | 0.00606 (0.00575, 0.00636) | 0.00613 (0.00583, 0.00643) |
| coding | theta_H | 14 | 0.01038 (0.00971, 0.01104) | 0.01041 (0.00974, 0.01107) | 0.01038 (0.00972, 0.01105) | 0.01041 (0.00976, 0.01108) |
| coding | theta_H | 15 | 0.00818 (0.00766, 0.00868) | 0.00818 (0.00768, 0.00870) | 0.00816 (0.00765, 0.00867) | 0.00818 (0.00767, 0.00869) |
| coding | theta_H | 16 | 0.00920 (0.00856, 0.00984) | 0.00927 (0.00864, 0.00994) | 0.00927 (0.00864, 0.00994) | 0.00927 (0.00863, 0.00992) |
| coding | theta_H | 17 | 0.00587 (0.00558, 0.00615) | 0.00590 (0.00561, 0.00618) | 0.00585 (0.00557, 0.00614) | 0.00590 (0.00562, 0.00619) |
| coding | theta_H | 18 | 0.00582 (0.00550, 0.00613) | 0.00581 (0.00550, 0.00612) | 0.00581 (0.00549, 0.00611) | 0.00581 (0.00550, 0.00613) |
| coding | theta_H | 19 | 0.00535 (0.00511, 0.00559) | 0.00535 (0.00510, 0.00559) | 0.00534 (0.00510, 0.00558) | 0.00535 (0.00510, 0.00559) |
| coding | theta_H | 20 | 0.00516 (0.00492, 0.00540) | 0.00517 (0.00493, 0.00541) | 0.00515 (0.00491, 0.00538) | 0.00517 (0.00493, 0.00541) |
| coding | theta_H | 21 | n/a | n/a | n/a | n/a |
| coding | theta_C | 1 | 0.00541 (0.00482, 0.00600) | 0.03614 (0.02031, 0.05447) | 0.03068 (0.01327, 0.05128) | 0.03630 (0.02036, 0.05526) |
| coding | theta_C | 2 | 0.00305 (0.00079, 0.00581) | 0.02116 (0.00691, 0.03807) | 0.01516 (0.00309, 0.03058) | 0.02142 (0.00686, 0.03878) |
| coding | theta_C | 3 | 0.00756 (0.00586, 0.00930) | 0.03671 (0.01890, 0.05630) | 0.01949 (0.00485, 0.03696) | 0.03719 (0.01957, 0.05754) |
| coding | theta_C | 4 | 0.01213 (0.00958, 0.01494) | 0.03092 (0.01828, 0.04606) | 0.01691 (0.00489, 0.03145) | 0.03045 (0.01805, 0.04530) |
| coding | theta_C | 5 | 0.01497 (0.01182, 0.01826) | 0.04031 (0.02528, 0.05708) | 0.02309 (0.00876, 0.03969) | 0.03875 (0.02465, 0.05517) |
| coding | theta_C | 6 | 0.00634 (0.00569, 0.00700) | 0.02216 (0.01135, 0.03584) | 0.01748 (0.00383, 0.03402) | 0.02197 (0.01149, 0.03605) |
| coding | theta_C | 7 | 0.00635 (0.00565, 0.00706) | 0.02525 (0.01256, 0.04130) | 0.02075 (0.00643, 0.03798) | 0.02515 (0.01253, 0.04087) |
| coding | theta_C | 8 | 0.00986 (0.00851, 0.01122) | 0.02481 (0.01638, 0.03463) | 0.02886 (0.01426, 0.04621) | 0.02429 (0.01634, 0.03390) |
| coding | theta_C | 9 | 0.00205 (0.00062, 0.00370) | 0.01716 (0.00455, 0.03319) | 0.01330 (0.00180, 0.02795) | 0.01766 (0.00434, 0.03352) |
| coding | theta_C | 10 | 0.00809 (0.00738, 0.00883) | 0.02062 (0.00697, 0.03770) | 0.01436 (0.00242, 0.02924) | 0.02036 (0.00651, 0.03744) |
| coding | theta_C | 11 | 0.00701 (0.00616, 0.00788) | 0.02062 (0.01286, 0.03017) | 0.03166 (0.01524, 0.05040) | 0.01905 (0.01218, 0.02759) |
| coding | theta_C | 12 | 0.00604 (0.00534, 0.00674) | 0.03882 (0.02167, 0.05822) | 0.02481 (0.00907, 0.04254) | 0.03882 (0.02216, 0.05821) |
| coding | theta_C | 13 | 0.00865 (0.00780, 0.00954) | 0.03276 (0.02183, 0.04579) | 0.04362 (0.02573, 0.06305) | 0.03267 (0.02159, 0.04549) |

| | | | | | | |
|---|---|---|---|---|---|---|
| **coding** | **theta_C** | **14** | 0.01072 (0.00840, 0.01311) | 0.03499 (0.01983, 0.05264) | 0.01957 (0.00511, 0.03659) | 0.03493 (0.01950, 0.05202) |
| **coding** | **theta_C** | **15** | 0.01060 (0.00932, 0.01192) | 0.03324 (0.01989, 0.04887) | 0.03394 (0.01825, 0.05208) | 0.03311 (0.01997, 0.04880) |
| **coding** | **theta_C** | **16** | 0.00188 (0.00072, 0.00308) | 0.01599 (0.00380, 0.03141) | 0.01388 (0.00233, 0.02905) | 0.01663 (0.00412, 0.03220) |
| **coding** | **theta_C** | **17** | 0.00613 (0.00562, 0.00664) | 0.01626 (0.01173, 0.02148) | 0.03195 (0.01645, 0.05032) | 0.01616 (0.01166, 0.02125) |
| **coding** | **theta_C** | **18** | 0.00739 (0.00666, 0.00814) | 0.01579 (0.00347, 0.03163) | 0.01300 (0.00173, 0.02773) | 0.01565 (0.00372, 0.03128) |
| **coding** | **theta_C** | **19** | 0.00763 (0.00699, 0.00828) | 0.02235 (0.01546, 0.03059) | 0.02683 (0.01549, 0.04090) | 0.02234 (0.01536, 0.03049) |
| **coding** | **theta_C** | **20** | 0.00713 (0.00648, 0.00778) | 0.03460 (0.02040, 0.05102) | 0.03804 (0.02098, 0.05742) | 0.03471 (0.02069, 0.05138) |
| **coding** | **theta_C** | **21** | n/a | n/a | n/a | n/a |
| **coding** | **theta_M** | **1** | 0.00160 (0.00145, 0.00176) | 0.00096 (8e-04, 0.00110) | 0.00052 (0.00026, 0.00077) | 0.00096 (0.00081, 0.00111) |
| **coding** | **theta_M** | **2** | 3e-05 (1e-05, 5e-05) | 2e-05 (1e-05, 3e-05) | 1e-05 (0.00000, 2e-05) | 2e-05 (1e-05, 3e-05) |
| **coding** | **theta_M** | **3** | 0.00155 (0.00120, 0.00190) | 0.00088 (0.00059, 0.00117) | 0.00021 (2e-05, 4e-04) | 0.00091 (0.00061, 0.00120) |
| **coding** | **theta_M** | **4** | 0.00417 (0.00348, 0.00489) | 0.00307 (0.00236, 0.00380) | 0.00163 (0.00045, 0.00284) | 0.00312 (0.00240, 0.00383) |
| **coding** | **theta_M** | **5** | 0.00895 (0.00746, 0.01050) | 0.00725 (0.00601, 0.00856) | 0.01316 (0.00566, 0.02343) | 0.00723 (0.00605, 0.00847) |
| **coding** | **theta_M** | **6** | 0.00108 (0.00098, 0.00118) | 0.00043 (0.00032, 0.00054) | 9e-05 (2e-05, 0.00018) | 0.00044 (0.00033, 0.00055) |
| **coding** | **theta_M** | **7** | 0.00182 (0.00164, 0.00199) | 0.00075 (0.00052, 0.00098) | 0.00027 (7e-05, 0.00047) | 0.00076 (0.00052, 0.00100) |
| **coding** | **theta_M** | **8** | 0.00461 (0.00408, 0.00516) | 0.00335 (0.00285, 0.00383) | 0.00327 (0.00236, 0.00418) | 0.00340 (0.00291, 0.00391) |
| **coding** | **theta_M** | **9** | 4e-05 (1e-05, 6e-05) | 2e-05 (1e-05, 3e-05) | 1e-05 (0.00000, 2e-05) | 2e-05 (1e-05, 4e-05) |
| **coding** | **theta_M** | **10** | 0.00016 (0.00014, 0.00018) | 2e-05 (1e-05, 2e-05) | 1e-05 (0.00000, 1e-05) | 2e-05 (1e-05, 2e-05) |
| **coding** | **theta_M** | **11** | 0.00344 (0.00306, 0.00381) | 0.00227 (0.00190, 0.00265) | 0.00223 (0.00158, 0.00290) | 0.00241 (0.00199, 0.00284) |
| **coding** | **theta_M** | **12** | 0.00123 (0.00109, 0.00136) | 0.00058 (0.00046, 0.00071) | 0.00024 (9e-05, 0.00038) | 0.00059 (0.00046, 0.00071) |
| **coding** | **theta_M** | **13** | 0.00561 (0.00512, 0.00610) | 0.00423 (0.00379, 0.00468) | 0.00540 (0.00488, 0.00596) | 0.00423 (0.00378, 0.00467) |
| **coding** | **theta_M** | **14** | 0.00163 (0.00132, 0.00192) | 0.00115 (0.00085, 0.00145) | 0.00023 (5e-05, 0.00044) | 0.00118 (0.00087, 0.00149) |
| **coding** | **theta_M** | **15** | 0.00387 (0.00347, 0.00425) | 0.00240 (0.00194, 0.00286) | 0.00192 (0.00137, 0.00248) | 0.00242 (0.00196, 0.00287) |
| **coding** | **theta_M** | **16** | 9e-05 (3e-05, 0.00014) | 3e-05 (1e-05, 5e-05) | 2e-05 (0.00000, 4e-05) | 4e-05 (1e-05, 6e-05) |
| **coding** | **theta_M** | **17** | 0.00306 (0.00282, 0.00330) | 0.00191 (0.00163, 0.00217) | 0.00107 (7e-04, 0.00145) | 0.00192 (0.00165, 0.00219) |
| **coding** | **theta_M** | **18** | 9e-05 (5e-05, 0.00012) | 1e-05 (0.00000, 2e-05) | 0.00000 (0.00000, 1e-05) | 1e-05 (1e-05, 2e-05) |
| **coding** | **theta_M** | **19** | 0.00304 (0.00282, 0.00326) | 0.00193 (0.00168, 0.00217) | 0.00171 (0.00134, 0.00209) | 0.00194 (0.00170, 0.00218) |
| **coding** | **theta_M** | **20** | 0.00282 (0.00260, 0.00304) | 0.00175 (0.00153, 0.00197) | 0.00168 (0.00137, 0.00198) | 0.00176 (0.00153, 0.00197) |
| **coding** | **theta_M** | **21** | n/a | n/a | n/a | n/a |
| **coding** | **theta_r** | **1** | 0.00922 (0.00884, 0.00962) | 0.00922 (0.00881, 0.00963) | 0.00941 (0.00900, 0.00981) | 0.00922 (0.00882, 0.00963) |
| **coding** | **theta_r** | **2** | 0.01300 (0.01235, 0.01363) | 0.01335 (0.01269, 0.01404) | 0.01338 (0.01270, 0.01404) | 0.01335 (0.01267, 0.01402) |
| **coding** | **theta_r** | **3** | 0.01022 (0.00962, 0.01082) | 0.01015 (0.00954, 0.01077) | 0.01048 (0.00985, 0.01109) | 0.01016 (0.00954, 0.01077) |
| **coding** | **theta_r** | **4** | 0.00972 (0.00909, 0.01038) | 0.00977 (0.00911, 0.01043) | 0.00989 (0.00923, 0.01056) | 0.00977 (0.00911, 0.01043) |
| **coding** | **theta_r** | **5** | 0.01038 (0.00976, 0.01097) | 0.01021 (0.00961, 0.01081) | 0.01040 (0.00979, 0.01100) | 0.01022 (0.00962, 0.01083) |
| **coding** | **theta_r** | **6** | 0.00989 (0.00942, 0.01035) | 0.01002 (0.00953, 0.01048) | 0.01013 (0.00964, 0.01060) | 0.01001 (0.00954, 0.01050) |
| **coding** | **theta_r** | **7** | 0.00938 (0.00892, 0.00984) | 0.00953 (0.00906, 0.01001) | 0.00955 (0.00908, 0.01003) | 0.00953 (0.00904, 0.01000) |
| **coding** | **theta_r** | **8** | 0.01014 (0.00961, 0.01069) | 0.01008 (0.00954, 0.01062) | 0.01022 (0.00966, 0.01077) | 0.01008 (0.00954, 0.01062) |
| **coding** | **theta_r** | **9** | 0.01156 (0.01087, 0.01226) | 0.01184 (0.01113, 0.01257) | 0.01189 (0.01117, 0.01264) | 0.01185 (0.01114, 0.01258) |
| **coding** | **theta_r** | **10** | 0.00873 (0.00833, 0.00911) | 0.00881 (0.00842, 0.00921) | 0.00890 (0.00850, 0.00931) | 0.00881 (0.00841, 0.00921) |
| **coding** | **theta_r** | **11** | 0.01052 (0.01006, 0.01099) | 0.01052 (0.01004, 0.01099) | 0.01057 (0.01010, 0.01104) | 0.01052 (0.01005, 0.01101) |
| **coding** | **theta_r** | **12** | 0.01062 (0.01020, 0.01104) | 0.01058 (0.01012, 0.01100) | 0.01092 (0.01048, 0.01136) | 0.01058 (0.01014, 0.01102) |
| **coding** | **theta_r** | **13** | 0.00903 (0.00861, 0.00946) | 0.00880 (0.00839, 0.00922) | 0.00902 (0.00860, 0.00945) | 0.00879 (0.00838, 0.00921) |
| **coding** | **theta_r** | **14** | 0.01081 (0.01020, 0.01144) | 0.01084 (0.01019, 0.01146) | 0.01102 (0.01040, 0.01167) | 0.01084 (0.01020, 0.01147) |
| **coding** | **theta_r** | **15** | 0.01138 (0.01082, 0.01196) | 0.01145 (0.01086, 0.01203) | 0.01151 (0.01094, 0.01209) | 0.01145 (0.01088, 0.01205) |
| **coding** | **theta_r** | **16** | 0.01068 (0.01004, 0.01132) | 0.01098 (0.01032, 0.01164) | 0.01097 (0.01031, 0.01164) | 0.01098 (0.01030, 0.01164) |
| **coding** | **theta_r** | **17** | 0.00895 (0.00853, 0.00935) | 0.00883 (0.00839, 0.00924) | 0.00909 (0.00868, 0.00951) | 0.00882 (0.00840, 0.00923) |
| **coding** | **theta_r** | **18** | 0.00890 (0.00842, 0.00934) | 0.00888 (0.00840, 0.00933) | 0.00896 (0.00850, 0.00943) | 0.00888 (0.00842, 0.00935) |
| **coding** | **theta_r** | **19** | 0.00978 (0.00934, 0.01021) | 0.00986 (0.00942, 0.01030) | 0.00989 (0.00944, 0.01033) | 0.00986 (0.00941, 0.01029) |
| **coding** | **theta_r** | **20** | 0.00882 (0.00843, 0.00921) | 0.00885 (0.00844, 0.00924) | 0.00896 (0.00856, 0.00936) | 0.00885 (0.00845, 0.00925) |
| **coding** | **theta_r** | **21** | 0.00789 (0.00726, 0.00851) | 0.00787 (0.00730, 0.00845) | 0.00787 (0.00728, 0.00845) | 0.00785 (0.00727, 0.00842) |
| **coding** | **theta_s** | **1** | 0.01174 (0.01108, 0.01241) | 0.00274 (0.00038, 0.00538) | 0.00921 (0.00840, 0.01002) | 0.00272 (0.00035, 0.00533) |
| **coding** | **theta_s** | **2** | 0.03018 (0.02809, 0.03230) | 0.00808 (0.00286, 0.01307) | 0.01574 (0.01326, 0.01816) | 0.00819 (0.00286, 0.01307) |
| **coding** | **theta_s** | **3** | 0.01872 (0.01717, 0.02031) | 0.00795 (0.00066, 0.01853) | 0.01336 (0.01181, 0.01486) | 0.00792 (0.00056, 0.01837) |
| **coding** | **theta_s** | **4** | 0.01544 (0.01406, 0.01680) | 0.00732 (0.00135, 0.01368) | 0.01140 (0.00954, 0.01324) | 0.00738 (0.00135, 0.01400) |
| **coding** | **theta_s** | **5** | 0.01616 (0.01490, 0.01742) | 0.01232 (0.00128, 0.02663) | 0.01534 (0.01350, 0.01719) | 0.01222 (0.00126, 0.02633) |
| **coding** | **theta_s** | **6** | 0.01681 (0.01543, 0.01819) | 0.00601 (0.00234, 0.00950) | 0.01055 (0.00925, 0.01188) | 0.00590 (0.00219, 0.00938) |
| **coding** | **theta_s** | **7** | 0.01349 (0.01252, 0.01446) | 0.00737 (0.00467, 0.00991) | 0.01015 (0.00916, 0.01114) | 0.00731 (0.00456, 0.00980) |
| **coding** | **theta_s** | **8** | 0.01527 (0.01394, 0.01656) | 0.01154 (0.00123, 0.02495) | 0.01361 (0.01140, 0.01588) | 0.01157 (0.00097, 0.02508) |
| **coding** | **theta_s** | **9** | 0.02563 (0.02360, 0.02775) | 0.00564 (0.00126, 0.00999) | 0.01242 (0.01008, 0.01476) | 0.00555 (0.00135, 0.00993) |
| **coding** | **theta_s** | **10** | 0.01170 (0.01070, 0.01273) | 0.00371 (0.00153, 0.00598) | 0.00892 (0.00787, 0.00996) | 0.00380 (0.00155, 0.00604) |
| **coding** | **theta_s** | **11** | 0.01317 (0.01229, 0.01407) | 0.00738 (0.00083, 0.01548) | 0.01251 (0.01132, 0.01367) | 0.00738 (0.00097, 0.01594) |
| **coding** | **theta_s** | **12** | 0.01729 (0.01620, 0.01834) | 0.00400 (0.00049, 0.00811) | 0.01111 (0.00994, 0.01227) | 0.00396 (0.00057, 0.00808) |
| **coding** | **theta_s** | **13** | 0.01247 (0.01158, 0.01336) | 0.01200 (0.00123, 0.02626) | 0.01329 (0.01231, 0.01428) | 0.01201 (0.00112, 0.02619) |
| **coding** | **theta_s** | **14** | 0.01920 (0.01784, 0.02059) | 0.00808 (0.00094, 0.01660) | 0.01443 (0.01295, 0.01595) | 0.00785 (0.00095, 0.01605) |
| **coding** | **theta_s** | **15** | 0.01632 (0.01473, 0.01791) | 0.00766 (0.00270, 0.01221) | 0.01234 (0.01042, 0.01423) | 0.00766 (0.00260, 0.01221) |
| **coding** | **theta_s** | **16** | 0.02462 (0.02276, 0.02652) | 0.00800 (0.00378, 0.01211) | 0.01301 (0.01097, 0.01499) | 0.00813 (0.00398, 0.01209) |
| **coding** | **theta_s** | **17** | 0.01290 (0.01190, 0.01390) | 0.00647 (0.00035, 0.01602) | 0.00993 (0.00881, 0.01106) | 0.00652 (0.00048, 0.01614) |
| **coding** | **theta_s** | **18** | 0.00966 (0.00860, 0.01075) | 0.00487 (0.00237, 0.00716) | 0.00837 (0.00729, 0.00946) | 0.00485 (0.00237, 0.00728) |
| **coding** | **theta_s** | **19** | 0.01337 (0.01245, 0.01432) | 0.00386 (0.00094, 0.00674) | 0.01046 (0.00927, 0.01167) | 0.00386 (0.00102, 0.00680) |
| **coding** | **theta_s** | **20** | 0.01267 (0.01177, 0.01356) | 0.00410 (0.00093, 0.00743) | 0.00993 (0.00865, 0.01120) | 0.00403 (0.00078, 0.00742) |
| **coding** | **theta_s** | **21** | 0.00154 (7e-04, 0.00241) | 6e-04 (0.00015, 0.00117) | 0.00057 (1e-04, 0.00115) | 0.00052 (7e-05, 0.00110) |
| **coding** | **theta_c** | **1** | n/a | 0.00592 (0.00553, 0.00632) | 0.00438 (0.00324, 0.00548) | 0.00577 (0.00528, 0.00624) |



| | | | | | | |
|---|---|---|---|---|---|---|
| **coding** | **theta_c** | **2** | n/a | 0.01875 (0.01714, 0.02040) | 0.01842 (0.01329, 0.02355) | 0.01848 (0.01675, 0.02026) |
| **coding** | **theta_c** | **3** | n/a | 0.00980 (0.00892, 0.01069) | 0.00918 (0.00644, 0.01181) | 0.00938 (0.00817, 0.01052) |
| **coding** | **theta_c** | **4** | n/a | 0.00984 (0.00871, 0.01100) | 0.00687 (0.00452, 0.00919) | 0.00925 (0.00750, 0.01087) |
| **coding** | **theta_c** | **5** | n/a | 0.00966 (0.00875, 0.01057) | 0.00702 (0.00331, 0.01108) | 0.00850 (0.00620, 0.01025) |
| **coding** | **theta_c** | **6** | n/a | 0.00822 (0.00758, 0.00886) | 0.00821 (0.00748, 0.00892) | 0.00814 (0.00749, 0.00881) |
| **coding** | **theta_c** | **7** | n/a | 0.00739 (0.00673, 0.00805) | 0.00660 (0.00569, 0.00752) | 0.00728 (0.00658, 0.00794) |
| **coding** | **theta_c** | **8** | n/a | 0.00843 (0.00765, 0.00925) | 0.00393 (0.00156, 0.00624) | 0.00761 (0.00584, 0.00901) |
| **coding** | **theta_c** | **9** | n/a | 0.01472 (0.01334, 0.01611) | 0.01211 (0.00888, 0.01527) | 0.01428 (0.01272, 0.01594) |
| **coding** | **theta_c** | **10** | n/a | 0.00908 (0.00859, 0.00956) | 0.00837 (0.00739, 0.00929) | 0.00899 (0.00847, 0.00952) |
| **coding** | **theta_c** | **11** | n/a | 0.00741 (0.00680, 0.00803) | 0.00289 (0.00118, 0.00462) | 0.00614 (0.00433, 0.00777) |
| **coding** | **theta_c** | **12** | n/a | 0.00822 (0.00769, 0.00875) | 0.00696 (0.00577, 0.00807) | 0.00814 (0.00760, 0.00870) |
| **coding** | **theta_c** | **13** | n/a | 0.00604 (0.00559, 0.00649) | 0.00034 (4e-05, 0.00081) | 0.00596 (0.00549, 0.00644) |
| **coding** | **theta_c** | **14** | n/a | 0.01163 (0.01063, 0.01269) | 0.00956 (0.00705, 0.01195) | 0.01086 (0.00917, 0.01239) |
| **coding** | **theta_c** | **15** | n/a | 0.00904 (0.00799, 0.01013) | 0.00625 (0.00403, 0.00845) | 0.00873 (0.00748, 0.00995) |
| **coding** | **theta_c** | **16** | n/a | 0.01325 (0.01201, 0.01442) | 0.00953 (0.00656, 0.01240) | 0.01271 (0.01113, 0.01423) |
| **coding** | **theta_c** | **17** | n/a | 0.00577 (0.00531, 0.00624) | 0.00464 (0.00360, 0.00564) | 0.00564 (0.00512, 0.00617) |
| **coding** | **theta_c** | **18** | n/a | 0.00793 (0.00740, 0.00845) | 0.00688 (0.00586, 0.00785) | 0.00783 (0.00730, 0.00843) |
| **coding** | **theta_c** | **19** | n/a | 0.00701 (0.00646, 0.00755) | 0.00505 (0.00390, 0.00622) | 0.00692 (0.00637, 0.00751) |
| **coding** | **theta_c** | **20** | n/a | 0.00622 (0.00575, 0.00671) | 0.00346 (0.00244, 0.00448) | 0.00613 (0.00559, 0.00664) |
| **coding** | **theta_c** | **21** | n/a | 0.01326 (0.00049, 0.02916) | 0.00994 (2e-04, 0.02363) | 0.01392 (0.00074, 0.02982) |
| **coding** | **theta_m** | **1** | n/a | 0.01186 (0.00764, 0.01684) | 0.01052 (0.00716, 0.01443) | 0.01285 (0.00774, 0.01890) |
| **coding** | **theta_m** | **2** | n/a | 0.00837 (0.00067, 0.01928) | 0.02025 (0.00812, 0.03512) | 0.01150 (0.00109, 0.02548) |
| **coding** | **theta_m** | **3** | n/a | 0.02248 (0.01182, 0.03529) | 0.02500 (0.01333, 0.03935) | 0.02642 (0.01258, 0.04288) |
| **coding** | **theta_m** | **4** | n/a | 0.00955 (0.00512, 0.01472) | 0.01189 (0.00931, 0.01474) | 0.01224 (0.00532, 0.02211) |
| **coding** | **theta_m** | **5** | n/a | 0.00878 (0.00483, 0.01332) | 0.01040 (0.00769, 0.01322) | 0.01539 (0.00522, 0.03242) |
| **coding** | **theta_m** | **6** | n/a | 0.00633 (0.00424, 0.00863) | 0.00466 (0.00378, 0.00561) | 0.00652 (0.00433, 0.00897) |
| **coding** | **theta_m** | **7** | n/a | 0.00688 (0.00477, 0.00916) | 0.00748 (0.00573, 0.00938) | 0.00711 (0.00486, 0.00957) |
| **coding** | **theta_m** | **8** | n/a | 0.01368 (0.00779, 0.02062) | 0.01323 (0.00823, 0.01931) | 0.01897 (0.00836, 0.03370) |
| **coding** | **theta_m** | **9** | n/a | 0.01822 (0.00617, 0.03263) | 0.01581 (0.00742, 0.02590) | 0.02056 (0.00726, 0.03662) |
| **coding** | **theta_m** | **10** | n/a | 0.01701 (0.00390, 0.03312) | 0.01254 (0.00381, 0.02342) | 0.02008 (0.00570, 0.03780) |
| **coding** | **theta_m** | **11** | n/a | 0.01017 (0.00639, 0.01451) | 0.01999 (0.00881, 0.03413) | 0.02309 (0.00670, 0.04651) |
| **coding** | **theta_m** | **12** | n/a | 0.01271 (0.00826, 0.01784) | 0.01266 (0.00861, 0.01736) | 0.01329 (0.00845, 0.01879) |
| **coding** | **theta_m** | **13** | n/a | 0.00798 (0.00574, 0.01038) | 0.01082 (0.00196, 0.02381) | 0.00826 (0.00590, 0.01084) |
| **coding** | **theta_m** | **14** | n/a | 0.01480 (0.00756, 0.02350) | 0.01327 (0.00924, 0.01775) | 0.01777 (0.00836, 0.02951) |
| **coding** | **theta_m** | **15** | n/a | 0.01125 (0.00709, 0.01621) | 0.01775 (0.01172, 0.02492) | 0.01212 (0.00728, 0.01782) |
| **coding** | **theta_m** | **16** | n/a | 0.02264 (0.00868, 0.03850) | 0.02669 (0.01434, 0.04141) | 0.02575 (0.01028, 0.04349) |
| **coding** | **theta_m** | **17** | n/a | 0.01523 (0.00964, 0.02182) | 0.01474 (0.01010, 0.02028) | 0.01623 (0.01000, 0.02383) |
| **coding** | **theta_m** | **18** | n/a | 0.01152 (0.00097, 0.02512) | 0.00981 (0.00285, 0.01921) | 0.01441 (0.00185, 0.03000) |
| **coding** | **theta_m** | **19** | n/a | 0.00760 (0.00563, 0.00969) | 0.00927 (0.00716, 0.01153) | 0.00776 (0.00577, 0.00998) |
| **coding** | **theta_m** | **20** | n/a | 0.00881 (0.00620, 0.01186) | 0.01208 (0.00888, 0.01576) | 0.00914 (0.00624, 0.01230) |
| **coding** | **theta_m** | **21** | n/a | 0.01000 (0.00026, 0.02389) | 0.01372 (0.00069, 0.02998) | 0.01367 (0.00049, 0.02945) |
| **coding** | **tau_r** | **1** | 0.00487 (0.00472, 0.00501) | 0.00489 (0.00474, 0.00504) | 0.00480 (0.00465, 0.00495) | 0.00489 (0.00474, 0.00503) |
| **coding** | **tau_r** | **2** | 0.00644 (0.00620, 0.00667) | 0.00638 (0.00614, 0.00661) | 0.00636 (0.00613, 0.00660) | 0.00638 (0.00613, 0.00660) |
| **coding** | **tau_r** | **3** | 0.00593 (0.00570, 0.00616) | 0.00600 (0.00576, 0.00623) | 0.00584 (0.00561, 0.00607) | 0.00600 (0.00576, 0.00623) |
| **coding** | **tau_r** | **4** | 0.00595 (0.00569, 0.00619) | 0.00596 (0.00571, 0.00620) | 0.00590 (0.00564, 0.00614) | 0.00596 (0.00570, 0.00620) |
| **coding** | **tau_r** | **5** | 0.00579 (0.00557, 0.00602) | 0.00589 (0.00567, 0.00611) | 0.00579 (0.00556, 0.00601) | 0.00589 (0.00567, 0.00611) |
| **coding** | **tau_r** | **6** | 0.00551 (0.00533, 0.00569) | 0.00548 (0.00529, 0.00565) | 0.00543 (0.00525, 0.00561) | 0.00548 (0.00530, 0.00565) |
| **coding** | **tau_r** | **7** | 0.00532 (0.00514, 0.00549) | 0.00528 (0.00510, 0.00545) | 0.00526 (0.00508, 0.00543) | 0.00528 (0.00510, 0.00545) |
| **coding** | **tau_r** | **8** | 0.00549 (0.00528, 0.00569) | 0.00554 (0.00534, 0.00574) | 0.00546 (0.00525, 0.00566) | 0.00554 (0.00533, 0.00573) |
| **coding** | **tau_r** | **9** | 0.00601 (0.00575, 0.00627) | 0.00597 (0.00572, 0.00623) | 0.00595 (0.00568, 0.00621) | 0.00597 (0.00570, 0.00622) |
| **coding** | **tau_r** | **10** | 0.00542 (0.00525, 0.00557) | 0.00540 (0.00524, 0.00555) | 0.00535 (0.00519, 0.00551) | 0.00540 (0.00524, 0.00556) |
| **coding** | **tau_r** | **11** | 0.00494 (0.00478, 0.00511) | 0.00496 (0.00479, 0.00512) | 0.00492 (0.00475, 0.00509) | 0.00496 (0.00478, 0.00512) |
| **coding** | **tau_r** | **12** | 0.00529 (0.00513, 0.00545) | 0.00533 (0.00517, 0.00549) | 0.00518 (0.00501, 0.00533) | 0.00533 (0.00517, 0.00549) |
| **coding** | **tau_r** | **13** | 0.00516 (0.00499, 0.00532) | 0.00528 (0.00511, 0.00543) | 0.00516 (0.00499, 0.00532) | 0.00528 (0.00510, 0.00544) |
| **coding** | **tau_r** | **14** | 0.00654 (0.00630, 0.00677) | 0.00656 (0.00632, 0.00679) | 0.00647 (0.00624, 0.00671) | 0.00655 (0.00631, 0.00678) |
| **coding** | **tau_r** | **15** | 0.00542 (0.00522, 0.00563) | 0.00542 (0.00521, 0.00562) | 0.00538 (0.00517, 0.00558) | 0.00542 (0.00521, 0.00562) |
| **coding** | **tau_r** | **16** | 0.00586 (0.00562, 0.00610) | 0.00582 (0.00557, 0.00605) | 0.00581 (0.00557, 0.00605) | 0.00582 (0.00558, 0.00605) |
| **coding** | **tau_r** | **17** | 0.00507 (0.00492, 0.00523) | 0.00514 (0.00497, 0.00529) | 0.00502 (0.00486, 0.00518) | 0.00514 (0.00497, 0.00529) |
| **coding** | **tau_r** | **18** | 0.00492 (0.00474, 0.00509) | 0.00493 (0.00475, 0.00510) | 0.00489 (0.00472, 0.00506) | 0.00493 (0.00476, 0.00510) |
| **coding** | **tau_r** | **19** | 0.00546 (0.00530, 0.00562) | 0.00545 (0.00529, 0.00561) | 0.00543 (0.00526, 0.00559) | 0.00545 (0.00529, 0.00560) |
| **coding** | **tau_r** | **20** | 0.00503 (0.00488, 0.00519) | 0.00504 (0.00489, 0.00519) | 0.00499 (0.00484, 0.00514) | 0.00504 (0.00489, 0.00519) |
| **coding** | **tau_r** | **21** | 0.00467 (0.00442, 0.00493) | 0.00470 (0.00447, 0.00493) | 0.00470 (0.00447, 0.00492) | 0.00471 (0.00448, 0.00493) |
| **coding** | **tau_s** | **1** | 0.00089 (8e-04, 0.00098) | 0.00472 (0.00448, 0.00494) | 0.00170 (0.00149, 0.00191) | 0.00471 (0.00448, 0.00493) |
| **coding** | **tau_s** | **2** | 6e-05 (2e-05, 1e-04) | 0.00559 (0.00486, 0.00629) | 0.00284 (0.00227, 0.00339) | 0.00556 (0.00479, 0.00629) |
| **coding** | **tau_s** | **3** | 0.00075 (0.00057, 0.00094) | 0.00583 (0.00546, 0.00618) | 0.00194 (0.00161, 0.00228) | 0.00583 (0.00545, 0.00617) |
| **coding** | **tau_s** | **4** | 0.00107 (0.00089, 0.00125) | 0.00551 (0.00486, 0.00608) | 0.00282 (0.00232, 0.00334) | 0.00552 (0.00483, 0.00607) |
| **coding** | **tau_s** | **5** | 0.00112 (0.00095, 0.00128) | 0.00580 (0.00550, 0.00607) | 0.00209 (0.00156, 0.00263) | 0.00580 (0.00551, 0.00607) |
| **coding** | **tau_s** | **6** | 0.00142 (0.00126, 0.00158) | 0.00483 (0.00431, 0.00532) | 0.00277 (0.00252, 0.00302) | 0.00485 (0.00435, 0.00534) |
| **coding** | **tau_s** | **7** | 0.00124 (0.00111, 0.00137) | 0.00409 (0.00351, 0.00472) | 0.00218 (0.00196, 0.00240) | 0.00411 (0.00350, 0.00472) |
| **coding** | **tau_s** | **8** | 0.00144 (0.00127, 0.00160) | 0.00541 (0.00508, 0.00570) | 0.00223 (0.00160, 0.00281) | 0.00541 (0.00508, 0.00571) |
| **coding** | **tau_s** | **9** | 7e-05 (2e-05, 0.00012) | 0.00538 (0.00472, 0.00596) | 0.00305 (0.00251, 0.00355) | 0.00539 (0.00473, 0.00596) |
| **coding** | **tau_s** | **10** | 0.00178 (0.00161, 0.00195) | 0.00482 (0.00439, 0.00521) | 0.00252 (0.00226, 0.00277) | 0.00480 (0.00438, 0.00522) |

| | | | | | | |
|---|---|---|---|---|---|---|
| coding | tau_s | 11 | 0.00109 (0.00097, 0.00121) | 0.00472 (0.00433, 0.00506) | 0.00133 (0.00109, 0.00160) | 0.00474 (0.00436, 0.00507) |
| coding | tau_s | 12 | 0.00093 (0.00082, 0.00105) | 0.00518 (0.00492, 0.00540) | 0.00235 (0.00209, 0.00260) | 0.00517 (0.00492, 0.00540) |
| coding | tau_s | 13 | 0.00160 (0.00148, 0.00172) | 0.00521 (0.00499, 0.00541) | 0.00161 (0.00147, 0.00177) | 0.00521 (0.00499, 0.00541) |
| coding | tau_s | 14 | 0.00081 (0.00065, 0.00098) | 0.00628 (0.00575, 0.00671) | 0.00232 (0.00193, 0.00270) | 0.00627 (0.00571, 0.00671) |
| coding | tau_s | 15 | 0.00179 (0.00161, 0.00197) | 0.00476 (0.00418, 0.00531) | 0.00278 (0.00241, 0.00314) | 0.00476 (0.00419, 0.00532) |
| coding | tau_s | 16 | 1e-04 (3e-05, 0.00016) | 0.00473 (0.00398, 0.00545) | 0.00260 (0.00216, 0.00303) | 0.00470 (0.00394, 0.00542) |
| coding | tau_s | 17 | 0.00168 (0.00155, 0.00181) | 0.00500 (0.00473, 0.00525) | 0.00239 (0.00217, 0.00260) | 0.00500 (0.00473, 0.00525) |
| coding | tau_s | 18 | 0.00184 (0.00164, 0.00204) | 0.00416 (0.00369, 0.00462) | 0.00229 (0.00202, 0.00256) | 0.00416 (0.00367, 0.00463) |
| coding | tau_s | 19 | 0.00172 (0.00160, 0.00184) | 0.00507 (0.00471, 0.00541) | 0.00266 (0.00240, 0.00292) | 0.00507 (0.00470, 0.00541) |
| coding | tau_s | 20 | 0.00150 (0.00139, 0.00162) | 0.00476 (0.00445, 0.00504) | 0.00239 (0.00209, 0.00267) | 0.00476 (0.00445, 0.00504) |
| coding | tau_s | 21 | 0.00362 (0.00324, 0.00398) | 0.00430 (0.00392, 0.00464) | 0.00432 (0.00392, 0.00468) | 0.00436 (0.00399, 0.00472) |
| coding | tau_c | 1 | n/a | 0.00052 (0.00042, 0.00061) | 0.00021 (8e-05, 0.00033) | 0.00053 (0.00043, 0.00062) |
| coding | tau_c | 2 | n/a | 4e-05 (3e-05, 6e-05) | 2e-05 (0.00000, 4e-05) | 4e-05 (2e-05, 7e-05) |
| coding | tau_c | 3 | n/a | 0.00043 (0.00028, 0.00058) | 8e-05 (0.00000, 0.00016) | 0.00045 (0.00029, 6e-04) |
| coding | tau_c | 4 | n/a | 8e-04 (0.00058, 0.00101) | 0.00022 (4e-05, 0.00042) | 0.00083 (6e-04, 0.00106) |
| coding | tau_c | 5 | n/a | 0.00099 (0.00082, 0.00114) | 0.00044 (0.00000, 0.00077) | 0.00104 (0.00086, 0.00123) |
| coding | tau_c | 6 | n/a | 0.00042 (0.00029, 0.00055) | 6e-05 (1e-05, 0.00012) | 0.00043 (3e-04, 0.00056) |
| coding | tau_c | 7 | n/a | 4e-04 (0.00025, 0.00056) | 1e-04 (2e-05, 0.00019) | 0.00041 (0.00025, 0.00058) |
| coding | tau_c | 8 | n/a | 0.00106 (0.00088, 0.00123) | 0.00073 (0.00042, 0.00107) | 0.00110 (0.00091, 0.00128) |
| coding | tau_c | 9 | n/a | 3e-05 (2e-05, 5e-05) | 2e-05 (0.00000, 4e-05) | 4e-05 (2e-05, 7e-05) |
| coding | tau_c | 10 | n/a | 5e-05 (2e-05, 7e-05) | 1e-05 (0.00000, 3e-05) | 4e-05 (3e-05, 6e-05) |
| coding | tau_c | 11 | n/a | 0.00072 (0.00058, 0.00086) | 6e-04 (0.00036, 0.00082) | 0.00081 (0.00062, 0.00099) |
| coding | tau_c | 12 | n/a | 0.00041 (3e-04, 5e-04) | 0.00013 (4e-05, 0.00022) | 0.00041 (0.00031, 0.00051) |
| coding | tau_c | 13 | n/a | 0.00115 (0.00103, 0.00128) | 0.00147 (0.00131, 0.00160) | 0.00116 (0.00103, 0.00128) |
| coding | tau_c | 14 | n/a | 0.00058 (0.00041, 0.00074) | 8e-05 (1e-05, 0.00016) | 6e-04 (0.00042, 0.00077) |
| coding | tau_c | 15 | n/a | 0.00094 (0.00069, 0.00118) | 0.00064 (0.00039, 0.00088) | 0.00096 (0.00071, 0.00120) |
| coding | tau_c | 16 | n/a | 3e-05 (1e-05, 6e-05) | 2e-05 (0.00000, 4e-05) | 4e-05 (2e-05, 7e-05) |
| coding | tau_c | 17 | n/a | 0.00098 (8e-04, 0.00115) | 4e-04 (0.00022, 0.00057) | 0.00100 (0.00082, 0.00117) |
| coding | tau_c | 18 | n/a | 3e-05 (1e-05, 5e-05) | 1e-05 (0.00000, 3e-05) | 3e-05 (1e-05, 5e-05) |
| coding | tau_c | 19 | n/a | 0.00091 (0.00075, 0.00106) | 6e-04 (4e-04, 8e-04) | 0.00091 (0.00075, 0.00106) |
| coding | tau_c | 20 | n/a | 0.00082 (0.00069, 0.00094) | 0.00065 (0.00049, 0.00082) | 0.00082 (0.00069, 0.00095) |
| coding | tau_c | 21 | n/a | 9e-04 (6e-05, 0.00172) | 0.00092 (6e-05, 0.00174) | 0.00106 (0.00019, 0.00191) |
| coding | phi_c<-m | 1 | n/a | n/a | 0.13595 (0.07830, 0.19593) | 0.00728 (0.00000, 0.01937) |
| coding | phi_c<-m | 2 | n/a | n/a | 0.10401 (0.00518, 0.20372) | 0.00698 (0.00000, 0.02301) |
| coding | phi_c<-m | 3 | n/a | n/a | 0.09711 (0.00000, 0.19803) | 0.01299 (0.00000, 0.03940) |
| coding | phi_c<-m | 4 | n/a | n/a | 0.32695 (0.23205, 0.42542) | 0.02137 (0.00000, 0.06803) |
| coding | phi_c<-m | 5 | n/a | n/a | 0.31102 (0.15996, 0.46466) | 0.04204 (0.00000, 0.11832) |
| coding | phi_c<-m | 6 | n/a | n/a | 0.02639 (0.01151, 0.04218) | 0.00282 (0.00000, 0.00841) |
| coding | phi_c<-m | 7 | n/a | n/a | 0.07026 (0.03709, 0.10615) | 0.00466 (0.00000, 0.01384) |
| coding | phi_c<-m | 8 | n/a | n/a | 0.24908 (0.12407, 0.38234) | 0.03137 (0.00000, 0.09080) |
| coding | phi_c<-m | 9 | n/a | n/a | 0.15048 (0.06718, 0.24245) | 0.01214 (0.00000, 0.03687) |
| coding | phi_c<-m | 10 | n/a | n/a | 0.04169 (0.00785, 0.07886) | 0.00386 (0.00000, 0.01209) |
| coding | phi_c<-m | 11 | n/a | n/a | 0.13794 (0.00000, 0.27567) | 0.05520 (0.00000, 0.12933) |
| coding | phi_c<-m | 12 | n/a | n/a | 0.10137 (0.05642, 0.15020) | 0.00281 (0.00000, 0.00859) |
| coding | phi_c<-m | 13 | n/a | n/a | 0.19210 (0.00176, 0.35197) | 0.00367 (0.00000, 0.01101) |
| coding | phi_c<-m | 14 | n/a | n/a | 0.18839 (0.10709, 0.27442) | 0.02317 (0.00000, 0.06168) |
| coding | phi_c<-m | 15 | n/a | n/a | 0.15730 (0.07160, 0.24443) | 0.01095 (0.00000, 0.03343) |
| coding | phi_c<-m | 16 | n/a | n/a | 0.15124 (0.04559, 0.25918) | 0.01604 (0.00000, 0.04798) |
| coding | phi_c<-m | 17 | n/a | n/a | 0.07873 (0.03629, 0.12391) | 0.00528 (0.00000, 0.01605) |
| coding | phi_c<-m | 18 | n/a | n/a | 0.05697 (0.01679, 0.10592) | 0.00448 (0.00000, 0.01414) |
| coding | phi_c<-m | 19 | n/a | n/a | 0.11574 (0.07158, 0.16181) | 0.00370 (0.00000, 0.01106) |
| coding | phi_c<-m | 20 | n/a | n/a | 0.17410 (0.11990, 0.23178) | 0.00442 (0.00000, 0.01314) |
| coding | phi_c<-m | 21 | n/a | n/a | 0.32184 (0.03771, 0.73888) | 0.49378 (0.01240, 0.96903) |
| coding | phi_m<-c | 1 | n/a | 0.51186 (0.47796, 0.54512) | n/a | 0.50635 (0.47215, 0.54122) |
| coding | phi_m<-c | 2 | n/a | 0.44214 (0.37537, 0.50479) | n/a | 0.43340 (0.36168, 0.50215) |
| coding | phi_m<-c | 3 | n/a | 0.53862 (0.49402, 0.58446) | n/a | 0.53205 (0.48471, 0.57871) |
| coding | phi_m<-c | 4 | n/a | 0.61804 (0.54779, 0.68624) | n/a | 0.60871 (0.53344, 0.68180) |
| coding | phi_m<-c | 5 | n/a | 0.64934 (0.60737, 0.69024) | n/a | 0.63273 (0.58153, 0.68286) |
| coding | phi_m<-c | 6 | n/a | 0.31478 (0.25730, 0.37125) | n/a | 0.31434 (0.25751, 0.37057) |
| coding | phi_m<-c | 7 | n/a | 0.36471 (0.29479, 0.43596) | n/a | 0.36202 (0.29172, 0.43635) |
| coding | phi_m<-c | 8 | n/a | 0.57075 (0.52290, 0.61786) | n/a | 0.55627 (0.50236, 0.61042) |
| coding | phi_m<-c | 9 | n/a | 0.42973 (0.36364, 0.49511) | n/a | 0.42014 (0.35121, 0.48919) |
| coding | phi_m<-c | 10 | n/a | 0.35381 (0.31389, 0.39544) | n/a | 0.34922 (0.30748, 0.38952) |
| coding | phi_m<-c | 11 | n/a | 0.57335 (0.52111, 0.62507) | n/a | 0.55867 (0.49750, 0.61703) |
| coding | phi_m<-c | 12 | n/a | 0.43229 (0.39754, 0.46689) | n/a | 0.43008 (0.39587, 0.46511) |
| coding | phi_m<-c | 13 | n/a | 0.50966 (0.47918, 0.54042) | n/a | 0.50719 (0.47672, 0.53863) |
| coding | phi_m<-c | 14 | n/a | 0.60629 (0.55528, 0.65595) | n/a | 0.58939 (0.53058, 0.64735) |
| coding | phi_m<-c | 15 | n/a | 0.38112 (0.30490, 0.45615) | n/a | 0.37351 (0.29823, 0.45334) |
| coding | phi_m<-c | 16 | n/a | 0.37992 (0.30787, 0.45231) | n/a | 0.36546 (0.28624, 0.44105) |
| coding | phi_m<-c | 17 | n/a | 0.43462 (0.38634, 0.47966) | n/a | 0.43319 (0.38545, 0.47877) |
| coding | phi_m<-c | 18 | n/a | 0.34645 (0.29424, 0.39884) | n/a | 0.34238 (0.28776, 0.39390) |
| coding | phi_m<-c | 19 | n/a | 0.39945 (0.35239, 0.44644) | n/a | 0.39598 (0.34791, 0.44359) |



| | | | | | | |
|---|---|---|---|---|---|---|
| coding | phi_m<-c | 20 | n/a | 0.42318 (0.38219, 0.46390) | n/a | 0.41928 (0.37683, 0.46020) |
| coding | phi_m<-c | 21 | n/a | 0.30713 (0.03269, 0.73111) | n/a | 0.50759 (0.04163, 0.99984) |

**Table S6. Bayes factors for comparing the four models (I, O, B, 0) using coding and noncoding data on each chromosome from Heliconius**

| chr | Coding loci | | | Noncoding loci | | | Coding loci | | | Noncoding loci | | |
|---|---|---|---|---|---|---|---|---|---|---|---|---|
| | ε = 1% | ε = 0.1% | ε = 0.01% | ε = 1% | ε = 0.1% | ε = 0.01% | ε = 1% | ε = 0.1% | ε = 0.01% | ε = 1% | ε = 0.1% | ε = 0.01% |
| | B_I0 | | | | | | B_O0 | | | | | |
| 1 | ∞ | ∞ | ∞ | ∞ | ∞ | ∞ | ∞ | ∞ | ∞ | ∞ | ∞ | ∞ |
| 2 | ∞ | ∞ | ∞ | ∞ | ∞ | ∞ | 0.7027 | 0.9524 | 1.6667 | 13.2100 | ∞ | ∞ |
| 3 | ∞ | ∞ | ∞ | ∞ | ∞ | ∞ | 0.3013 | 0.3356 | 0.2326 | ∞ | ∞ | ∞ |
| 4 | ∞ | ∞ | ∞ | ∞ | ∞ | ∞ | ∞ | ∞ | ∞ | ∞ | ∞ | ∞ |
| 5 | ∞ | ∞ | ∞ | ∞ | ∞ | ∞ | ∞ | ∞ | ∞ | ∞ | ∞ | ∞ |
| 6 | ∞ | ∞ | ∞ | ∞ | ∞ | ∞ | 1.1865 | ∞ | ∞ | ∞ | ∞ | ∞ |
| 7 | ∞ | ∞ | ∞ | ∞ | ∞ | ∞ | 500.0000 | ∞ | ∞ | ∞ | ∞ | ∞ |
| 8 | ∞ | ∞ | ∞ | ∞ | ∞ | ∞ | 5.2083 | 2.5000 | 2.0000 | ∞ | ∞ | ∞ |
| 9 | ∞ | ∞ | ∞ | ∞ | ∞ | ∞ | 769.2308 | ∞ | ∞ | ∞ | ∞ | ∞ |
| 10 | ∞ | ∞ | ∞ | ∞ | ∞ | ∞ | 0.6036 | 2.2883 | 2.5000 | ∞ | ∞ | ∞ |
| 11 | ∞ | ∞ | ∞ | ∞ | ∞ | ∞ | 0.3866 | 0.3191 | 0.2481 | ∞ | ∞ | ∞ |
| 12 | ∞ | ∞ | ∞ | ∞ | ∞ | ∞ | ∞ | ∞ | ∞ | ∞ | ∞ | ∞ |
| 13 | ∞ | ∞ | ∞ | ∞ | ∞ | ∞ | 0.5836 | 0.7722 | 0.7463 | ∞ | ∞ | ∞ |
| 14 | ∞ | ∞ | ∞ | ∞ | ∞ | ∞ | ∞ | ∞ | ∞ | ∞ | ∞ | ∞ |
| 15 | ∞ | ∞ | ∞ | ∞ | ∞ | ∞ | ∞ | ∞ | ∞ | ∞ | ∞ | ∞ |
| 16 | ∞ | ∞ | ∞ | ∞ | ∞ | ∞ | 4.7059 | 5.7143 | ∞ | ∞ | ∞ | ∞ |
| 17 | ∞ | ∞ | ∞ | ∞ | ∞ | ∞ | 149.2537 | ∞ | ∞ | ∞ | ∞ | ∞ |
| 18 | ∞ | ∞ | ∞ | ∞ | ∞ | ∞ | 2.4728 | 17.8571 | ∞ | ∞ | ∞ | ∞ |
| 19 | ∞ | ∞ | ∞ | ∞ | ∞ | ∞ | ∞ | ∞ | ∞ | ∞ | ∞ | ∞ |
| 20 | ∞ | ∞ | ∞ | ∞ | ∞ | ∞ | ∞ | ∞ | ∞ | ∞ | ∞ | ∞ |
| 21 | 227.2727 | ∞ | ∞ | ∞ | ∞ | ∞ | 250.0000 | ∞ | ∞ | ∞ | ∞ | ∞ |
| | B_BI | | | | | | B_BO | | | | | |
| 1 | 0.0136 | 0.0090 | 0.0073 | 0.0101 | 0.0025 | 0.0021 | ∞ | ∞ | ∞ | ∞ | ∞ | ∞ |
| 2 | 0.0129 | 0.0063 | 0.0054 | 0.0116 | 0.0053 | 0.0046 | ∞ | ∞ | ∞ | ∞ | ∞ | ∞ |
| 3 | 0.0185 | 0.0130 | 0.0140 | 0.0109 | 0.0046 | 0.0043 | ∞ | ∞ | ∞ | ∞ | ∞ | ∞ |
| 4 | 0.0254 | 0.0196 | 0.0202 | 0.0111 | 0.0049 | 0.0046 | ∞ | ∞ | ∞ | ∞ | ∞ | ∞ |
| 5 | 0.0410 | 0.0325 | 0.0369 | 0.0122 | 0.0067 | 0.0063 | ∞ | ∞ | ∞ | ∞ | ∞ | ∞ |
| 6 | 0.0103 | 0.0034 | 0.0028 | 0.0100 | 0.0017 | 0.0011 | ∞ | ∞ | ∞ | ∞ | ∞ | ∞ |
| 7 | 0.0113 | 0.0051 | 0.0047 | 0.0104 | 0.0036 | 0.0032 | ∞ | ∞ | ∞ | ∞ | ∞ | ∞ |
| 8 | 0.0357 | 0.0306 | 0.0284 | 0.0125 | 0.0067 | 0.0061 | ∞ | ∞ | ∞ | ∞ | ∞ | ∞ |
| 9 | 0.0177 | 0.0124 | 0.0133 | 0.0146 | 0.0093 | 0.0086 | ∞ | ∞ | ∞ | ∞ | ∞ | ∞ |
| 10 | 0.0109 | 0.0041 | 0.0033 | 0.0105 | 0.0038 | 0.0031 | ∞ | ∞ | ∞ | ∞ | ∞ | ∞ |
| 11 | 0.0406 | 0.0273 | 0.0188 | 0.0107 | 0.0041 | 0.0035 | ∞ | ∞ | ∞ | ∞ | ∞ | ∞ |
| 12 | 0.0103 | 0.0033 | 0.0028 | 0.0100 | 0.0018 | 0.0012 | ∞ | ∞ | ∞ | ∞ | ∞ | ∞ |
| 13 | 0.0107 | 0.0042 | 0.0040 | 0.0100 | 0.0019 | 0.0014 | ∞ | ∞ | ∞ | ∞ | ∞ | ∞ |
| 14 | 0.0313 | 0.0241 | 0.0200 | 0.0118 | 0.0059 | 0.0051 | ∞ | ∞ | ∞ | ∞ | ∞ | ∞ |
| 15 | 0.0164 | 0.0112 | 0.0104 | 0.0108 | 0.0046 | 0.0043 | ∞ | ∞ | ∞ | ∞ | ∞ | ∞ |
| 16 | 0.0216 | 0.0151 | 0.0132 | 0.0142 | 0.0086 | 0.0081 | ∞ | ∞ | ∞ | ∞ | ∞ | ∞ |
| 17 | 0.0117 | 0.0056 | 0.0052 | 0.0100 | 0.0021 | 0.0017 | ∞ | ∞ | ∞ | ∞ | ∞ | ∞ |
| 18 | 0.0113 | 0.0046 | 0.0041 | 0.0100 | 0.0016 | 0.0010 | ∞ | ∞ | ∞ | ∞ | ∞ | ∞ |
| 19 | 0.0107 | 0.0042 | 0.0040 | 0.0103 | 0.0039 | 0.0034 | ∞ | ∞ | ∞ | ∞ | ∞ | ∞ |
| 20 | 0.0111 | 0.0051 | 0.0046 | 0.0102 | 0.0031 | 0.0026 | ∞ | ∞ | ∞ | ∞ | ∞ | ∞ |
| 21 | 0.9251 | 1.0638 | 0.8333 | 1.9778 | 3.7453 | 4.5455 | 0.8897 | 0.8130 | 1.1111 | 1.6667 | 1.8762 | 1.4925 |
| | B_IO | | | | | | B_B0 | | | | | |
| 1 | n/a | n/a | n/a | n/a | n/a | n/a | ∞ | ∞ | ∞ | ∞ | ∞ | ∞ |
| 2 | n/a | n/a | n/a | n/a | n/a | n/a | ∞ | ∞ | ∞ | ∞ | ∞ | ∞ |
| 3 | n/a | n/a | n/a | n/a | n/a | n/a | ∞ | ∞ | ∞ | ∞ | ∞ | ∞ |
| 4 | n/a | n/a | n/a | n/a | n/a | n/a | ∞ | ∞ | ∞ | ∞ | ∞ | ∞ |
| 5 | n/a | n/a | n/a | n/a | n/a | n/a | ∞ | ∞ | ∞ | ∞ | ∞ | ∞ |
| 6 | n/a | n/a | n/a | n/a | n/a | n/a | ∞ | ∞ | ∞ | ∞ | ∞ | ∞ |
| 7 | n/a | n/a | n/a | n/a | n/a | n/a | ∞ | ∞ | ∞ | ∞ | ∞ | ∞ |
| 8 | n/a | n/a | n/a | n/a | n/a | n/a | ∞ | ∞ | ∞ | ∞ | ∞ | ∞ |
| 9 | n/a | n/a | n/a | n/a | n/a | n/a | ∞ | ∞ | ∞ | ∞ | ∞ | ∞ |
| 10 | n/a | n/a | n/a | n/a | n/a | n/a | ∞ | ∞ | ∞ | ∞ | ∞ | ∞ |
| 11 | n/a | n/a | n/a | n/a | n/a | n/a | ∞ | ∞ | ∞ | ∞ | ∞ | ∞ |
| 12 | n/a | n/a | n/a | n/a | n/a | n/a | ∞ | ∞ | ∞ | ∞ | ∞ | ∞ |
| 13 | n/a | n/a | n/a | n/a | n/a | n/a | ∞ | ∞ | ∞ | ∞ | ∞ | ∞ |
| 14 | n/a | n/a | n/a | n/a | n/a | n/a | ∞ | ∞ | ∞ | ∞ | ∞ | ∞ |
| 15 | n/a | n/a | n/a | n/a | n/a | n/a | ∞ | ∞ | ∞ | ∞ | ∞ | ∞ |
| 16 | n/a | n/a | n/a | n/a | n/a | n/a | ∞ | ∞ | ∞ | ∞ | ∞ | ∞ |
| 17 | n/a | n/a | n/a | n/a | n/a | n/a | ∞ | ∞ | ∞ | ∞ | ∞ | ∞ |
| 18 | n/a | n/a | n/a | n/a | n/a | n/a | ∞ | ∞ | ∞ | ∞ | ∞ | ∞ |
| 19 | n/a | n/a | n/a | n/a | n/a | n/a | ∞ | ∞ | ∞ | ∞ | ∞ | ∞ |
| 20 | n/a | n/a | n/a | n/a | n/a | n/a | ∞ | ∞ | ∞ | ∞ | ∞ | ∞ |
| 21 | n/a | n/a | n/a | n/a | n/a | n/a | 1.8147 | 1.8768 | 1.9444 | 3.6445 | 5.6215 | 6.0380 |
